# Should I stay or should I go? Spatiotemporal dynamics of bacterial biofilms in confined flows

**DOI:** 10.1101/2024.06.07.597893

**Authors:** Massinissa Benbelkacem, Gabriel Ramos, Fatima El Garah, Yara Abidine, Christine Roques, Yohan Davit

## Abstract

Most bacteria live in sessile biofilms that colonize the confined channels, pores and crevices of natural and engineered structures. In these environments, flow delivers nutrients necessary for growth while simultaneously generating mechanical stresses that cause detachment from surfaces. Bacteria, in turn, colonize flow passages, increasing hydraulic resistance and modifying transport properties. Although the importance of advective transport and hydrodynamic forces on bacterial populations is well established, the complex feedback mechanisms governing biofilm development in confined geometries remain poorly understood. Here, we study how couplings between flow and bacterial development control the spatiotemporal dynamics of *Pseudomonas aeruginosa* in microchannel flows. We demonstrate that nutrient availability primarily drives the longitudinal distribution of biomass along the channel, while competition between growth and flow-induced detachment controls the transverse distribution and temporal dynamics. We find that biofilms undergo successive cycles of sloughing and regrowth, causing persistent fluctuations in the hydraulic resistance and biomass that prevent the system from ever reaching a true steady state. Our results indicate that these self-sustained fluctuations are a signature effect in confined flows, originating from a pressure build-up as growing bacteria obstruct flow paths. We further show that the sloughing dynamics can be described as a jump stochastic process with gamma-distributed interevent times, analogous to other bursting events such as earthquakes or avalanches. This stochastic framework provides a quantitative approach to characterizing the inherent randomness and apparent irreproducibility of biofilm experiments, opening new avenues for predictive modeling of biofilms in confined systems.

## Introduction

Fluid flow and transport phenomena control many aspects of the life of bacteria (***Krsmanovic et al., 2021***), from the motility of cells (***Busscher and van der Mei, 2006; Rodesney et al., 2017***) to the morphology of biofilm colonies (***Wang et al., 2022***), and even ecological interactions within populations (***Battin et al., 2016***). As a testimony to the importance of flow, bacteria have evolved specific strategies adapted to their mechanical environment and the rheology of fluids around them (***Dufrêne and Persat, 2020***). Bacteria have also evolved mechanisms to detect gradients of nutrients or toxic substances and adapt their movement accordingly (***Sampedro et al., 2014***). Recent studies further suggest that bacteria have mechanosensing and rheosensing capabilities (***Dufrêne and Persat, 2020***). *Pseudomonas aeruginosa* has been found to regulate the fro operon in response to flow-modulated H2O2 concentrations (***Sanfilippo et al., 2019; G.C. Padron, 2023***). Shear also modifies the intracellular levels of cyclic-di-GMP in cells of *P. aeruginosa* attached to a surface, initiating a sessile phenotype (***Rodesney et al., 2017***).

Even in dense surface-associated colonies known as biofilms (***Costerton et al., 1995; Flemming et al., 2016***), where cells are partially isolated from the fluid by a matrix of self-produced extracellular polymeric substances (***Flemming and Wingender, 2010***), flow still plays a fundamental role (***Thomen et al., 2017; Rusconi et al., 2011***). The flow of a viscous fluid around the matrix generates forces that can induce detachment or remodel the biofilm (***Besemer et al., 2007; Stoodley et al., 1998***) – the matrix essentially behaves as a viscoelastic material (***Peterson et al., 2015***) and thus can flow in response to stress (***Gloag et al., 2020***). Furthermore, key solutes, such as oxygen or nutrients, are transported in the fluid before they can diffuse in the biofilm and be consumed. This can lead to complex couplings between advective transport by flow, diffusion in the fluid and in the biofilm, and uptake by the cells (***Taherzadeh et al., 2012; Picioreanu et al., 2000***). It has also been recently demonstrated that flow can generate spatial heterogeneities in the activation of quorum-sensing within populations, as a result of autoinducers being washed away by flow in zones at high Péclet number (***Kim et al., 2016; Emge et al., 2016; Mukherjee and Bassler, 2019***).

The vast majority of bacteria live in biofilms that colonize confined geometries (***Conrad and Poling-Skutvik, 2018***). In such systems, biofilms can occupy a large portion of the available space and thus severely restrict flow. This tends to generate a two-way coupling between biofilm growth and transport phenomena whereby flow and transport mediate the development of the biofilm, but in return the development of the biofilm also modifies flow and transport (***Rittmann, 1993; Taylor and Jaffé, 1990a; Taylor et al., 1990; Taylor and Jaffé, 1990b,c***; ***Vandevivere and Baveye, 1992b,a***; ***Stewart and Fogler, 2001; Telgmann et al., 2004; Drescher et al., 2013***). In porous media, for instance, ***Kurz et al. (2022***) showed that a *Bacillus subtilis* biofilm can clog a large part of the pore space, leaving only few preferential flow channels where a competition between shear-induced detachment and growth drives intermittency and spontaneous pressure fluctuations. Such bioclogging is an important component of systems as diverse as bioreactors in the food industry (***Verran, 2002***), biofilters for wastewater processing (***Aslam et al., 2018***), pipe flow for water distribution (***Cowle et al., 2014***), heat exchangers (***García and Trueba, 2020***), catheters used in medicine (***Bixler and Bhushan, 2012***), soil bioremediation (***Singh et al., 2006***), enhanced oil recovery (***Sen, 2008; Kryachko, 2018***) or biobarriers (***Lennox and Ashe, 2009***). Understanding the fundamentals of biofilm growth and feedback mechanisms with flow is therefore a crucial step towards developing better approaches in health and engineering.

Here, we investigate the role of couplings between nutrient transport, growth and detachment on biofilm development in microchannel flows. To do so, we developed a microfluidic setup generating a constant flow rate in a microchannel where a *P. aeruginosa* PAO1 GFP biofilm develops. This microfluidic system is further combined with timelapse microscopy, cellular microbiology and mathematical modeling to study the interactions between biofilm and flow, in particular the dynamics and spatiotemporal fluctuations.

## Results

Our experimental approach is summarized in Fig 1. We inoculated cells of *P. aeruginosa* PAO1 GFP in a microchannel and then continuously flowed a culture medium at constant flow rate to observe biofilm development, which is contained in the central part of the channel through the use of UVC irradiation (see Material and Methods). In what follows, we first study the effects of nutrient limitation on the spatial distribution of biofilm. This allows us to identify flow regimes where nutrients are in excess and biofilm development is primarily driven by the interactions with flow. We then focus on regimes without limitations to detail the role of flow-induced detachment on the temporal dynamics.

**Figure 1.**
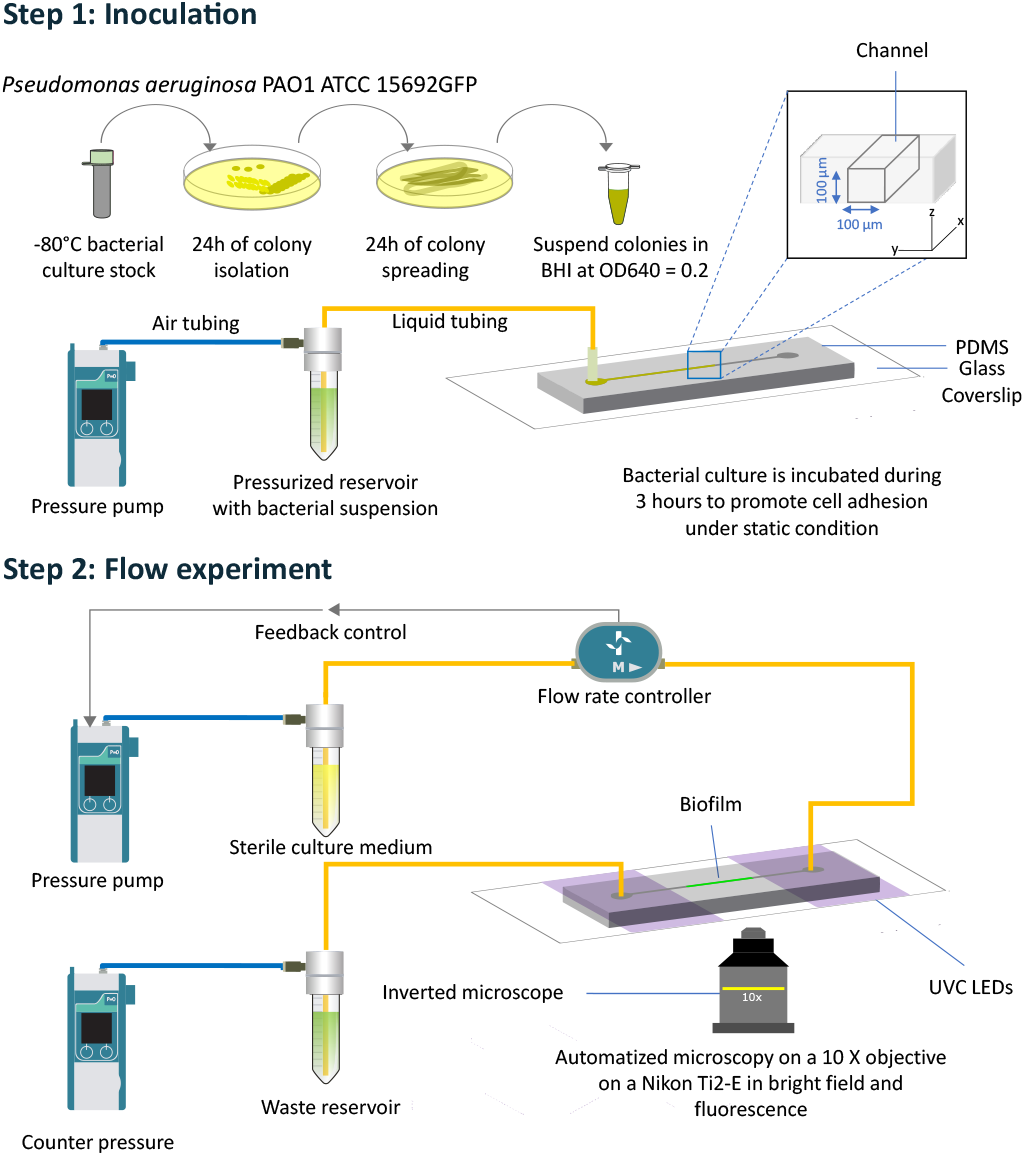
Schematic representation of the system and of the two main experimental steps. The first step is *P. aeruginosa* PAO1 GFP culture and inoculation in the microchannel (PDMS on glass, 100 *μm* × 100 *μm* cross-section) using a pressure pump. The bacterial suspension is then left for 3 hours without flow to allow cells to adhere. The second step consists in flowing the culture medium at constant flow rate through the microchannel, while recording pressure fluctuations and imaging biofilm development via optical microscopy. UVC radiation is used during the second step of the experiment to constrain the biofilm in a specific part of the channel.

### Nutrient limitation controls the longitudinal distribution of the biofilm

To assess the impact of nutrient limitation, we performed experiments at flow rates over several orders of magnitude (*Q* = 2 × 10^−2^, 2 × 10^−1^, 2 × 10^0^, and 2 × 10^+1^*μL*/*min*) and therefore different total fluxes of nutrients. Fig 2a shows the time-integrated distributions of the GFP fluorescence in the longitudinal direction for the different flow rates. We found that the active biomass expressing GFP is relatively uniform for flow rates between 0.2 and 20*μL*/*min* – the sharp decrease on the left-hand side at the outlet corresponds to the effect of the UVCs. For 0.02 *μL*/*min*, however, we observed a maximum value of the fluorescence intensity at the inlet on the right-hand side and a distinct decrease when moving towards the outlet.

**Figure 2.**
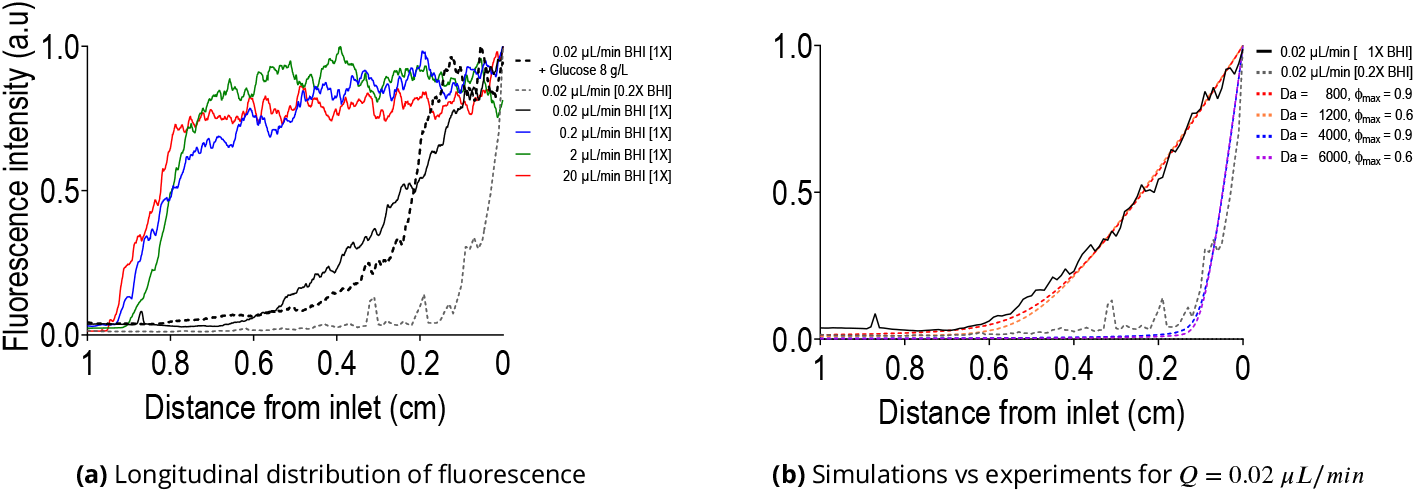
Impact of nutrient limitation on the longitudinal distribution of biofilm. (a): Time-integrated fluorescence intensity over 72h for the flow rates *Q* = 0.02 *μL*/*min*, 0.2 *μL*/*min*, 2 *μL*/*min* and 20 *μL*/*min* with 1× concentrated brain heart infusion (BHI) culture medium, along with *Q* = 0.02 *μL*/*min* with either 0.2× BHI or 1× BHI supplemented with 8 *g*/*L* glucose. Nutrient limitation is observed only for *Q* = 0.02 *μL*/*min* and is strongly dependent upon the concentration of BHI components. (b): Simulations for *Q* = 0.02 *μL*/*min* for two values of *ϕ*_max_ and the corresponding Damköhler numbers. The Damköhler numbers for the case with 0.2× BHI were simply obtained by multiplying those for the case 1× BHI by a factor 5. Each experimental curve in (a) and (b) is averaged over 3 replicates. In the model, we considered that the fluorescence signal is proportional to the product of biomass and nutrient concentration. We then integrated this signal in time and normalized it with the maximum value as 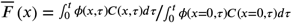.

Considering that the growth of *P. aeruginosa* PAO1 GFP is aerobic here *– P. aeruginosa* is a facultative anaerobe and can perform denitrification in specific anaerobic environments by anoxic respiration using nitrogenated compounds as final electron acceptors (***Arat et al., 2015***) but this produces less energy (***Bartberger et al., 2002***) – we hypothesized that this heterogeneity in biofilm development along the channel stems from a limitation of solute species required for growth, either oxygen or one of the nutrients in the growth medium. For the largest flow rates, advective transport through the channel should be fast compared to consumption, so that even bacteria at the outlet receive sufficient levels of nutrients and oxygen for biofilm development. For the lowest flow rate, however, one of the components introduced at the inlet is rapidly consumed by bacteria and becomes limiting, so that growth decreases with the distance from the inlet.

To better understand this limitation, we repeated experiments at 0.02 *μL*/*min* either with BHI diluted 5 times or by supplementing 1×BHI with additional glucose. Results in Fig 2a with the glucose supplementation show a similar longitudinal distribution of biofilm compared to assays without glucose, thus suggesting that this carbon source is not the limiting nutrient – it was confirmed by mass spectrometry that only a small fraction of the glucose was consumed. Results with the five times diluted BHI, however, show a much narrower window of biofilm growth, therefore suggesting that one of the components in the BHI is becoming limiting. Although oxygen probably features gradients within the biofilm (***Folsom et al., 2010***) and in the longitudinal direction, the inlet concentration of oxygen is identical for the two cases 1×BHI and 0.2×BHI, therefore indicating that oxygen is not the primary component limiting growth. Another factor that may further alleviate oxygen limitations is that PDMS is highly permeable to oxygen, so that there are in fact two sources of oxygen in our system: dissolved oxygen in the culture medium and oxygen coming from/through the PDMS.

To explore these different hypotheses and quantify the characteristic times of reactive transport, we simulated the development of the biofilm inside the channels, taking into account the couplings between biofilm growth and nutrient transport. The limiting nutrient is treated as a solute being transported by advection/diffusion and consumed by bacteria. We considered that mass transport is much faster than bacterial growth and division, so that the problem is quasi-steady for solute transport (***Picioreanu et al., 2000***). The limiting nutrient was thus modeled as

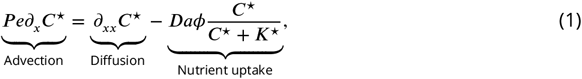

where 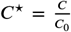 is the non-dimensionalized solute concentration with *C* [*kg* × *m*^−3^] the concentration and *C*_0_ [*kg* × *m*^−3^] the inlet concentration, *x* is the longitudinal coordinate system normalized with the length of the channel *L* = 10 *mm, K* [*kg* × *m*^−3^] the half-saturation constant and *ϕ* is the fraction of the cross section occupied by the biofilm (*ϕ* = ^Ω^biofilm/Ωtotal with Ω_biofilm_ the area of the cross section occupied by the biofilm and Ω_total_ the total area of the cross section). *P e* is the Péclet number defined as the ratio of longitudinal diffusion to advection times, 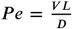 with *V* [*m* × *s*^−1^] the average velocity in the empty channel and *D* an estimate diffusion coefficient of the solute – as a reference, for a generic value *D* = 10^−9^*m*^2^ ×*s*^−1^, we have *P e* ≃ 330 for 0.02 *μL*/*min. Da* is the Damköhler number defined as the ratio of diffusive to reactive times, 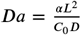 with *α* [*kg* × *m*^−3^ × *s*^−1^] the uptake rate. The term *ϕ* in the nutrient uptake indicates that consumption is proportional to the volume fraction of biofilm. Reference values are presented in Table 1.

**Table 1.** Summary of various quantities in the experiments and models. *σ*_0_ is the shear stress in empty channel. 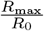 is the corresponding shear rate. *D* is an estimate diffusion coefficient for the limiting component. *P e* is the Péclet number. *ϕ*_max_ is the maximum volumic fraction of biofilm. *Da* is the Damköhler number. *τ*_reac_ is the reaction time. *P e*/*Da* is the ratio of Péclet to Damköhler numbers. To calculate dimensionless numbers, we used *D* = 10^−9^ *m*^2^*s*^−1^ and *L* = 10 *mm*.

| Conditions | Variables |  |  |  |  |  |  |
| --- | --- | --- | --- | --- | --- | --- | --- |
| | $\sigma_0$ (Pa) | $\dot{\gamma}_0$ (s<sup>-1</sup>) | $Pe$ | $\phi_{\max}$ | $Da$ | $\tau_{\text{reac}}$ (s) | $Pe/Da$ |
| $Q = 0.02 \mu\text{L}/\text{min}$<br>[BHI] = 0.2X | $2.4 \times 10^{-3}$ | $2.4 \times 10^0$ | $3.3 \times 10^2$ | 0.6 | $6.0 \times 10^3$ | 17 | $8.3 \times 10^{-2}$ |
| | | | | 0.9 | $4.0 \times 10^3$ | 25 | $5.5 \times 10^{-2}$ |
| $Q = 0.02 \mu\text{L}/\text{min}$<br>[BHI] = 1X | | | | 0.6 | $1.2 \times 10^3$ | 83 | $4.2 \times 10^{-1}$ |
| | | | | 0.9 | $8.0 \times 10^2$ | 125 | $2.8 \times 10^{-1}$ |
| $Q = 0.2 \mu\text{L}/\text{min}$<br>[BHI] = 1X | $2.4 \times 10^{-2}$ | $2.4 \times 10^1$ | $3.3 \times 10^3$ | 0.989 | $10^3$ | $\sim 100$ | $3.3 \times 10^0$ |
| $Q = 2 \mu\text{L}/\text{min}$<br>[BHI] = 1X | $2.4 \times 10^{-1}$ | $2.4 \times 10^2$ | $3.3 \times 10^4$ | 0.968 | | | $3.3 \times 10^1$ |
| $Q = 20 \mu\text{L}/\text{min}$<br>[BHI] = 1X | $2.4 \times 10^0$ | $2.4 \times 10^3$ | $3.3 \times 10^5$ | 0.907 | | | $3.3 \times 10^2$ |
| $Q = 200 \mu\text{L}/\text{min}$<br>[BHI] = 1X | $2.4 \times 10^1$ | $2.4 \times 10^4$ | $3.3 \times 10^6$ | 0.786 | | | $3.3 \times 10^3$ |

**Table 2.**
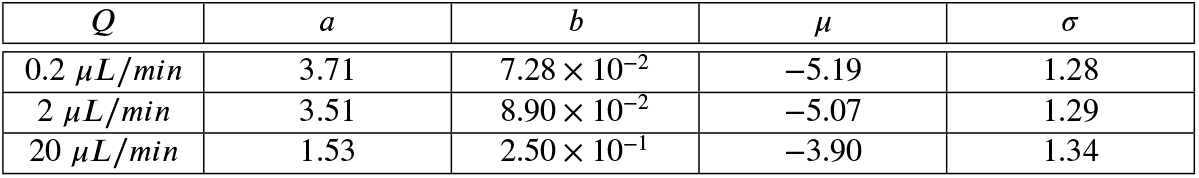
Values obtained from direct fitting of the experimental data.

| $Q$ | $a$ | $b$ | $\mu$ | $\sigma$ |
| --- | --- | --- | --- | --- |
| 0.2 $\mu\text{L}/\text{min}$ | 3.71 | $7.28 \times 10^{-2}$ | -5.19 | 1.28 |
| 2 $\mu\text{L}/\text{min}$ | 3.51 | $8.90 \times 10^{-2}$ | -5.07 | 1.29 |
| 20 $\mu\text{L}/\text{min}$ | 1.53 | $2.50 \times 10^{-1}$ | -3.90 | 1.34 |

For the biomass growth, we used a model capturing the coupled effects of growth and flow-induced removal,

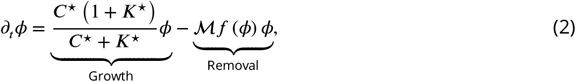

where 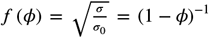 with 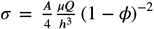 the average tangential stress at the biofilm solid interface (see SI) and 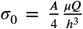 the shear stress in the empty square cross-section channel (see Table 1 for reference values). Here *A* is a scalar parameter characterizing the geometry of the cross-section colonization – 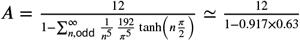 (***Bruus, 2008***) for a uniform layer of biofilm with a square cross-section for flow passage. We also have 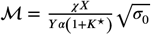, which is the dimensionless number characterizing the competition between growth and detachment, containing the rate of biomass detachment 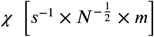.

A number of works express the detachment rate as proportional to the square root of the fluid shear stress at the biofilm fluid interface (***Coyte et al., 2017***). Here this could be written as 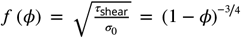 with *τ*_shear_ the shear stress at the biofilm fluid interface. Such approaches, however, were initially developed for flows in reactor systems, where shear stress is dominant (***Duddu et al., 2009; Rittman, 1982***), and neglect the contribution of the pressure stress to detachment:***Characklis et al. (1982***), for instance, measured the rate of biofilm loss considering different shear stresses generated by rotating the inner annulus of reactor at different speeds and found a linear relationship between biofilm loss rate and rotational speed. Upon clogging the microchannel, we expect the pressure difference to generate a significant force on the biofilm and to play an important role on detachment. We therefore consider a contribution of detachment scaling as (1 − *ϕ*)^−1^.

Assuming that ℳ∈ ]0, 1], the model for the biomass can be written in a simpler form as

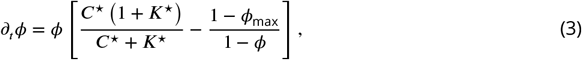

with *ϕ*_max_ the maximum volume fraction of biofilm, defined as ℳ= 1 − *ϕ*_max_, *ϕ*_max_ ∈ [0, 1[ and *ϕ* ∈ [0, *ϕ*_max_]. The only non-trivial steady state,

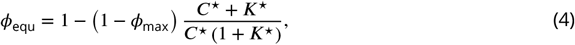

reflects an equilibrium between the amount of biomass created and the amount of biomass detached. The trivial steady state *ϕ*_equ_ = 0 is stable when detachment exceeds growth, i.e. when *C*^⋆^ is lower than 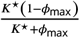. In assuming that ℳ ∈]0, 1] and eliminating cases where ℳ, > 1 we have not considered situations of systematic detachment *ϕ*_equ_ = 0 for any value of the concentration, since this is not a situation that we encountered experimentally.

Fig 2b compares results of the model with those of the experimental fluorescence for the flow rate 0.02 *μL*/*min* and *K*^⋆^ = 0.1. To roughly assess the sensitivity of our simulation to uncertainties in the value of *ϕ*_max_, we further considered two extreme values, *ϕ*_max_ ≃ 0.6 and *ϕ*_max_ ≃ 0.9. The model shows good agreement with experiments for *Da*_BHI×1_ ≃ 1200 in the case *ϕ*_max_ ≃ 0.6 and *Da*_BHI×1_ ≃ 800 in the case *ϕ*_max_ ≃ 0.9. The corresponding characteristic reaction time for nutrient uptake, *τ*_reac_ = ^*C*^_0_/*α*, ranges from ≃ 83*s* to ≃ 125*s*. We further obtain an excellent correspondence between the model and the experimental data for the 5 times dilution of the BHI without any fitting, simply by multiplying the Damköhler number by a factor 5 – recall that, by definition, *Da* = ^*αL*2^/*C*_0_ *D* with *C*_0_ the inlet concentration so that dividing *C*_0_ by 5 implies multiplying the Damköhler number by a factor 5. The fact that the behavior of the system with 5 times dilution of the BHI can be obtained without any fitting confirms that the primary limitation is indeed one of the components of the BHI, not oxygen.

With these estimations of characteristic times for uptake, we can also evaluate whether any form of transverse limitation of nutrient is expected by comparing times for transverse diffusion and uptake. The characteristic time for diffusion in the transverse direction is 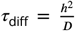 ≃ 50 *μm* with *h* a characteristic length for the distance between the flow channel and the wall. Considering again a diffusion coefficient in the biofilm *D* = 10^−9^*m*^2^/*s* for the limiting nutrient, we have 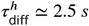. Diffusive transport across the channel is therefore significantly faster than uptake, implying that there is no form of limitation in the transverse direction.

The complete picture for nutrient transport is therefore as follows. In the longitudinal direction, the distribution of biomass is driven by a competition between advective transport and nutrient uptake. This can be quantified using the ratio of the Péclet and Damköhler numbers, ^*P e*^/*Da*. When advective transport is faster than uptake, ^*P e*^/*Da* > 1, the distribution of the biofilm is uniform. On the other hand, when ^*P e*^/*Da* < 1, uptake is faster than advective transport so that growth is nutrient limited. Table 1 summarizes the values of ^*P e*^/*Da* for the different cases and shows that the regime ^*P e*^/*Da* < 1 is reached only for the flow rate 0.02 *μL*/*min*. In the transverse direction, there is no nutrient limitation and the amount of biomass is determined by an equilibrium between growth and flow-induced detachment. The details of this mechanisms, and its impact on the temporal dynamics, is the core of the following sections.

### Hydrodynamic stresses affect all stages of development

We focus on flow rates *Q* = 0.2, 2 and 20 *μL*/*min* where nutrient limitation is negligible, allowing us to decouple the effects of molecular transport from hydrodynamic stresses. We describe the temporal dynamics through two classes of measurements: the time evolution of the pressure and timelapse microscopy imaging. The evolution of the pressure was monitored in the inlet reservoir, while imposing a constant flow rate. From this measurement, we reconstructed the evolution in time of the hydraulic resistance of the zone of interest in the channel, *R* (*t*), where biofilm developed (see Material and Methods). This allowed us to determine the dynamics of the ratio 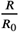, with the hydraulic resistance of the empty channel, as shown in Fig 3 (see also supplementary information figures S5, S8 and S11) and to analyze the different stages in the colonization of the channel.

**Figure 3.**
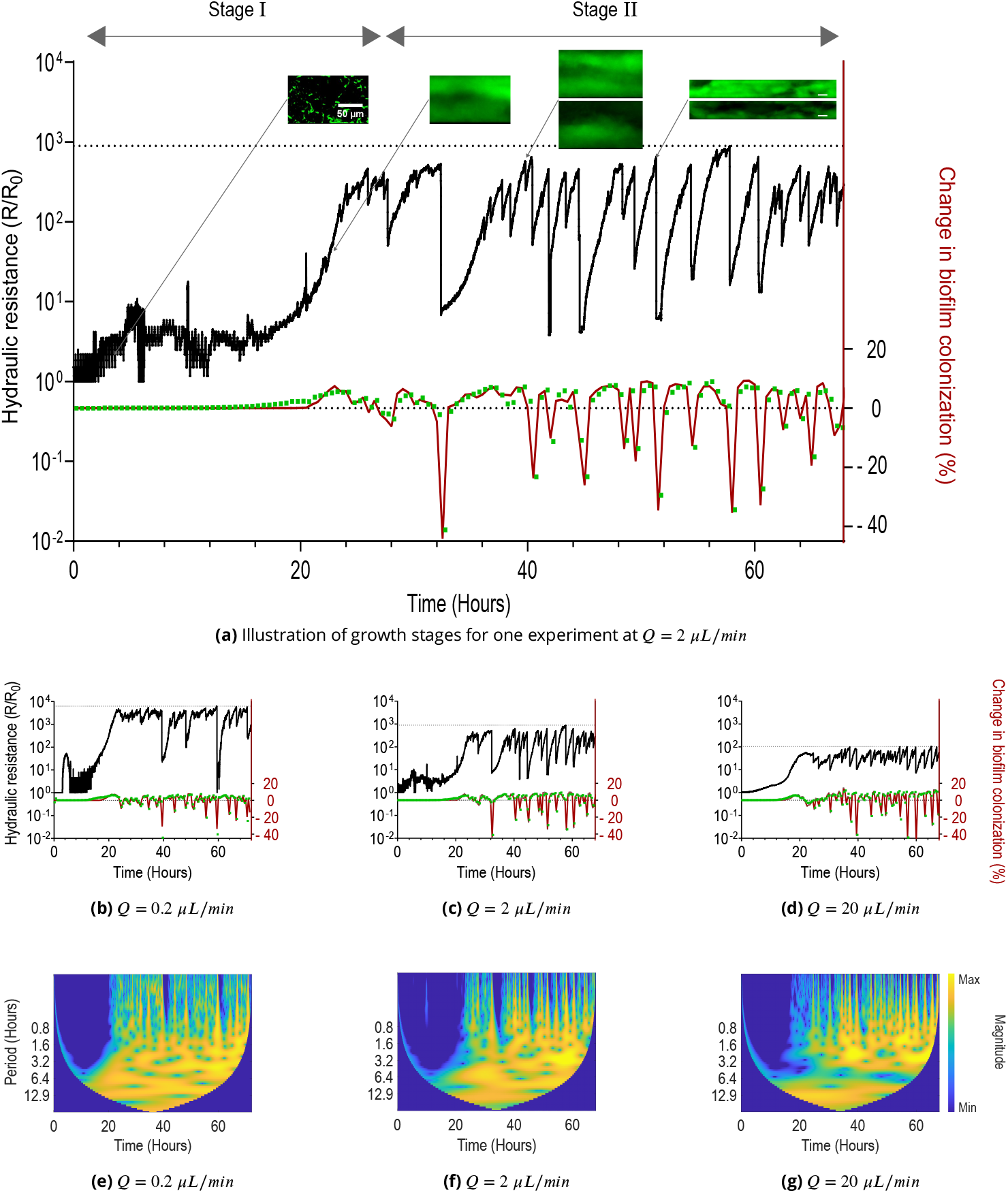
Temporal dynamics of growth and detachment for *Q* = 0.2 *μL*/*min*, 2 *μL*/*min* and 20 *μL*/*min*. (a): Summary of the two main stages of biofilm development. The figure shows the evolution of the hydraulic resistance (black solid line) in time, calculated from pressure measurements, and microscopy images corresponding to the different phases. It also shows changes in biofilm colonization extracted from either integrated fluorescence intensity (green squares) or from image segmentation (red solid line). (b), (c) and (d): Temporal dynamics of growth and detachment for the different flow rates. (e), (f) and (g): Wavelet scalograms of hydraulic resistance time series, corresponding to (b), (c) and (d). Blue stripes at short periods (near 0 hours) indicate sharp, rapid changes corresponding to sloughing events. The curved envelope at the bottom represents the cone of influence where edge effects occur.

Combined with a model for the distribution of biofilm in the cross-section of the channel – in which the biofilm was treated as as a uniform layer – it was used to indirectly evaluate a mean volume fraction of biofilm, *ϕ*. We further visualized directly the channel using timelapse microscopy with differential interference contrast, bright field and fluorescence imaging. We validated our approach by comparing *ϕ* calculated from hydraulic resistance with *ϕ* estimated from fluorescence measurements. Strong correlation between these independent methods (mean r = 0.68 - 0.77 per condition, Fig. S21) confirms that pressure-based measurements accurately capture biofilm dynamics and that the uniform layer assumption is reasonable.

We found that biofilm development occurs in different phases, in a way that is consistent across flow rates. We propose to decompose the dynamics in two main components: Stage I that corresponds to the initial adhesion, growth and saturation; and stage II that features large fluctuations in the amount of biomass in the channel. Fig 3a illustrates our decomposition in two stages on the temporal evolution of 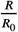 in the case *Q* = 0.2 *μL*/*min*. The following sections detail the dynamics of Stages I and II.

#### Early stage I is driven by surface adhesion, motility, division and colony formation

Our inoculation process results in the sparse attachment of individual bacterial cells on the boundaries of the channel. On the glass slide, we evaluated the initial density to be about 38800 cells per *mm*^2^. Growth of adhered cells started straight away upon flowing the culture medium, with little to no lag time. The apparent macroscopic lag, as visible in Fig 3, does not stem from a lag at the cellular level, but rather from the sensitivity of the pressure measurements. The relative hydraulic resistance, 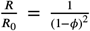, can be linearized as 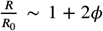 when *ϕ* ≪ 1. The evolution of 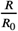 with *ϕ* is thus affine at the beginning of the experiment, with small changes in the hydraulic conductivity that could not be detected in our experimental system. This generates some amount of noise in the early signal, that tends to decrease when increasing flow rate.

To characterize the growth time at the cellular level, we calculated an apparent doubling time at the cellular level from microscopy images of cells attached to the glass slide in the very early stage (from 0 to 3.5 hours, see Fig 4e). We first segmented images from differential interference contrast microscopy to identify individual bacteria on the surface and then fitted linearly the log of the number of cells as a function of time. Calculated doubling times were measured as, on average, about 198 minutes (standard deviation ~ 14 minutes), 96 minutes (standard deviation ~ 8 minutes) and 117 minutes (standard deviation ~ 8 minutes) for, respectively, *Q* = 0.2 *μL*/*min*, 2 *μL*/*min* and 20 *μL*/*min*. As a reference, the doubling time in liquid culture was measured as about 110 minutes (standard deviation of ~ 10 minutes). Although understanding exactly what generates this dependence of the doubling time upon the flow rate is beyond the scope of this paper, it is worth noting that nutrient limitation is not likely to play a role, as we have previously validated the fact that there is no limitation for *Q* ≥ 0.2 *μL*/*min*, even when biofilm has formed. Mechanisms directly related to flow could mediate these phenomena (***Sanfilippo et al., 2019; G.C. Padron, 2023; Rodesney et al., 2017***), as well as heterogeneities in the rate of division – for instance between motile and adhered cells (***Rossy et al., 2019***) – or in the attachment/detachment dynamics. After this initial phase, colonies expand into mature biofilms, yielding a much sharper increase in hydraulic resistance. This increase stems from the combination of bacterial growth and the nonlinear relationship between between ^*R*^/*R*_0_ and *ϕ*, ^*R*^/*R*_0_ = (1 − *ϕ*)^−2^.

**Figure 4.**
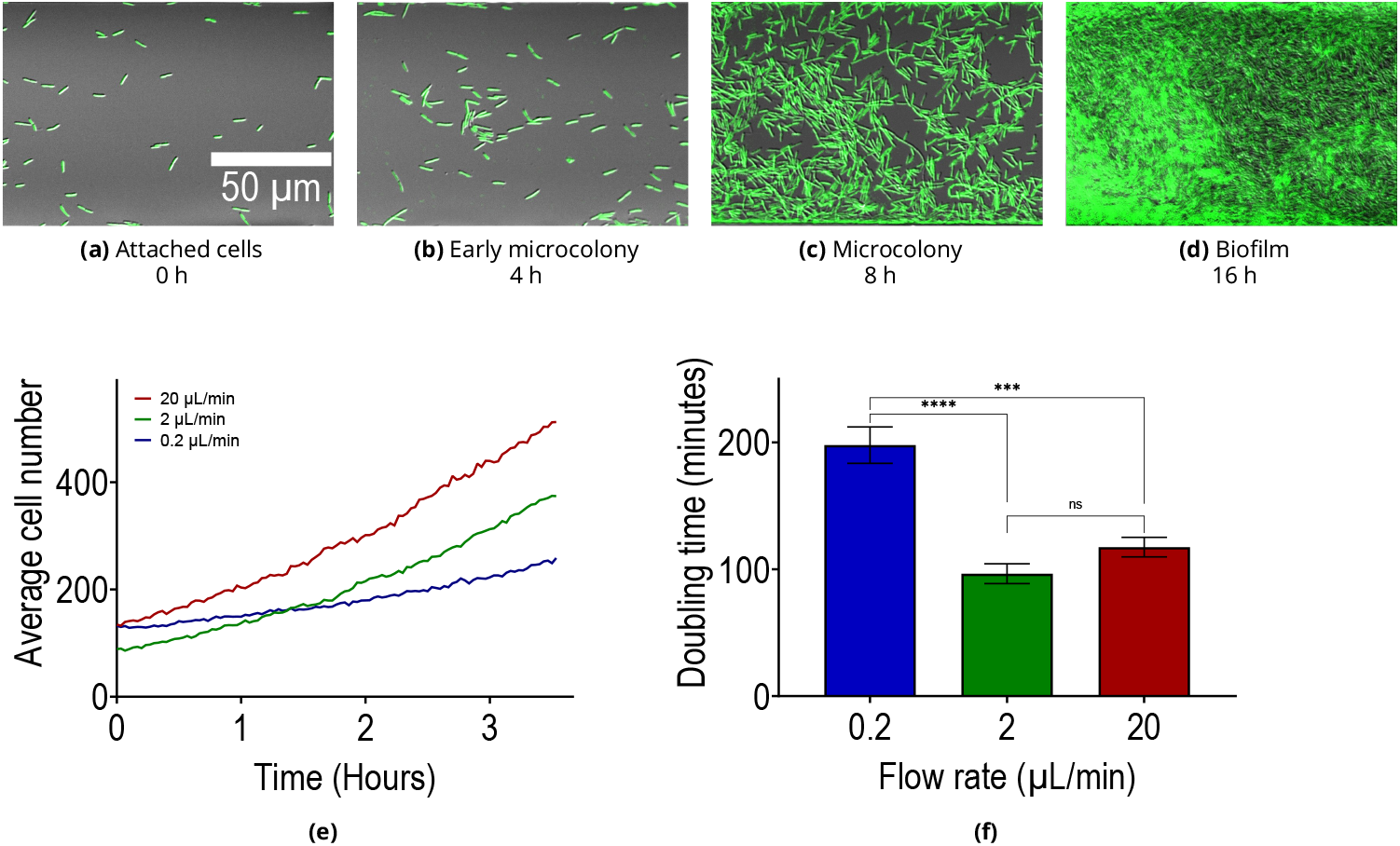
Flow modifies the apparent doubling time of bacteria on surfaces. (a), (b), (c) and (d): Composite brightfield and GFP images of development stages, starting from single cells that form microcolonies and then evolve towards a biofilm. (e) and (f) show, respectively, the average number of cells on the surface as a function of time and the corresponding doubling time for flow rates (*Q* = 0.2 *μL*/*min*, 2 *μL*/*min* and 20 *μL*/*min*). For (f), statistical differences were examined by unpaired Student’s t-test with Gaussian distribution of data and equal standard deviations. Error bars indicate standard error of mean (SEM) and symbols denote statistical significance (****: p<0.0001, ***: p<0.001, ns: p > 0.05). The doubling time was calculated by a linear fitting of the logarithm of the number of cells. The slope was used to estimate growth rate and doubling time. Cell count was calculated from image segmentation of four positions in two channels to generate 8 measurements by condition (*n= 8*) for (*Q* = 0.2 *μL*/*min*, 2 *μL*/*min* and 20 *μL*/*min*).

#### Late Stage I reflects an equilibrium between growth and detachment

We observe an inflection of the signal in late Stage I in Fig 3a, 3b, 3c, 3d. This inflection corresponds to the end of the rapid growth phase, with flow-induced detachment progressively increasing, until detachment equilibrates with growth. To further understand this effect, let us consider the mass balance of biofilm in the zone of interest – the zone where biofilm grows in between the two UVC irradiation zones – in the channel. The inlet flux of bacteria is zero, since our UVC system prevents growth outside the zone of interest. The source of biomass therefore only results from the uptake of nutrients, which yields division of bacterial cells and EPS production. The sink of biomass is due to flow-induced removal, which includes parts of the biofilm that are displaced out of the zone of interest through the flow/remodelling of the biofilm at the outlet, and parts that are washed away by erosion and seeding dispersal (***Kaplan, 2010; Krsmanovic et al., 2021***). This model captures a competition between growth and different types of smooth detachment – continuous, non-catastrophic biomass loss (as opposed to sudden sloughing events) – which occurs through three mechanisms: (1) biofilm flow, where viscoelastic deformation causes biofilm to move out of the observation zone; (2) erosion, the continuous release of single cells or small clusters from the biofilm surface; and (3) seeding, the release of cells from within the biofilm. In the absence of nutrient limitation, we can estimate the evolution of the biovolume in the channel from the ordinary differential equation

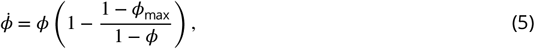

which is a direct simplification of Eq 3 with *ϕ* describing an average volume fraction in the entire channel, and thus being only a function of time. This model captures a competition between growth and different types of smooth detachment (flow, seeding, erosion).

#### Stage II features sloughing-induced jump events followed by re-growth

After growth and detachment start to equilibrate at the end of Stage I, sloughing events become particularly important. These are visible in Fig 3a, 3b, 3c, 3d as jumps in the hydraulic resistance. Wavelet scalogram analysis of the hydraulic resistance time series (Fig 3e, 3f, 3g) reveals the temporal dynamics of biofilm detachment throughout the growth cycle. The blue stripes at short periods (near 0 hours) indicate sharp, rapid changes in hydraulic resistance corresponding to sloughing events. Unlike time-averaged or Fourier methods, wavelet analysis uniquely identifies when these abrupt detachment events occur, showing that sloughing is an intermittent rather than continuous process. The jumps correspond to a range of different events, from minor detachment – where a relatively small portion of the biomass is detached – to major sloughing – where a large portion of the biomass is detached. Fig 5b shows example microscopy images of such events.

**Figure 5.**
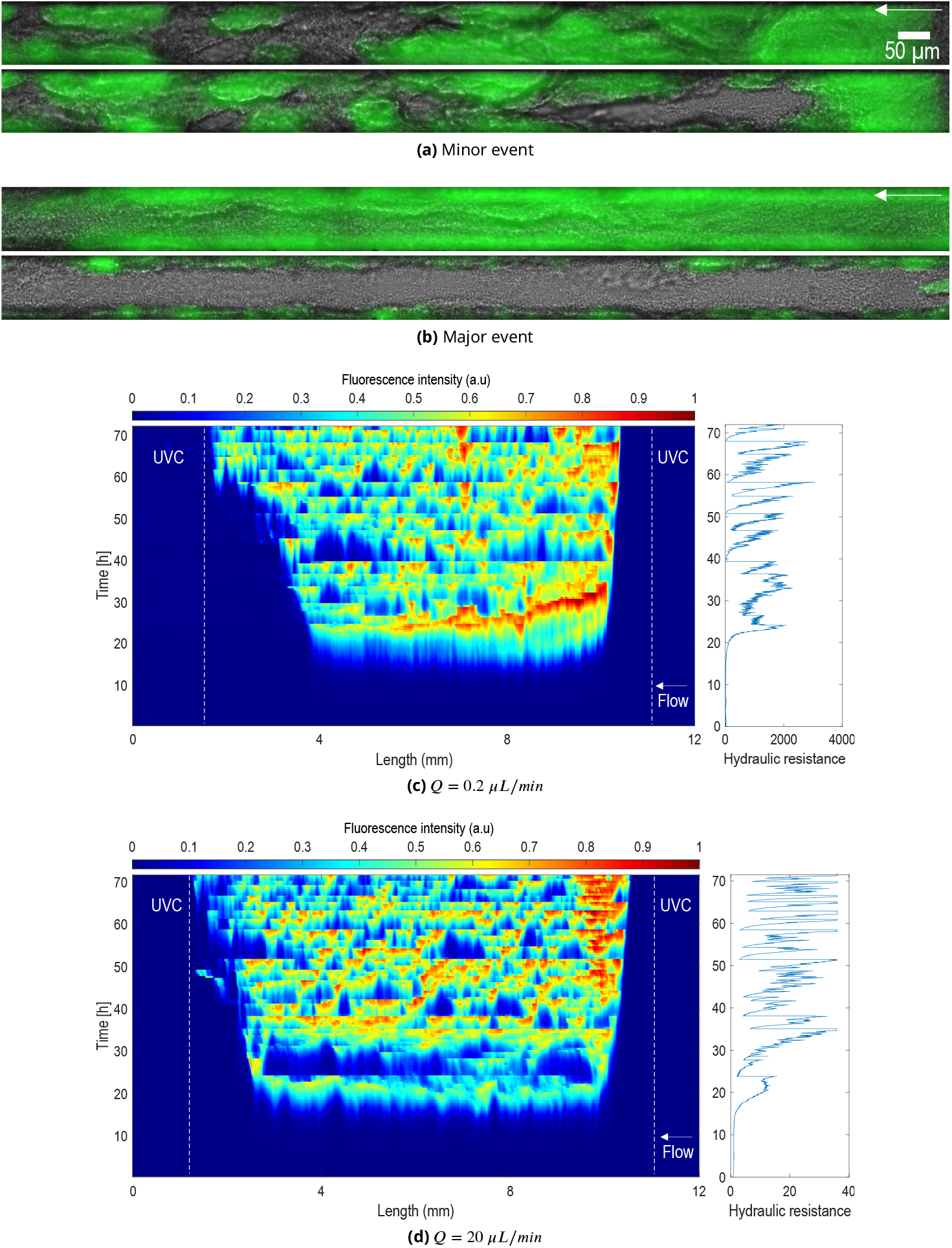
Spatiotemporal dynamics of sloughing in Stage II. (a) and (b): composite bright field and GFP image of a ≃40 hour biofilm during Stage II. Image before (top, time t) and after (bottom, time t + 30 minutes) for (a) a minor sloughing event and (b) a major sloughing event. (c) and (d): kymographs showing the fluorescence intensity (averaged in the transverse direction *y*) as a function of both the longitudinal direction (*x*) and time. Fluorescence intensity values were normalized by the maximum value. Plots on the right-hand side show the corresponding normalized hydraulic resistance *R*_max_/*R*_0_ as a function of time.

To better visualize the spatiotemporal dynamics and connect the different observations for the pressure and microscopy, we plotted kymographs in Fig 5c, 5d (see also SI figures S7, S10 and S13), showing both variations in time and in space, along with the corresponding hydraulic resistance signal. These graphics show clearly the correlation between the pressure signal on the right-hand side and the detachment events on the kymographs. We can readily identify that large drops in pressure correspond to large events with detachment over almost the entire length of the micro-channel. We also visualize a range of detachment events corresponding to various sizes of biofilm being detached, along the rapid increase of resistance and biomass following each sloughing event.

Each sloughing event is followed by a rapid growth phase resembling late Stage I, leading to a cycle of successive sloughing and re-growth. In each re-growth phase, we observe a sigmoid-like shape of the hydraulic resistance signal that is similar to Stage I. As in Stage I, the maximum stems from an equilibrium between growth and detachment. *ϕ*_max_ thus corresponds to a critical value of the stress, *σ*_crit_, that generates enough detachment to compensate growth. From Eqs 2-3, this reads

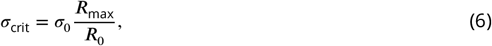

with *R*_max_ the maximum value of the hydraulic resistance. The maximum value of the hydraulic resistance over all stages of development decreases by orders of magnitudes from 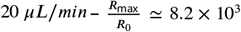 for 0.2, ≃ 8.4 × 10^2^ for 2 and ≃ 1.0 × 10^2^ for 20. We can thus estimate the critical stress at about *σ*_crit_ ≃ 100 − 200 *P a*.

#### Pressure stress becomes prominent in late Stage I and Stage II

We now ask the question of whether the hydrodynamic stress is dominated by shear or pressure. We are particularly interested in understanding the nature of *σ*_crit_. Fig 6 shows simple 2D flow simulations illustrating the types of stress induced by the flow of a viscous fluid upon the biofilm. We see that the shear stress becomes particularly strong in the bottlenecks and that the pressure difference also builds up, therefore generating both shear and pressure stresses. In the initial stages of bacterial development, when only individual cells and microcolonies are adhered to surfaces, we expect the dominant stress to be shear. However, what happens in later stages for large fractions of biofilm?

**Figure 6.**
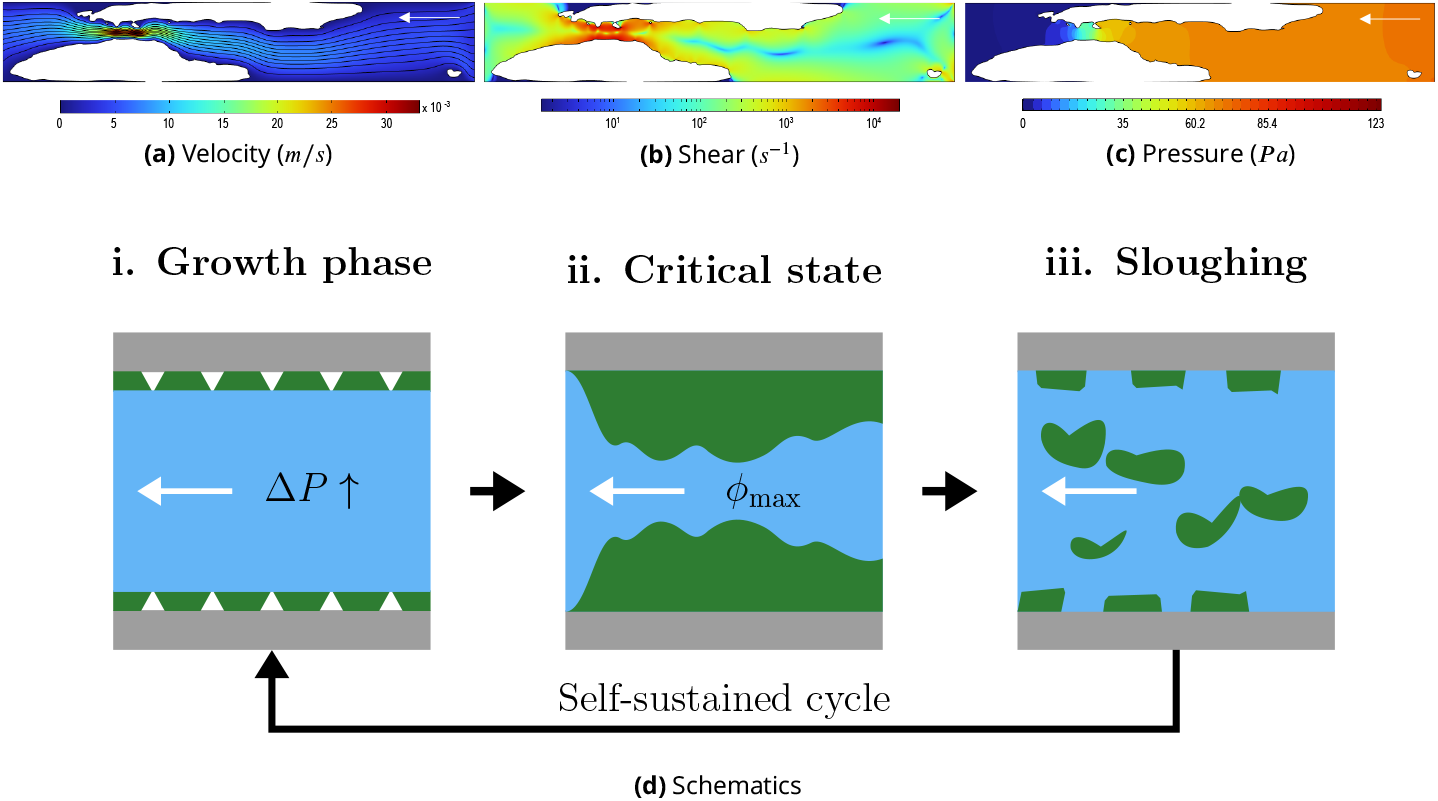
Mechanisms of pressure increase. (a), (b) and (c): 2D simulations of flow around a biofilm (white) in a channel of 100 *μm* width and 500 *μm* length using COMSOL multiphysics. (a): Streamlines and magnitude of the velocity (*m*/*s*). (b): Shear rate in (*s*^−1^). (c): Pressure field in (*P a*). (d) Schematics of growth and sloughing. The white arrows indicate the flow direction.

A simple approach to quantifying the relative importance of stresses in our system is to consider the case of uniform film growth between the UVC zones (see details in supplementary information). The tangential stress at the solid/biofilm surface is 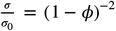, with a contribution from shear 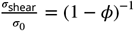 and from pressure 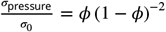. In the case of a fixed flow rate, we therefore have 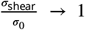 and 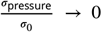 in the limit *ϕ* → 0; *σ*_shear_ = *σ*_pressure_ for *ϕ* = 0.5; and *σ*_pressure_ > *σ*_shear_ for *ϕ* > 0.5. This simple conceptualization confirms that pressure stress tends to become dominant when a large portion of the channel is colonized, so that *σ*_crit_ is primarily a pressure stress. Furthermore, sloughing events mainly develop when the volume fraction is large suggesting that these events are also driven by pressure stress.

#### Maximum clogging depends on the flow rate

In our experiments, the maximum values of the volume fraction, estimated from hydraulic resistance, are *ϕ*_max_ (*Q* = 0.2) = 0.989 for 0.2, *ϕ*_max_ (*Q* = 2) = 0.966 for 2 and *ϕ*_max_ (*Q* = 20) = 0.901 for 20 (see Fig 7a, 7b, 7c. and Table 1) – remarkably, this maximum is very reproducible across replicates, with standard deviations for volume fractions in the range of 10^−3^ in our experiments (Fig 7).

**Figure 7.**
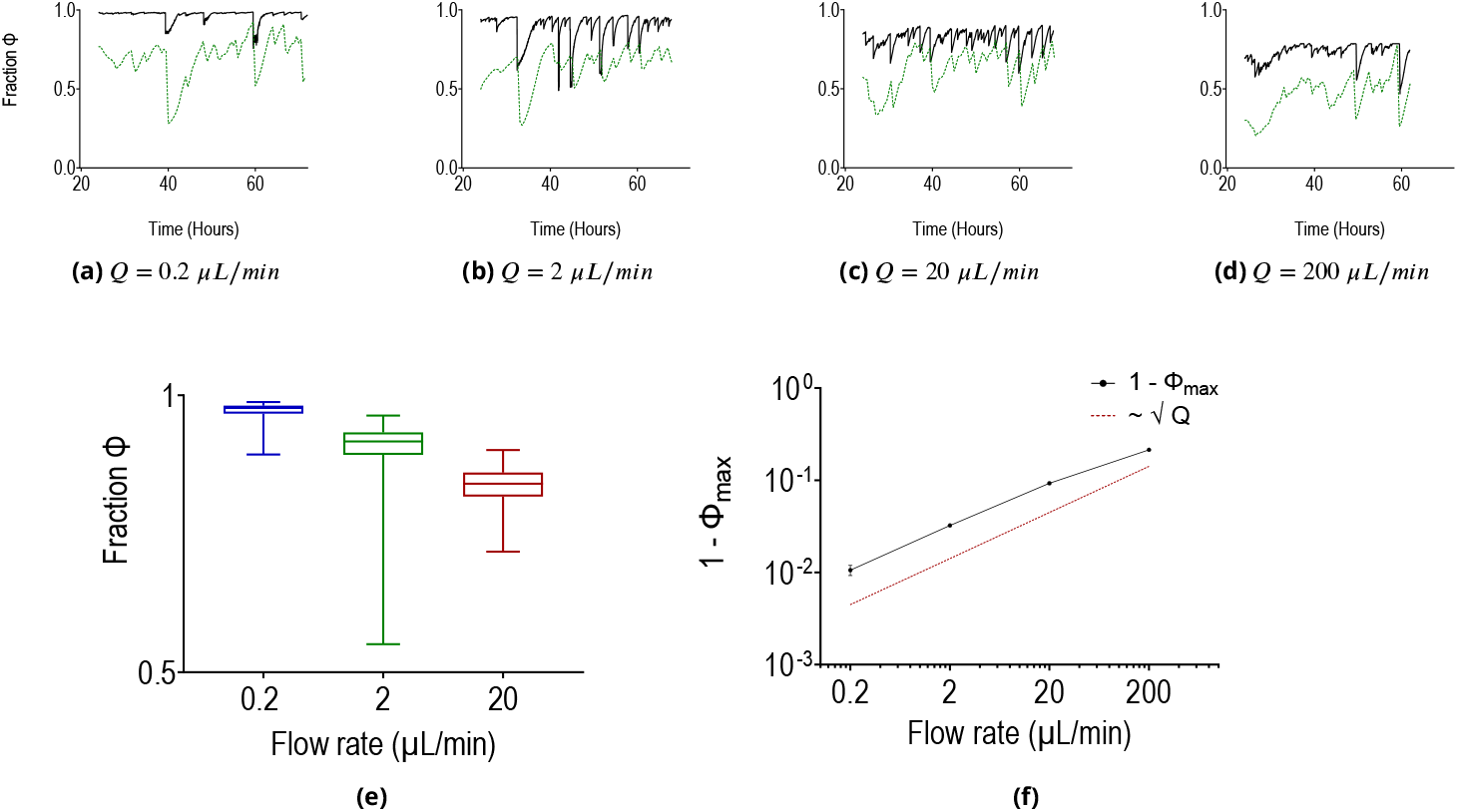
Evolution of the distribution and maximum of the volume fraction for the different flow rates. (a), (b), (c) and (d): Volume fraction in the microchannel, calculated from hydraulic resistance (black solid line) or estimated from integrated GFP intensity (green dotted line), for the different flow rates. These independent measurements are strongly correlated (Fig. S21). (e): Distribution of biofilm fraction calculated from hydraulic resistance between 24 and 72 h for all flow rates, represented as whisker boxes. (f): Log-log plot of 1 − *ϕ*_max_, with volume fraction calculated from hydraulic resistance, as a function of the flow rate (*Q* = 0.2 *μL*/*min*, 2 *μL*/*min*, 20 *μL*/*min* and 200 *μL*/*min*). The red dotted line simply shows the slope for an evolution with the square root of the flow rate.

The model Eq 5 for Stage I is consistent with this dependence upon the flow rate. By definition of *ϕ*_max_ in Eqs 2-3 and in Eq 5, we have

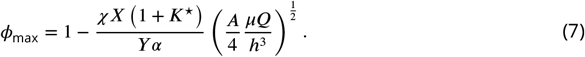

We can express this relatively to a reference flow rate as

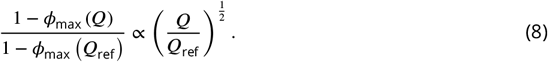

Upon considering *Q*_ref_ ≡ 0.2 with *ϕ*_max_ *Q*_ref_ = 0.989, we obtain 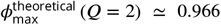 and 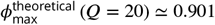 from Eq 8. These values are in excellent agreement with the experiments (see Table 1).

To further validate this idea that *ϕ*_max_ decreases with the square root of the flow rate (Eq 8), we performed a single experiment at 200 *μL*/*min*. Results in Fig 7d indicate that the behavior is similar to that of other flow rates, but with a significantly lower value of *ϕ*_max_ ≃ 0.786. We also see that the scaling remains remarkably similar with 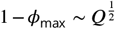 in the range [0.2, 200] – equivalently, the scaling for the hydraulic resistance is *R*_max_ ~ *Q*^−1^ (see Fig S19). The fact that the model can recover the correct scalings is an important validation of our modeling hypotheses, in particular that the detachment rate depends on square root of the hydrodynamic stress.

#### Maximum clogging depends on the composition of the biofilm

Modifications of the composition and mechanical properties of the biofilm should have a significant impact on the value of *ϕ*_max_. Among *P. aeruginosa* PAO1 polysaccharides – Pel, Psl and alginate – Pel and Psl are considered the two primary components of the EPS matrix structure (***Ryder et al., 2007***), controlling the cohesive and adhesive strength of the biofilm. We therefore performed experiments with mutants that cannot produce either Pel or Psl, with the idea that this should modify the value of *χ*. The Psl mutant is a *pslD* deficient Δ*pslD* obtained from *P. aeruginosa* PAO1 by non-polar allelic exchange (***Colvin et al., 2012***). The Pel mutant is a pelF deficient strain Δ*pelF* also obtained by allelic exchange (***Colvin et al., 2011***). Liquid culture showed that the lag and doubling times were, respectively, 233.36 and 105.5 minutes. Fig 8 (see also S15, S16, S17 and S18) shows the behavior of mutants compared to the wild type. We found that the volume fraction of biofilm was significantly lower for both mutants and that the Psl mutant was more strongly affected than the Pel mutant. The Psl mutant only weakly attached to the surface and was more prone to detachment – an observation that is consistent with the prominent role of Psl in the mechanics of PAO1 biofilms (***Colvin et al., 2011***). For the case 2 *μL*/*min*, we even observed an almost complete detachment of the biofilm with only few cells remaining attached and making re-growth possible after a sloughing event. At 20 *μL*/*min*, no effect was visible on the pressure measurements.

**Figure 8.**
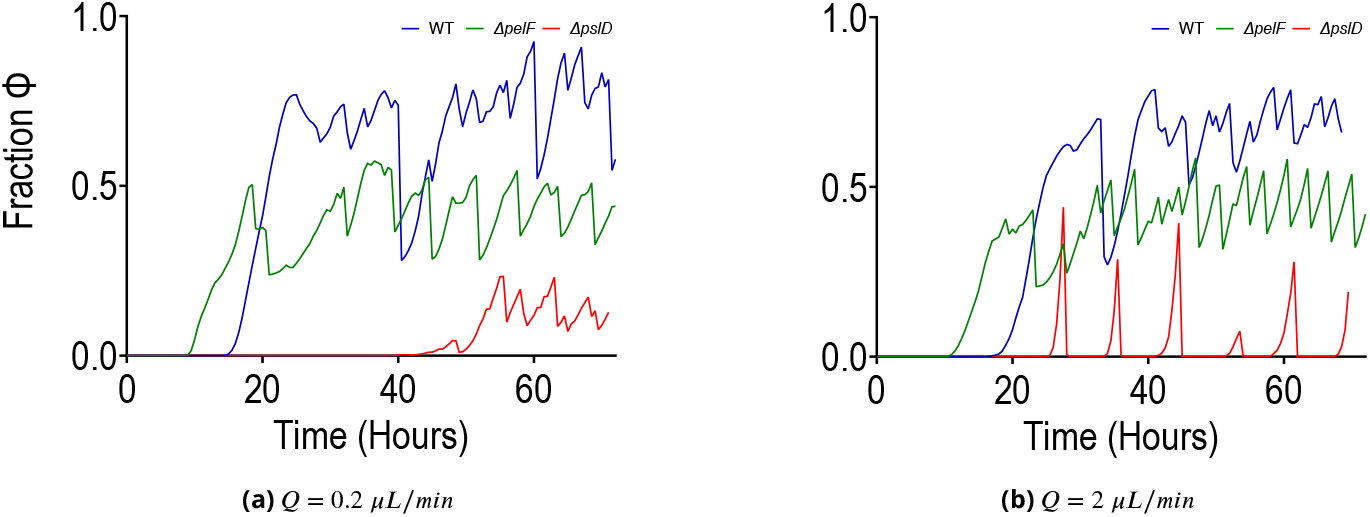
Fraction of biofilm obtained from image segmentation for the Δ*psl* (red line) and Δ*pel* strains (green line) compared to the wild-type strain (blue line), for (*Q* = 0.2 *μL*/*min*, 2 *μL*/*min*).

### Biofilm development is stochastic

The model presented so far can describe Stage I and the signal in between two sloughing events, but not the catastrophic jumps associated with sloughing. To characterize the latter, we first performed a frequency analysis of the signal, as shown in Fig 9a. This approach did not prove very informative as it essentially shows a power spectrum typical of noise, with a slope roughly smaller than −2. This result, however, motivated the construction of a more physical representation of the process. The basis of this representation is the observation that fluctuations have a very specific signature on the hydraulic resistance: they first feature a sharp decrease due to sudden sloughing, followed by a slower increase due to growth. Since growth and smooth biofilm detachment are already described in the model, this observation suggests that we only need to capture the sudden sloughing to improve the description. The model takes the form of the following stochastic differential equation,

**Figure 9.**
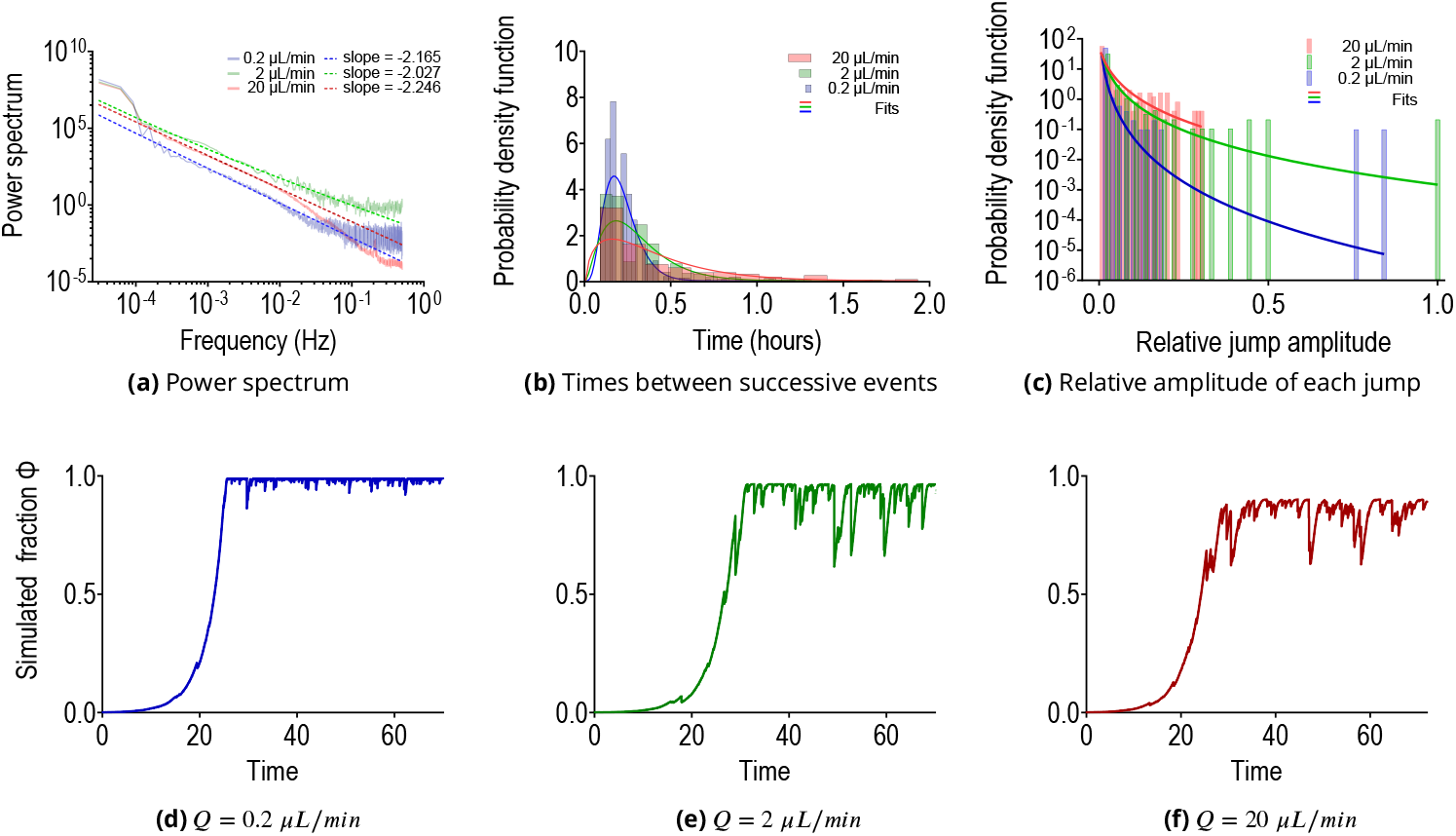
Stage II detachment events can be described as a jump process. (a) Power spectrum for the different flow rates (*Q* = 0.2 *μL*/*min*, 2 *μL*/*min* and 20 *μL*/*min*, 3 replicates for each). Slopes are indicative and were calculated in the interval between 0 and 0.25 Hz. (b) Probability density function of the time between two successive jump events, *δt*, for (*Q* = 0.2 *μL*/*min*, 2 *μL*/*min* and 20 *μL*/*min*). The histograms are calculated from data from all experiments/replicates, while the solid lines are fitted Gamma distributions (*δt*)^*a*−1^/(*b*^*a*^Γ(*a*)) exp (−*δt*/*b*). (c) Probability density function of the relative amplitude of jump events, 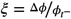 for (*Q* = 0.2 *μL*/*min*, 2 *μL*/*min* and 20 *μL*/*min*). The histograms are calculated from all experiments/replicates, while the solid lines are fitted log-normal distributions 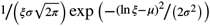.(d), (e) and (f): Stochastic simulations of the volume fraction as a function of time for (*Q* = 0.2 *μL*/*min*, 2 *μL*/*min* and 20 *μL*/*min*), which can be compared to experimental results in Fig 7.

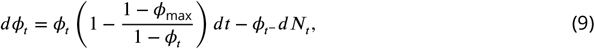

with *ϕ*_*t*_ describing the average fraction of biofilm in the microchannel, *N* the jumps and 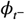 the value of *ϕ* at time *t*^−^ just before the jump.

The challenging part of this representation is to accurately describe the randomness of the process *N* – which is likely strongly dependent upon spatial heterogeneities, for instance in the initial attachment of bacteria. We characterized *N* in our experiments via the distributions of both the times between two successive jumps and the relative amplitudes of the jumps – i.e. the amplitude of each jump divided by the value of the volume fraction just before the jump 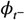. Fig 9b, 9c shows histograms of these distributions extracted from signal processing of the experimental data (see Material and Methods). We see that the distribution of times between successive jumps, *δt*, can be well approximated by a Gamma distribution – solid lines in Fig 9b for a fit of the experimental data (see Supplementary figure S20 for quantile-quantile plots). The relative amplitude of each jump 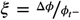 was better represented by log-normal distributions – solid lines in Fig 9c for a fit of the experimental data. Example realizations of the simulations are presented in Fig 9d, 9e, 9f for the flow rates used in the experiments. Simulations are strikingly similar to experimental data, thus suggesting that sloughing can indeed be accurately modeled as a stochastic jump process.

## Discussion

### Regimes of nutrients and oxygen transport

Our experimental setup, which implements a novel UVC strategy to confine biofilm development within a specific zone in the channels, allowed us to finely control boundary conditions and study the role of molecular transport on biofilm development. Our results suggest that, in confined channels, the longitudinal distribution of biofilm ensues from a competition between advective transport of solutes along the channel and uptake by the cells. Modeling molecular transport as an advection-diffusion-reaction equation, this effect can be quantified using the ratio of longitudinal Péclet and Damköhler numbers, ^*Pe*^/*Da*, which compares advection to reaction. When ^*Pe*^/*Da* < 1, biofilm growth is nutrient-limited and localized close to the inlet. When ^*Pe*^/*Da* > 1, biofilm development is close to homogeneous. In the transverse direction, the limitation can be characterized in a similar way via a transverse Damköhler number, *Da*_transverse_, defined as the ratio of reaction and transverse diffusion. In our experiments, we always had *Da*_transverse_ < 1 with only weak gradients and no transverse limitation.

Although the competition between advective/diffusive transport and uptake is a universal feature of biofilm development in flow, the solute that becomes limiting depends on the details of the setup. In our experiments, we found that nutrients were restricting growth, while other works in microchannel flows, such as (***Thomen et al., 2017***), found that the effect of oxygen was dominant. Several factors play an important role here, including the composition of the growth medium, the catabolism of the microorganisms, the kinetics or even the resilience of bacteria to low oxygen concentration. For instance, ***Thomen et al. (2017***) focused on an *E. coli* K12 derivative. *E. coli* can respond strongly to micro-aerobiosis (***Colón-González et al., 2004***), whereas *P. aeruginosa* is known to be more resilient (***Sabra et al., 2002; Alvarez-Ortega and Harwood, 2007***). Factors that affect transport characteristic times, such as the flow velocity or the dimensions of the channels, also modify the competition with uptake. The length of the channel modulates the time for advective transport along the channel and therefore the ratio ^*Pe*^/*Da*. The width and height control the characteristic time for transverse diffusion. In ***Thomen et al. (2017***), for instance, the width of the channel is 1000 *μm* so that diffusion takes about 100 times longer and *Da*_transverse_ is 100 times larger than for 100 *μm* channels.

### Temporal dynamics and hydrodynamic stresses

With these regimes clearly identified for molecular transport, we could then study the effect of hydrodynamic stresses on biofilm development and, in particular, the role of flow-induced detachment on the spatiotemporal dynamics. We suppressed any form of nutrient limitation, either transverse or longitudinal, by focusing on flow rates ≥ 0.2 *μL*/*min* with ^*Pe*^/*Da* > 1 and *Da*_transverse_ < 1. In so doing, we found that the time response featured two distinct stages: the hydraulic resistance first followed a relatively standard sigmoid-like pattern (Stage I), but then switched to a different regime with large self-sustained fluctuations (Stage II). As discussed in what follows, we argue that this Stage II is a signature effect of biofilm development in confined flow.

Stage I began just after initial inoculation, as soon as the culture medium started flowing in the microchannel. Early Stage I was dominated by cell adhesion, division, surface motility and microcolony formation. The average velocity in the channel at 0.2 *μL*/*min* is above 300 *μm*/*s* so bulk motility is likely not playing an important role when *Q* ≥ 0.2 *μL*/*min* (***Rossy et al., 2019***). Contrary to ***Thomen et al. (2017***) – where no direct initiation of the biofilm of *E. coli* biofilm was possible above a shear stress of 10 *mP a* – our results show that *P. aeruginosa* PAO1 strongly adhered to the surface and could initiate biofilm formation for all shear stresses. We even observed that the apparent doubling time within the first few hours of growth was the smallest in the case at 2 *Pa*. Stage I then evolved towards the standard steps of biofilm formation for *P. aeruginosa* (***Rasamiravaka et al., 2015; Krsmanovic et al., 2021***), with clonal microcolonies that extended, joined together and grew into mature biofilms. The exponential growth and EPS production yielded a sharp increase of the hydraulic resistance by several orders of magnitude, which is consistent with previous works (***Cunningham et al., 1991; Desmond et al., 2022***).

Stage II was characterized by large fluctuations in the hydraulic resistance and in the quantity of biomass in the channel. We showed that these fluctuations are the result of successive cycles of growth and sloughing events: parts of the biofilm get detached by the flow upon reaching a critical stress, generating sharp discrete-like events of detachment, followed by a fast re-growth of the biofilm. In both late Stage I and in Stage II, pressure-induced stresses play a crucial role. Most works have focused on shear stress (***Krsmanovic et al., 2021; Kurz et al., 2022; Telgmann et al., 2004; Wagner et al., 2010; Paul et al., 2012***) because, in many situations, such as biofilm development in large reactors, the effect of pressure can be neglected. However, this is not the case in confined systems, where biofilm can clog a large proportion of the flow channels and the pressure build-up induced by clogging can generate an important force on the biofilm.

### Maximum clogging

One aspect that stands out in our experiments, because of its physical meaning and reproducibility across replicates, is the concept of maximum clogging. In Stage I and after each sloughing in Stage II, curves for the hydraulic resistance had a sigmoid-like shape, with an inflection that was a signature of a competition between growth and smooth detachment due to erosion, seeding and biofilm flow outside the zone of interest. We found that the maximum value of the hydraulic resistance across the entire experiment scales with the inverse of the flow rate, which is consistent with our modeling hypothesis that smooth detachment scales with the square root of the tangential stress at the solid/biofilm interface (which includes pressure contributions), rather than with shear stress at the biofilm/fluid interface as in previous reactor studies. This concept of maximum clogging can also be expressed as a critical value of the hydrodynamic stress generating enough detachment to compensate growth. In our framework, this equilibrium stress was in the order of 100-200 Pascals for the wild type and did not vary significantly with the flow rate.

To further understand the role of the EPS matrix composition on maximum clogging, we studied the behavior of Δpsl and Δpel mutants. Pel and Psl are the two main exopolysacharides produced by nonmucoid strains of *P. aeruginosa*. For both mutants, the maximum amount of biofilm in the microchannel was lower than for the wild type, thus indicating that these mutants are more easily detached by the flow. We also found that Psl had a stronger impact than Pel on the spatiotemporal dynamics and, in particular, on maximum clogging. The Δpsl mutant even featured major sloughing events for 2 *μL*/*min* that led to a near-complete removal of the biofilm. On the other hand, Pel featured a dynamics similar to that of the wild type, but with a reduced maximum clogging.

Although critical stresses are widely used in characterizing flow-induced detachment of biofilms, they have a broad range of different meanings and values. Critical stresses have been used to characterize a form of complete detachment from the solid surface (***Jiang et al., 2021***). ***Ohashi and Harada (1996)*** described an adhesion strength as the stress required to remove the biofilm – upon conceptualizing the problem as a surface to surface bonding, the adhesive strength has also been defined as the work required to detach the biofilm from the surface per surface area (in Joule per *m*^2^) (***Chen et al., 1998***). ***Ohashi and Harada (1996)*** found values for the shear strength in the hundreds of Pascals. ***Lau et al. (2009***) measured an adhesive pressure – adhesive force measured by microbead force spectroscopy divided by contact area – for *P. aeruginosa* PAO1 that was in the tens of Pascals. ***Körstgens et al. (2001***) studied the yield strength of a mucoid *P. aeruginosa* strain, discussing its role in the mechanical failure of the biofilm. Considering the biofilm as a viscoelastic gel with plastic flow properties, they found a stress at failure close to 1000 *Pa*. ***Lee et al. (2023***) show that bio-aggregates of *E. coli* at pore throats become fluidized above a critical value of the shear stress with a yield point at roughly 1.8 *Pa*. To establish a clear link between these values, our critical stress at 100-200 Pascals and the observed dynamics of sloughing in Stage II, future works will need to further refine our understanding of the role of the different variables on flow-induced detachment, including distributions of stresses within biofilms, adhesive strength to surfaces, mechanical properties of the biofilm, or bacterial regulation through quorum sensing (***Emerenini et al., 21 juil. 2015***) and/or rhamnolipids production (***Boles et al., 2005***).

### Self-sustained fluctuations

Fluctuations of the hydraulic resistance were previously reported in porous media flows. ***Howell and Atkinson (1976***) showed that, in trickling filters, constant influent concentration and operating conditions can yield large fluctuations induced by sloughing events. (***Stewart and Scott Fogler, 2002***) found very large oscillations in the pressure signal due to the formation of successive stable or unstable dextran plugs of *Leuconostoc mesenteroides* biofilm, behaving in a way similar to a yield stress fluid, and allowing flow only through breakthrough channels – see also discussions on the flow of yield stress fluids through porous media in (***Talon, 2022***). ***Sharp et al. (2005***) showed that “the pore channels are dynamic, changing in sized, number and location with time” in a flat plate reactor colonized by *Vibrio fischeri* biofilm. ***Kurz et al. (2022***) found successive cycles of growth and shear-induced detachment in the preferential flow paths for *Bacillus subtilis* biofilm development in a microfluidic system with cylindrical obstacles. ***Bottero et al. (2013***) modeled the coupled effects of flow, clogging and detachment to study the mechanisms that control these self-sustained fluctuations, in particular what leads to the clogging of a preferential flow path and the unclogging of another one – contrary to the development of stable flow paths.

Porous media have an inherent degree of complexity in the geometry that, combined with the nonlinear response of biofilms, can lead to a range of complex mechanisms. For instance, some of the works above-mentioned identified changes in the flow paths associated with the fluctuations. Our work shows that, even in a single channel with continuous flow, these fluctuations occur and prevent the system from reaching a true steady state. We therefore argue that the self-sustained fluctuations observed in Stage II are a signature of biofilm development in confined flows, even in the simplest geometries. The idea is that the strong bioclogging in confined geometries generates a pressure build-up that ultimately leads to sloughing, followed by re-growth. Our work therefore highlights the ubiquity of these fluctuations, which are likely playing an important role in the development and spreading of biofilms in a range of different systems, such as clogging in pipes, catheters (***Stickler, 2014***) or stents (***Guaglianone et al., 2010***), with applications ranging from infections, to environmental processes and engineering systems.

### Stochastic modeling

Our approach to modeling was constructed in two main parts. We first proposed differential equations to describe nutrient transport coupled with the dynamics of biofilm development in the channel, including growth and smooth detachment. The sharp jumps corresponding to sloughing were then added to this first layer of the model as a jump stochastic process. Although it seems quite natural to characterize the statistics of detachment (***Wilson et al., 2004***), there are actually very few attempts to treat such problems in the framework of stochastic processes. ***Howell and Atkinson (1976***) modeled sloughing in trickling filters through a conceptualization as connected filter units describing the discrete pieces of packing material in the filter. They introduced randomness by permitting sloughing at multiples of a fixed time interval. ***Bohn et al. (2007***) used an approach combining a logistic growth with random sloughing events through a stochastic differential equation. They described sloughing via a discrete expression of the amplitude of jumps occurring independently at each time step, which allowed them do describe daily fluctuations in light absorbance data for phototrophic biofilms development in a flow-lane incubator. In our approach, the jump process was characterized by two random variables: the interevent time between two successive jumps and the relative amplitudes of the jumps. Through the analysis of experimental data, we reconstructed the probability density functions for these random variables, with respectively Gamma and log-normal distributions.

There have been previous discussions (***Lewandowski et al., 2004***) on the determinism of biofilm formation, suggesting that sloughing events are intrinsically random events that generate large fluctuations, prevent the system from reaching a steady state and hinder the reproducibility of longterm experiments. Our work confirms that sloughing is integral to biofilm development (***Telgmann et al., 2004***) but shows that, although a true steady state is never reached, the fluctuations can be precisely characterized using stochastic modeling. This approach paves a way forward in terms of reproducibility: even though the state of the biofilm at any given time may not be reproducible, the randomness of the process may very well be.

Our approach to characterizing bursting events in terms of the distribution of the amplitude and interevent time is reminiscent of the description of other physical systems, such as avalanches (***Maaß et al., 2015***). For earthquakes, for example, a Gamma distribution for interevent times has been found in many different geographic regions (***Corral, 2004***). The fact that interevent times for biofilm sloughing also seem to follow a Gamma distribution may point towards specific physical mechanisms (***Kumar et al., 2020***), which could be used to better understand the physics of sloughing. One interesting perspective of this work is also to assess the universality of these distributions across microorganisms and whether we could define classes of bacteria with specific signatures on the stochastic process. With the same idea, we could further evaluate the impact of ecological interactions in multispecies biofilm or the effect of various molecules, such as biocides, on the sloughing dynamics.

## Material and Methods

### Bacteria and cultures

Experiments were performed using *Pseudomonas aeruginosa* PAO1 GFP (ATCC 15692GFP) strain, along with PAO1 GFP Δ*pslD* and Δ*pelF* mutants obtained from ***Colvin et al. (2011***, 2012) and built by non-polar deletion through allelic replacement of *pslD* and *pelF* operon and harbor pMRP9-1 plasmid expressing GFP. Bacteria were subcultured and grown in brain heart infusion (Sigma Aldrich, Saint-Quentin-Fallavier, France). Cultures were prepared from −80°C frozen aliquots spread on tryptic soy agar plate (Sigma Aldrich, Saint-Quentin-Fallavier, France) supplemented with 300 µg/mL of ampicillin (Sigma Aldrich) then incubated at 30°C during 24 h. Liquid cultures were prepared from the second subcluture on tryptic soy agar by spreading a 24 hours single colony diluted in brain heart infusion media supplemented with 300 µg/mL of ampicillin. 1*X* concentrated BHI media was prepared by dissolving 37 *g* of commercial powder in demineralized water and autoclaved with a liquid cycle (121°C for 15 minutes). 0.2*X*concentrated BHI media was obtained by dissolving 7.4 *g* while a third 1*X* solution was supplemented by 8 *g*/*L* of D-Glucose (Sigma Aldrich).

### Growth rate measurements in liquid culture

We determined growth curves for the first 35 hours of incubation using a microplate reader. All strains were refreshed from –80°C freezer stocks: 2 µL of frozen culture was diluted in 1 mL BHI supplemented with 300 µg/mL ampicillin. Initial optical density (OD_640_) was below the detection limit of the spectrophotometer. 100 µL of each suspension was transferred to a 96-well plate in an automatic microplate reader (TECAN-PC2017, 640 nm, 26°C). Note that microfluidic experiments were performed at room temperature. Each strain was measured in triplicate wells (technical replicates from the same biological sample) under static conditions, with 10 seconds of shaking at 90 rpm before each reading. Growth dynamics were measured every 10 minutes for 35 hours. For each well, 9 measurement points at different spatial positions were acquired and averaged at each time point. Doubling time was estimated during the exponential growth phase using the slope of the linear fit of the ln-transformed growth curve (doubling time = ln(2)/slope). The exponential phase was identified by excluding the lag phase, which was estimated using the DMfit tool (linear biphasic model).

### Microfabrication

Microchannels were fabricated using standard soft lithography techniques. Microchannel molds were prepared by depositing 100 µm SUEX sheets on a silicium wafer via photolithography. The negative mold was cleaned by isopropanol and silanized with trichloromethylsilane (Sigma Aldrich). Square cross-section channels had dimensions of 100 µm height by 100 µm width and 20 mm length. The chips were prepared with a 10% wt/wt cross-linking agent in the polydimethylsiloxane solution (Sylgard 184 Silicone Elastomer Kit, Dow Corning). PDMS was cleaned with isopropanol at 80 °C for 30 minutes and plasma-bonded to a clean glass coverslide.

### Inoculation and flow experiments

BHI suspensions were adjusted at optical density at *OD*_640 *nm*_= 0.2 (10^8^ *CF U*/*mL*) and inoculated inside the microchannels from the outlet, up to approximately ^3^/4 of the channel length in order to keep a clean inlet. The system was let at room temperature (25°C) for 3h under static conditions. Flow experiments were then performed at 0.02, 0.2, 2, 20 and 200 *μL*/*min* constant flow rates for 72h in the microchannels at room temperature. For the experiments at 0.2, 2, 20 and 200 *μL*/*min*, the fluidic system was based on a sterile culture medium reservoir pressurized by a pressure controller (Fluigent FlowEZ) and connected with a flow rate controller (Fluigent Flow unit). The flow rate was maintained constant by using a controller with a feedback loop adjusting the pressure in the liquid reservoir. The reservoir was connected to the chip using Tygon tubing (Saint Gobain Life Sciences Tygon™ ND 100-80) of 0.52 mm internal diameter and 1.52 mm external diameter, along with PEEK tubing (Cytiva Akta pure) with 0.25 mm inner diameter adapters for flow rate controller. The waste container was also pressurized by another independent pressure controller to reduce air bubble formation in the inlet part. For the experiments at 0.02 *μL*/*min*, we used an Harvard Phd2000 syringe pump for the flow.

### UVC irradiation

The biofilm is constrained in a part of the microchannel using UVC irradiation directly through the PDMS of the microfluidic chips, thus reducing contamination risk and avoiding unwanted progression/growth of *P. aeruginosa* in the inlet and tubing for several days of experiment (***Ramos et al., 2023***). This eliminates parasitic consumption of nutrients in various parts of the fluidic system and maintain a controlled boundary condition with a fixed concentration of nutrients at the inlet of our zone of interest.

To this end, the inlet and outlet of the microchannels were exposed to UVC light by a system of UVC LEDs (***Ramos et al., 2023***). The system consists of a 3D printed part called the guide carrying in its backside a PCB with a LED light source that delivers UV-C light of 1W power. The light beam follows a straight trajectory until a 45° mirror positioned in the front side of the guide, which reflects the light parallel to the PDMS chip and irradiate it. UVC guides were positioned in both sides of the PDMS microchannel and separated by a 1.2 cm distance to keep a central area unexposed. UVC power is measured by a radiometer after mirror reflection and delivers a power of 200 *mW* /*m*^2^equivalent to 2 *mJ* /*cm*^2^. The guide is elevated to fit with the PDMS chip dimensions and contains a barrier located in the front side that blocks the diffusion of light through the PDMS polymer in the horizontal direction.

### Mass spectrometry analysis

Mass spectrometry analysis was performed on a Thermo Scientific Q Exactive Focus Orbitrap LCMS/MS system connected to a LC device Thermo Scientific Vanquish UPLC system with PDA detection. Analytical separations were performed on a 150 x 2.1 mm Thermo Hypersyl Gold C18 column (1.9 μm) using an MeCN/H2O 1% formic acid gradient. Data were captured in full MS scan mode and processed using Chromeleon 7 software.

### Imaging for the biofilm experiment

Bacterial development was imaged for a period of 72 hours with a timestep of 30 minutes at 25°C on an inverted microscope (Ti-2E, Nikon) using a digital camera (back-illuminated PCO edge). Time-lapse images were acquired using brightfield and fluorescence microscopy (Sola light source 10% intensity with 30 ms exposure with 500 *nm* excitation and 513 *nm* emission combined with (FITC filter). Images were obtained with a focal plan at the glass/liquid interface. These images had dimensions of 30086 x 154 pixels obtained after multi position scanning using automatic Nikon platform and assembled by Nikon NIS software of single images with 0.65 µm/pixel using a 10X magnification Nikon objective (NA = 0.3).

### Image analysis for the biofilm distribution in the longitudinal direction

Fluorescence images were loaded as a matrix (30086 x 154) in MATLAB (MathWorks). For each time acquisition, the signal was first integrated in the transverse direction to obtain a mean distribution in the longitudinal direction. In plotting curves to analyze the effect of nutrients, all timepoints were then averaged to obtain one dimensional curves of the mean longitudinal profile. For kymographs, one dimensional curves for each time point were stacked together to describe the spatiotemporal dynamics. The intensity values were normalized to the maximum values of each replicate over all times. For each flow and nutrient condition, three biological replicates were performed.

### Image analysis for the biofilm segmentation

To estimate changes in channel colonization, GFP images were binarized using a machine learning software (Ilastik) (***Berg et al., 2019***). Images were pre-treated with imageJ (***Schneider et al., 2012***) by normalizing all pixel values between 0 and 65535 grayscale levels. In Ilastik, biofilm structures were first differentiated from the empty flow path using pixel-level manual labeling during pixel classification where visible patches of biofilm and empty channel were annotated manually by mouse cursor. Pixel classification workflow employs a Random Forest classifier, known for its generalization properties. Several samples of biofilm and background images were used to train the classifier by annotating pixels with corresponding labels, allowing the algorithm to learn and make predictions in real-time. The chosen features were color, intensity, edges, and texture. The generated probability maps indicating the likelihood of each class at every pixel were used for the object classification and were thresholded at a value of 0.6 with no size filter. Thresholding is a process involved in converting continuous probability maps generated from pixel classification into binary segmentation images by setting a threshold, where pixels above the threshold are classified as belonging to an object. The size, intensity, position and convexity of the biofilm objects was exported in .cvs format and further analysed in matlab. Volumic fraction of biofilm in microchannel was calculated from the sum of the size of segmented objects divided by the interest growth area (non UVC irradiated central part of microchannel).

### Initial adhesion

Separate experiments were performed to study the behavior of cells in the initial phases of attachment in order to increase spatial and temporal resolution. Liquid cultures were prepared following the same protocol as described previously (see Bacteria and cultures). Sterile 1× concentrated BHI culture medium supplemented by 300 *μg*/*mL* of ampicillin was flowed under constant flow rate (*Q* = 0.2 *μL*/*min*, 2 *μL*/*min* and 20 *μL*/*min*) after 3 hours under static conditions at 26° C.

Images were obtained with a 40× Nikon objective (NA = 0.95) using a differential interference contrast (DIC) brightfield with 2 minutes per frame during 3.5 Hours. Replicates were obtained by imaging four positions per channel and each condition was performed in two distinct channels forming two distinct biological replicates to obtain n = 8 replicates by condition (*Q* = 0.2 *μL*/*min*, 2 *μL*/*min* and 20 *μL*/*min*). The images were further segmented using Ilastik with the same parameters as described before (see Image analysis for the biofilm segmentation) and single cells were selected as objects with an area higher than 20 *μm*^2^ to avoid counting dust particles and artefacts that could be considered as distinct objects by Ilastik. The doubling time was calculated in the window between 0 and 3.5 hours by a a linear fitting of the logarithm of the number of cells. The slope was used to estimate growth rate and doubling time. Cell count was calculated from image segmentation of four positions in two channels to generate 8 replicates by condition (n= 8) for (*Q* = 0.2 *μL*/*min*, 2 *μL*/*min* and 20 *μL*/*min*).

### Wavelet analysis

Time series of the pressure data were investigated with a wavelet analysis to identify temporal variations of spectral power (***Torrence and Compo, 1998***). Wavelet analysis was carried out using a Morlet wavelet, the product of a sinusoidal wave and a Gaussian envelope, with a frequency parameter of 6 and scale width of 300. We then applied a continuous wavelet transform (CWT) results which gives wave power coefficients dependent on the scale or period and the time, as well as a cone of influence (COI), where edge effects become important. The time series were zero-padded to reduce the edge errors. The curved envelope at the bottom of the scalogram in Fig 3 represents the cone of influence, where edge effects limit the reliability of wavelet coefficients at longer periods near the beginning and end of the time series.

### Data analysis for the calculation of the hydraulic resistance and volume fraction

Our approach to quantifying biofilm volume fraction relies on pressure drop measurements, which inherently capture the three-dimensional architecture of the biofilm. The uniform layer assumption is supported by: (i) the excellent quantitative agreement between predicted and measured scaling laws (*ϕ*_max_ ∝ *Q*^1/2^, Fig 7f), (ii) the consistency of *ϕ*_max_ values across different flow rates and replicates, and (iii) the strong correlation between model-derived *ϕ*(*t*) and integrated fluorescence intensity (Figs 3 and 7). While fluorescence microscopy provides information on a limited depth in the channel, its agreement with the pressure-based *ϕ*(*t*) suggests it captures the overall colonization dynamics.

Recorded pressure fluctuations in the reservoir were converted to hydraulic resistance in the 10 mm zone between UVC LEDs where biofilm develops. We write the difference between the two pressure reservoirs (one at the inlet and one at the outlet) as Δ*P*_res_ and express it as

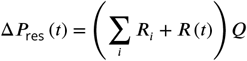

with *Q* the imposed flow rate, ∑_*i*_ *R*_*i*_ the sum of the hydraulic resistance of the different part of the hydraulic network and *R* (*t*) the hydraulic resistance of the the 10 mm zone between UVC LEDs. To estimate *R* (*t*), we evaluated ∑_*i*_ *R*_*i*_ from the mean value of Δ*P*_res_, which we write 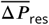, over a few hours of the experiment where biofilm growth is not yet observable. We then use

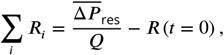

which yields

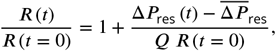

with *R* (*t* = 0) in the square cross-section of size *h*_0_ = 100 *μm* that can be estimated as

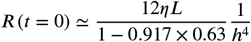

with *η* the viscosity of the culture medium at 25°C, *L* = 10 *mm* the length of the channel and *h* the width/height of the channel. Since we considered 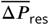 for the normalization, and because the low pressure signal is relatively noisy at the beginning, we also only use the part of the signal for which 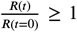 – it is a important, for instance, to obtain non-negative volume fractions of biofilm that are only the result of the initial noise. Upon assuming that the biofilm forms a uniform layer on the sides of the channel (see schematics in Supplementary Section I), we also have

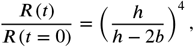

with *b* the thickness of the biofilm layer. The volume fraction of biofilm is 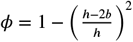 so that

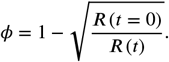

To validate this approach, we performed correlation analysis between *ϕ* calculated from hydraulic resistance and *ϕ* estimated from integrated fluorescence intensity. Strong positive correlations (r = 0.68–0.77, p < 0.0001) were observed across all flow rates, confirming that pressure-based measurements accurately capture biofilm dynamics (Supplementary Figure S21).

### Frequency analysis

Welch power spectral analysis ***Welch (1967)*** of the pressure signal was carried out by dividing the time signal into smaller segments, calculating their periodogram and finally averaging accross the frequencies, resulting in a power spectral density (PSD) estimate. This method allows for PSD estimate that is less noisy than usual periodograms. Here, we used the Matlab tool pwelch to estimate the PSD using hanning window with an overlap of 50%. A linear fit was then applied on the interval in the low frequency range.

### Construction of the probability density functions for jumps and fits

Volume fraction data were processed using a homemade Matlab code. Each dataset was first subsampled by keeping only 1 in 150 points. The subsampled signal was then differentiated and only negative values corresponding to detachment events were conserved. Jumps were then identified through the selection of local minima. For each flow rate, all the values for the times between two successive jumps and the relative amplitude of the jumps – amplitude of the jump relative to the value of the volume fraction just before the jump – for the different replicates were aggregated to construct the distributions.

Gamma distributions were then fitted to the experimental data for the times between successive jumps and lognormal distributions were used for the amplitude of the jumps. Values for the different parameters are summarized in Table.2.

### Numerical simulation of the stochastic process

Here we describe how we simulated the evolution of the volume fraction *ϕ*, solving

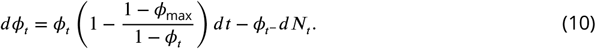

The simulation was done using a homemade Matlab code. For each case, based on the previously calculated distributions, we first generated two sets of random numbers corresponding to the times between successive jumps and to the relative amplitude of the jumps – therefore allowing us to completely determine the jump process, *N*. We then simply solved the ordinary differential equation

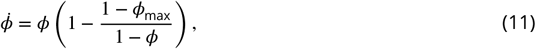

between jumps using the Matlab solve ode45. Upon reaching a jump, the simulation was stopped and the jump implemented, before proceeding to treating the next interval.

### Comsol flow simulations

Flow simulation was performed using a finite element approach in Comsol v6.1. The channel was treated as a rectangle of 850 *μm* length by 100 *μm* width. Biofilm, as shown in white in Fig 6, was added from a segmentation of real experimental images. We then solved incompressible Stokes flow with no-slip/no-penetration boundary conditions on the solid and on the biofilm surface. The inlet condition was an imposed velocity, while the outlet was an imposed pressure.

## Supporting information

Main supplementary file

## Acknowledgments

This work is part of a project that has received funding from the European Research Council (ERC) under the European Union’s Horizon 2020 research and innovation programme (grant agreement no 803074). This work was partially supported by the LAAS-CNRS micro and nanotechnologies platform, a member of the French Renatech network. We thank the entire BEBOP team for daily interactions and assistance; Julien Lefort and Emmanuel Libert for their technical support; Terence Desclaux for the introduction to wavelet analysis; Matthew R. Parsek and Maria Zori for providing the Pel and Psl mutants; and Laure Latapie for her support with mass spectrometry.

## Supplementary information

### I. MODELS

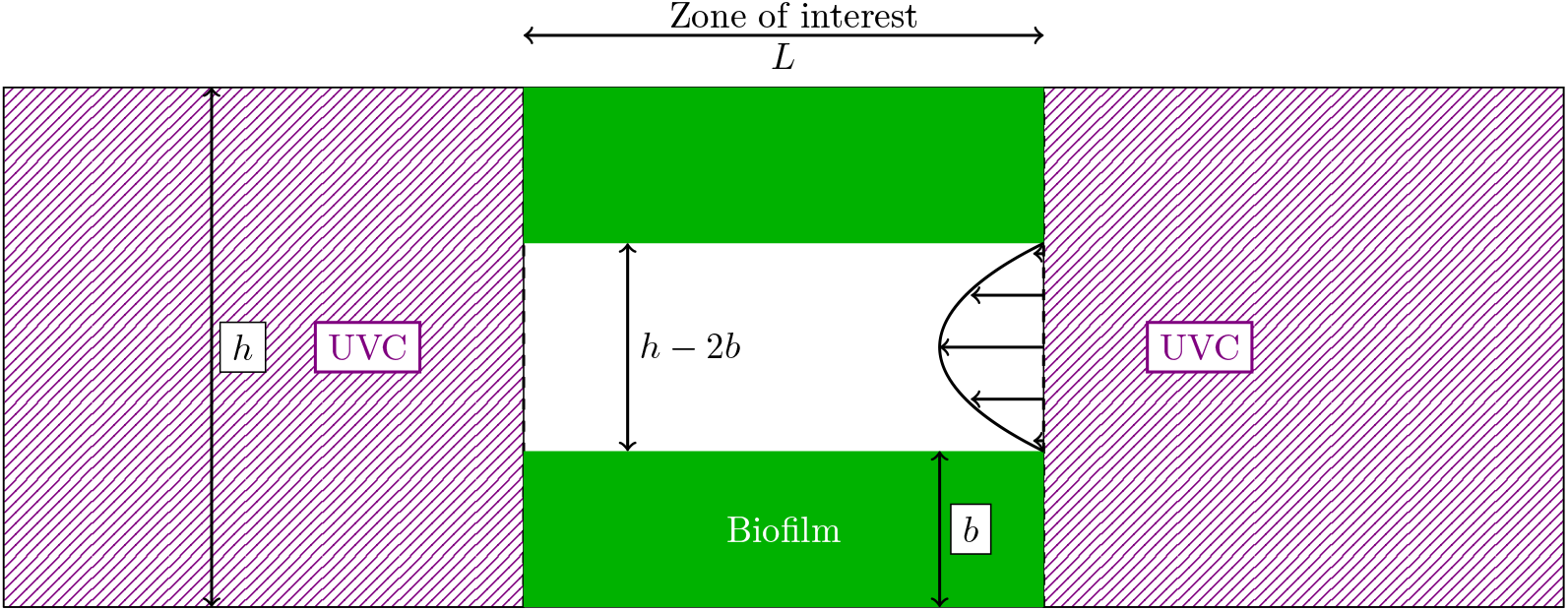

#### 1. Nutrient transport

The limiting nutrient is treated as a solute being transported by advection/diffusion and consumed by bacteria. We consider a cross-section average concentration of the form *C* = ∫_*A*(*x*)_ *c* (*x, y, z*) *dydz* with *A* the section at any given *x* – *x* being the longitudinal direction. The transport reads

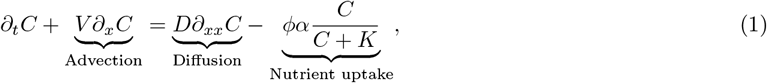

where *C* [*kg* ×*m*^−3^] is the concentration, *x* is the longitudinal coordinate system, *ϕ* is the cross-section volume fraction of biofilm, *V* [*m* × *s*^−1^] the average velocity in the empty channel, *D* = 10^−9^*m*^2^ × *s*^−1^ an estimate diffusion coefficient of the solute, *α* [*kg* × *m*^−3^ × *s*^−1^] the uptake rate and *K* [*kg* × *m*^−3^] the half-saturation constant. As we impose a constant flow rate for each experiment, the velocity averaged over the entire cross-section remains constant, even for different values of *ϕ*. The term *ϕ* in the nutrient uptake indicates that consumption is proportional to the volume fraction of biofilm.

We considered that nutrient transport is quasi-steady, since solute transport is much faster than growth. We further nondimensionalized this equation as

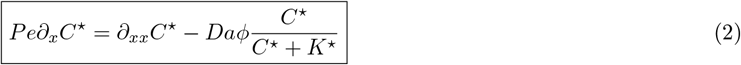

with 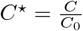 with the inlet concentration *C*_0_ [*kg* × *m*^−3^] the length of the channel *L* = 10 *mm*, the Péclet number 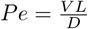 and *Da* the Damköhler number defined as the ratio of diffusive to reactive times, 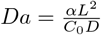.

#### 2. Flow

We treat the biofilm as a uniform layer on the solid surface, so that the flow channel always has a square cross-section. We also neglect biofilm porosity at this scale and consider a no-slip boundary condition at the boundary with the fluid. Porosity is a component of biofilm structures, resulting from the polymeric nature of the EPS matrix, mechanical forces, and biological processes such as cell death or predation. When considering flow-biofilm interactions, this porosity may allow fluid flow through the biofilm, with reported permeability values spanning an extremely broad range from 10^−15^ to 10^−7^ (Kurz et al., 2023). However, we argue that biofilm permeability is not the primary driver in our system:

- In microscopy visualization, our biofilms form dense structures where flow around the biofilm through narrow channels dominates over flow through the porous biofilm matrix.
- We performed microrheology experiments (data not shown) in these biofilms by imaging the Brownian motion of nanoparticles in the biofilm. Their trajectories indicate that, in our conditions, the viscoelastic flow of the biofilm itself largely dominates over the flow of culture medium through the biofilm matrix.
- We argue that the extreme variability in reported permeability values (spanning several orders of magnitude, Kurz et al., 2023) reflects not only differences in experimental systems, but also fundamental challenges in defining and measuring permeability for viscoelastoplastic biofilms (the biofilm itself is actually flowing). Given this uncertainty, incorporating permeability into our model would introduce parameters that cannot be reliably constrained from literature or independently measured in our setup. Our approach (i.e. treating the biofilm as impermeable and focusing on flow obstruction) avoids this parametrization complexity while successfully capturing the observed dynamics.
- Our model successfully predicts the observed scaling laws (main text Fig. 7f) and hydraulic resistance dynamics (main text Fig. 3) without invoking permeability, suggesting that flow obstruction rather than flow penetration is the dominant mechanism.

With these hypotheses, we write the pressure drop across the channel as

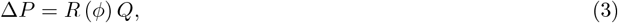

with Δ*P* the pressure difference, *Q* the flowrate and *R* the hydraulic resistance (which is a function of the volume fraction of biofilm). The hydraulic resistance in the empty channel is

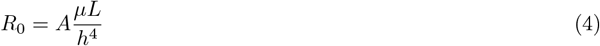

With 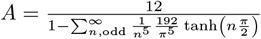 – which can be approximated as 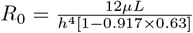 with 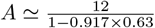. When the biofilm grows as a uniform layer, we also have

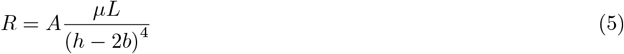

To obtain an expression in terms of the volume fraction of biofilm, we use 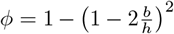 and 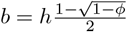 – and also *h* − 2*b* = *h* (1 − *ϕ*)^1/2^. We thus have

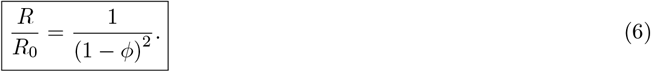

#### 3. Hydrodynamic stresses

We first write *σ*_0_ the shear stress on the liquid/solid surface of the channel when *ϕ* = 0. To obtain an explicit expression for this stress, the easiest way to proceed is to notice that, in the *x* direction, tangential to the surface of the channel, the force exerted by the pressure difference between the inlet and outlet is equal to the viscous force exerted by the fluid on the solid. This can be easily shown by integrating ∇ · ***σ*** = 0, with ***σ*** = −*p***I** + *µ* ∇**v** +^T^ ∇**v** and *p* the pressure and *µ* the viscosity, over a control volume that is the the zone of interest between UVCs in the channel. With 4*hL* the solid surface in the zone of interest, we have *σ*_0_4*hL* = *h*^2^Δ*P* where *L* is the length of the control volume, *h* is the width of the square channel and Δ*P* is the pressure difference between inlet and outlet. We therefore have

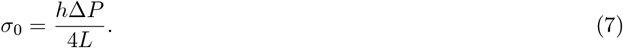

We can also write Δ*P* = *R*_0_*Q* with the hydraulic resistance in the square cross-section channel, 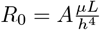. Therefore, we have

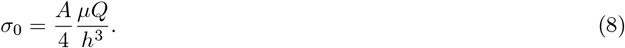

With the same type of reasoning, the total force along *x* applied by the fluid on the biofilm, including both shear and pressure is 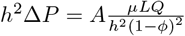. Therefore, the total stress on the biofilm solid surface is

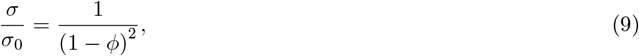

with a contribution from pressure

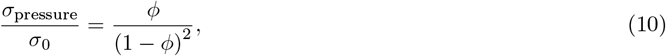

and from shear

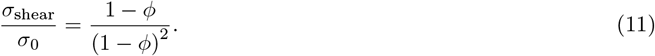

Also note that the shear stress at the biofilm fluid interface is

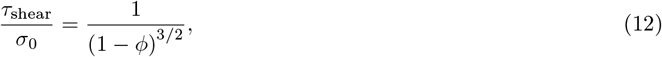

with the difference being simply the ratio of biofilm/solid and biofilm/fluid surfaces.

#### 4. Biofilm growth

We model biofilm development as

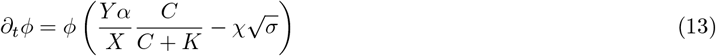

with *Y* a yield coefficient (mass of biofilm produced by mass of limiting nutrient consumed), *X* the density of the biofilm *X* [*kg* × *m*^−3^] and *χ* the sensitivity to biofilm detachment 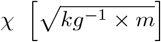. We non-dimensionalize with time 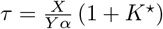 in order to obtain

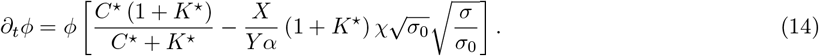

Recalling that 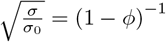 and writing the dimensionless number 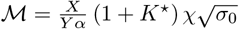, we obtain

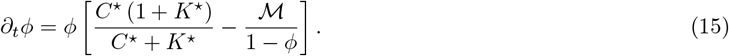

Assuming that ℳ ≤ 1, which is consistent with the observation that biofilm is never completely removed from the channels, we can also write this as

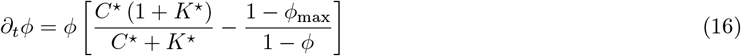

with *ϕ*_max_ the maximum value of *ϕ*.

In the case with no nutrient limitation, we assume *C*^⋆^ = 1 everywhere in the channel, so that we have

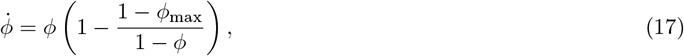

with *ϕ* describing an average volume fraction in the entire channel, and thus being only a function of time. We can also describe sloughing by modifying this equation as

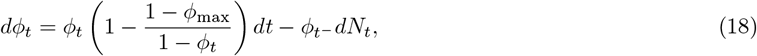

with *ϕ*_*t*_ describing the average fraction of biofilm in the microchannel, *N* the jumps and 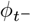 the value of *ϕ* at time *t*^−^ just before the jump.

## II. SUPPLEMENTARY FIGURES

**Figure S1.**
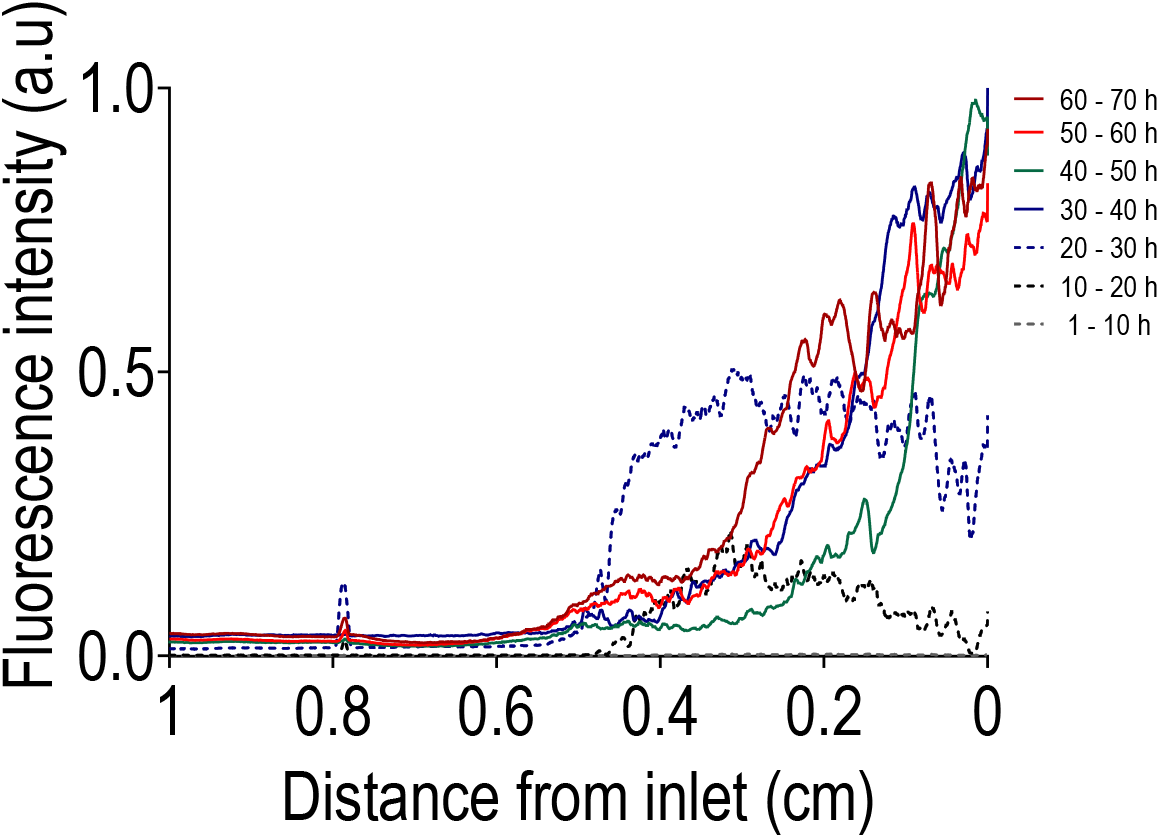
Time evolution (10 hours by curve) for *Q* = 0.02 *µL/min* of the averaged (n = 3) longitudinal distribution of the biofilm and the impact of nutrient limitation.

**Figure S2.**
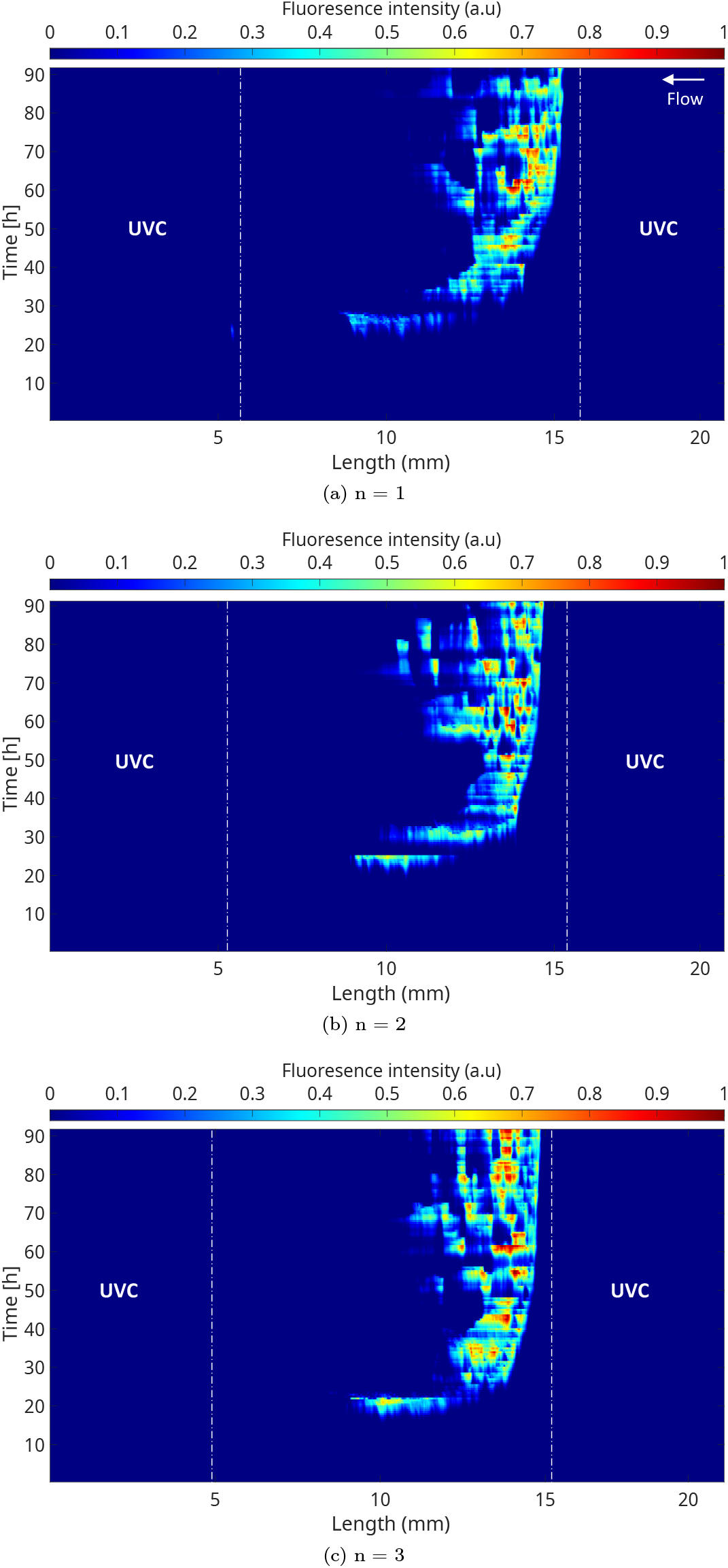
Kymograph representations based on the GFP signal for *Q* = 0.02 *µL/min* with [1X] BHI showing the spatio-temporal dynamics of the biofilm longitudinal distribution. Fluorescence intensity signal was normalized to the maximum signal intensity value.

**Figure S3.**
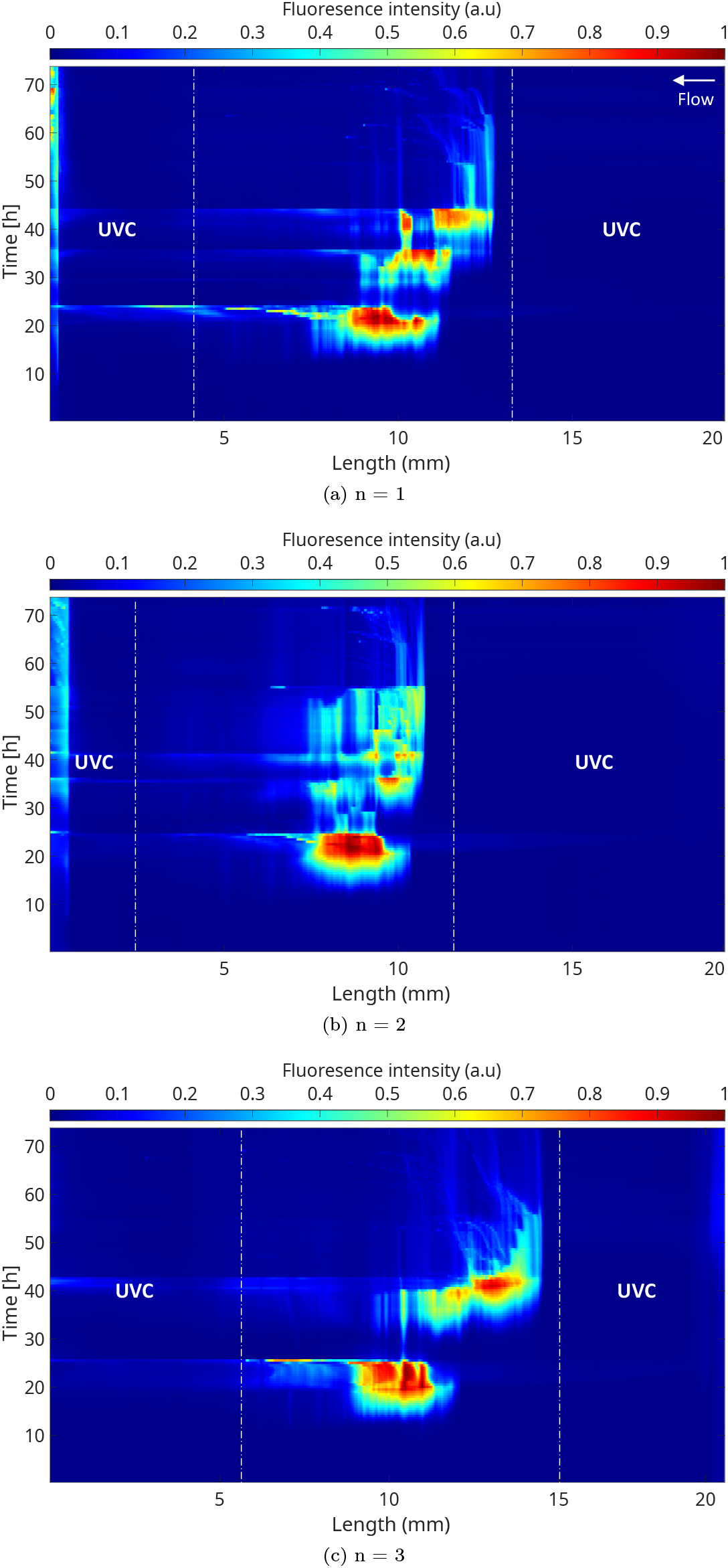
Kymograph representation for *Q* = 0.02 *µL/min* with [1X] BHI supplemented by 8 *g/L* glucose. Fluorescence intensity signal was normalized to the maximum signal intensity value.

**Figure S4.**
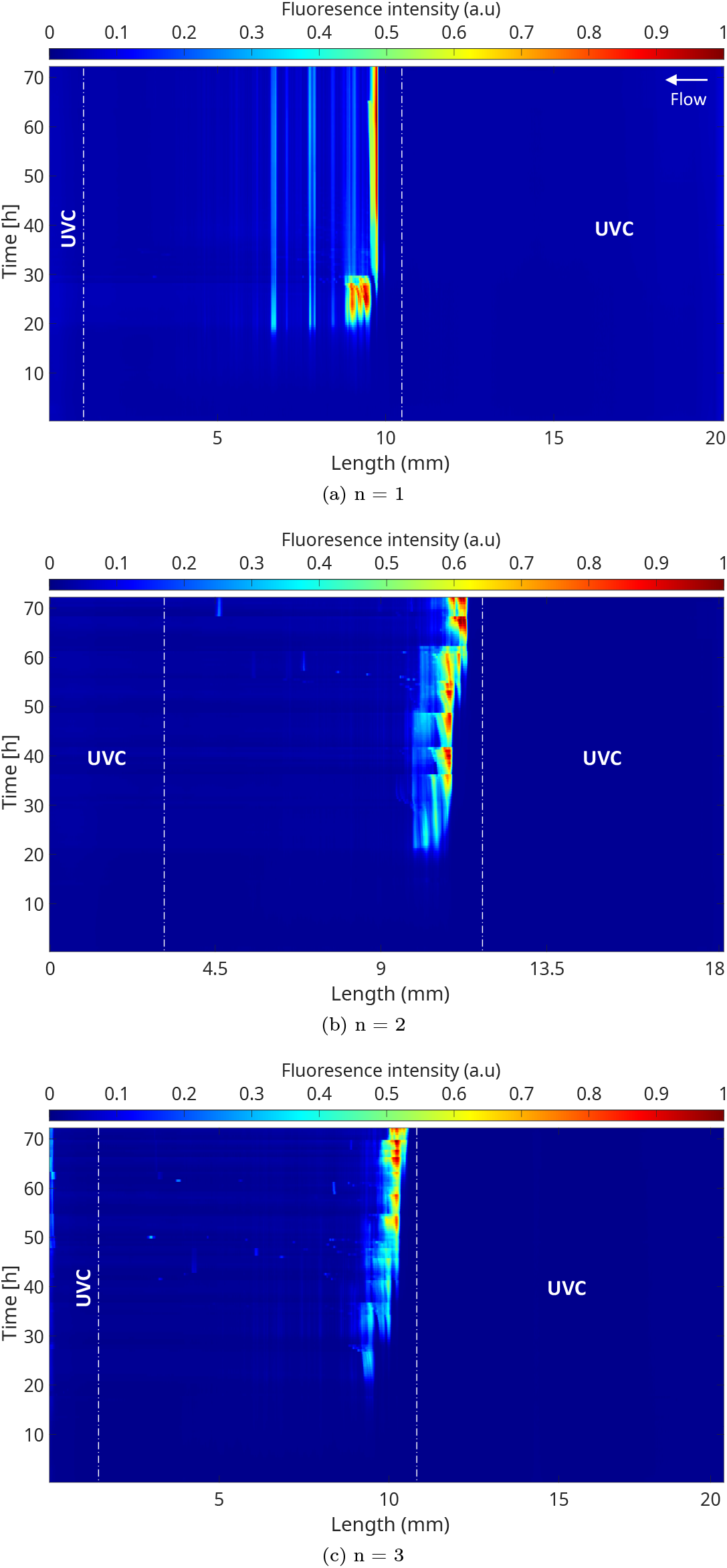
Kymograph representation for *Q* = 0.02 *µL/min* with [0.2X] BHI. Fluorescence intensity signal was normalized to the maximum signal intensity value.

**Figure S5.**
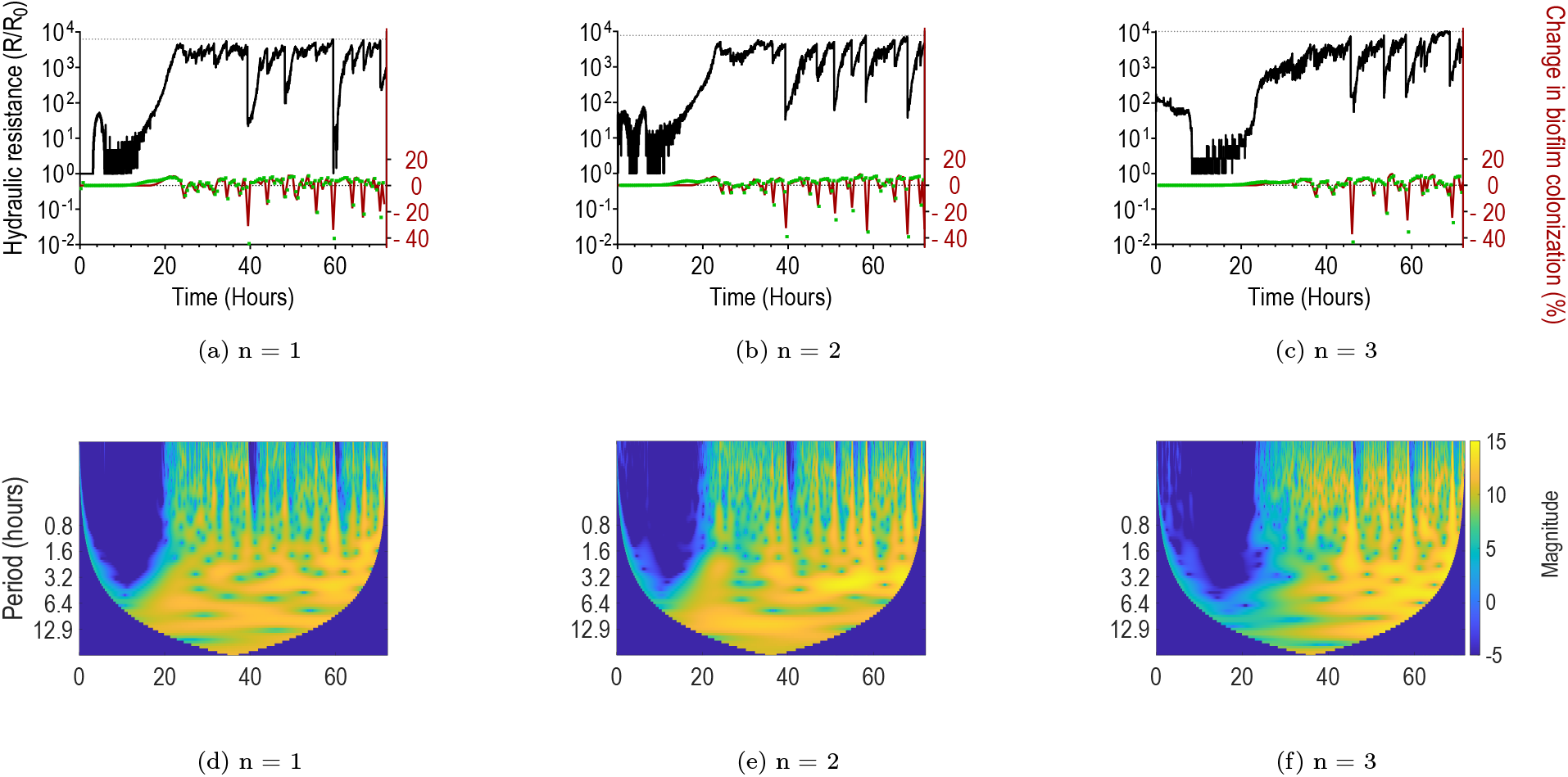
Temporal dynamics of growth and detachment for *Q* = 0.2 *µL/min*. (a), (b) and (c) show the variation of hydraulic resistance (black line) for three replicates, along with the percentage of colonization extracted from image segmentation (red line) and the integrated fluorescence intensity signal (green). Thin dotted lines on the top of y axis indicate the maximum value reached by hydraulic resistance. (d), (e) and (f): corresponding wavelet scalograms for the pressure signal.

**Figure S6.**
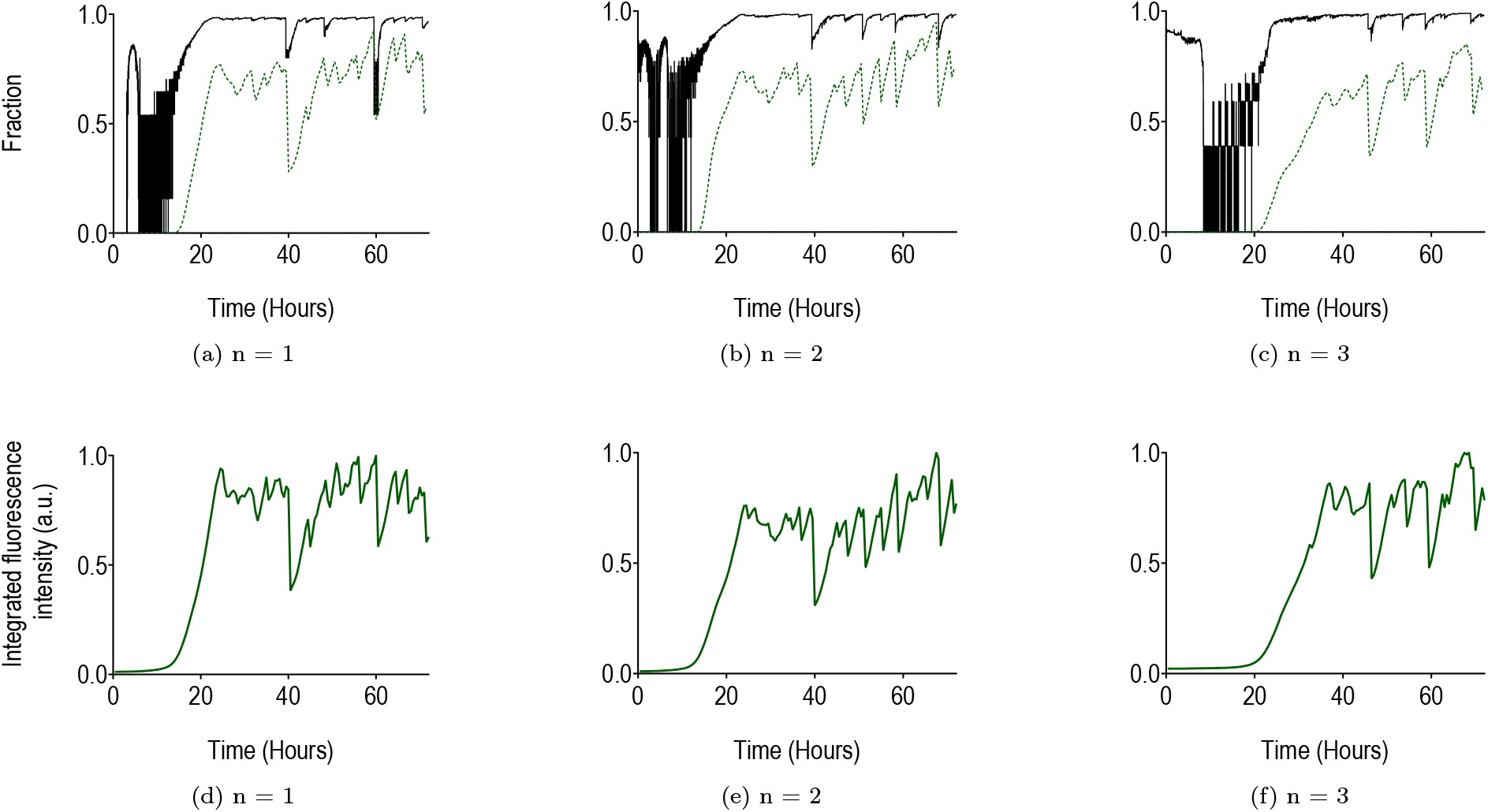
Temporal dynamics of growth and detachment for *Q* = 0.2 *µL/min*. (a), (b) and (c) show the evolution of the volume fraction as a function of time, with that calculated from hydraulic resistance (black line) and from segmentation (green dotted line). (d), (e) and (f): normalized integrated fluorescence intensity signal as a function of time.

**Figure S7.**
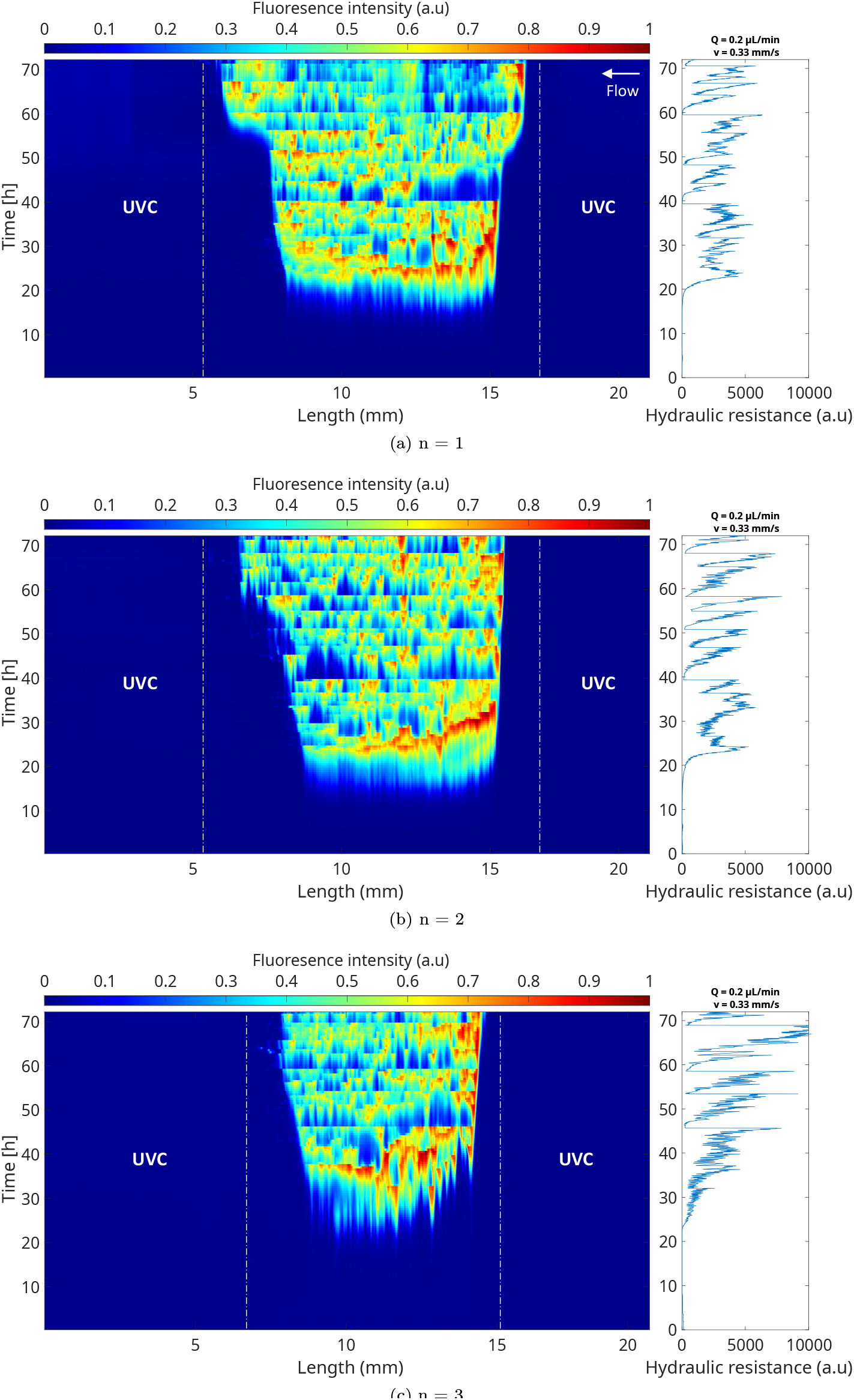
Kymograph representation for *Q* = 0.2 *µL/min* with [1X] BHI. Fluorescence intensity signal was normalized to the maximum signal intensity value. Plots on the right show the corresponding evolution of the hydraulic resistance.

**Figure S8.**
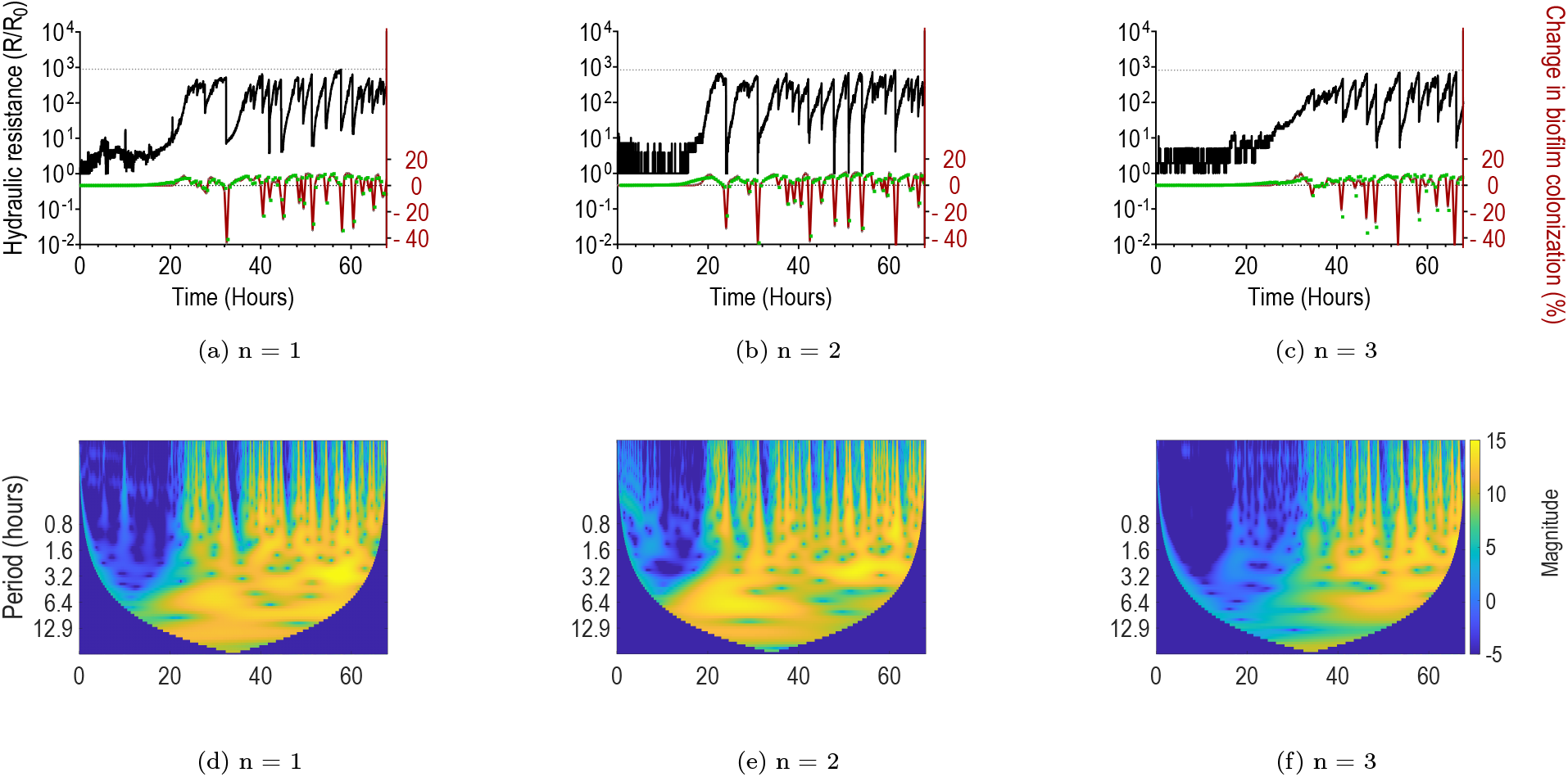
Temporal dynamics of growth and detachment for *Q* = 2 *µL/min*. (a), (b) and (c) show the variation of hydraulic resistance (black line) for three replicates, along with the percentage of colonization extracted from image segmentation (red line) and the integrated fluorescence intensity signal (green). Thin dotted lines on the top of y axis indicate the maximum value reached by hydraulic resistance. (d), (e) and (f): corresponding wavelet scalograms for the pressure signal.

**Figure S9.**
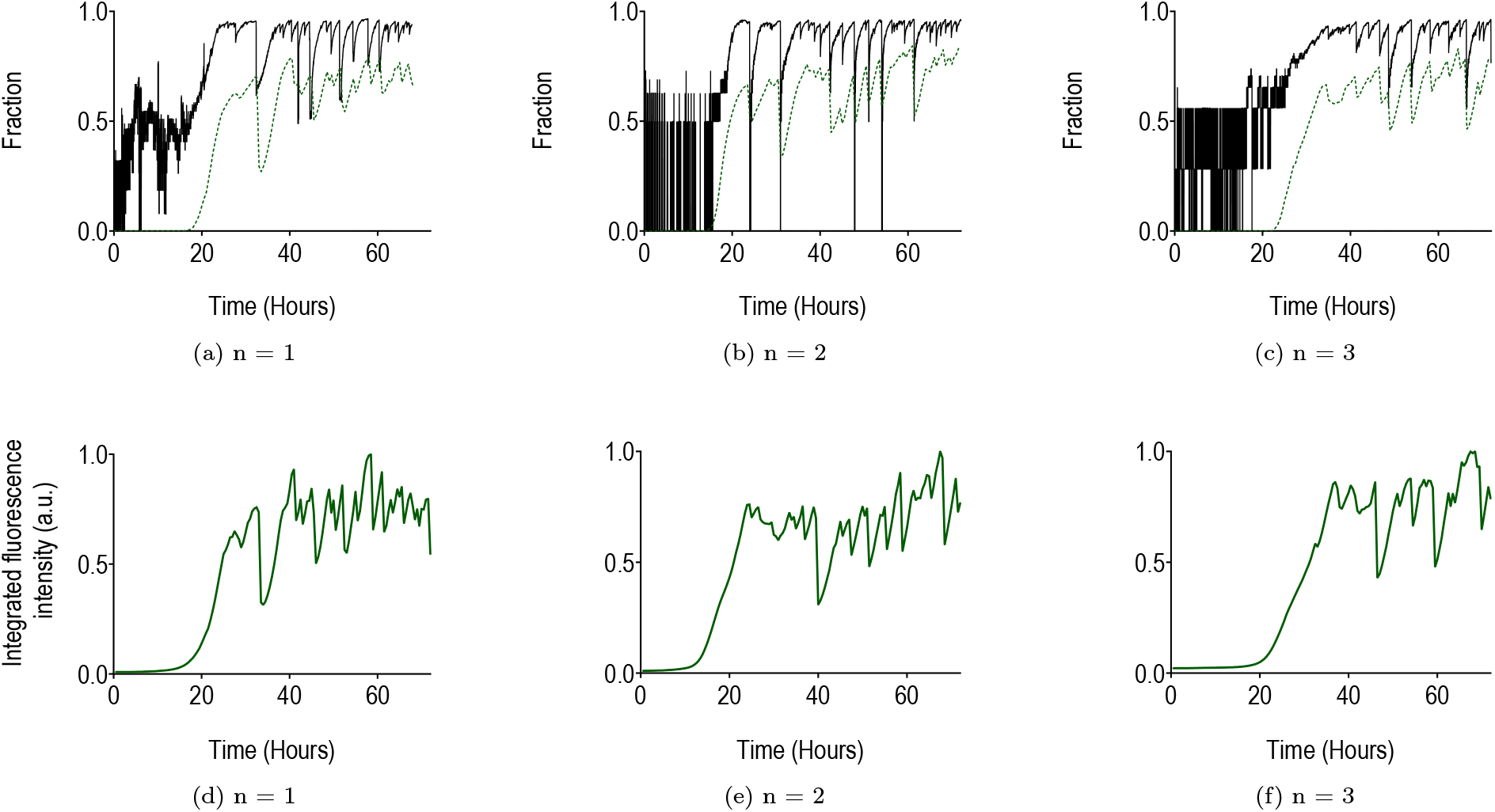
Temporal dynamics of growth and detachment for *Q* = 2 *µL/min*. (a), (b) and (c) show the evolution of the volume fraction as a function of time, with that calculated from hydraulic resistance (black line) and from segmentation (green dotted line). (d), (e) and (f): normalized integrated fluorescence intensity signal as a function of time.

**Figure S10.**
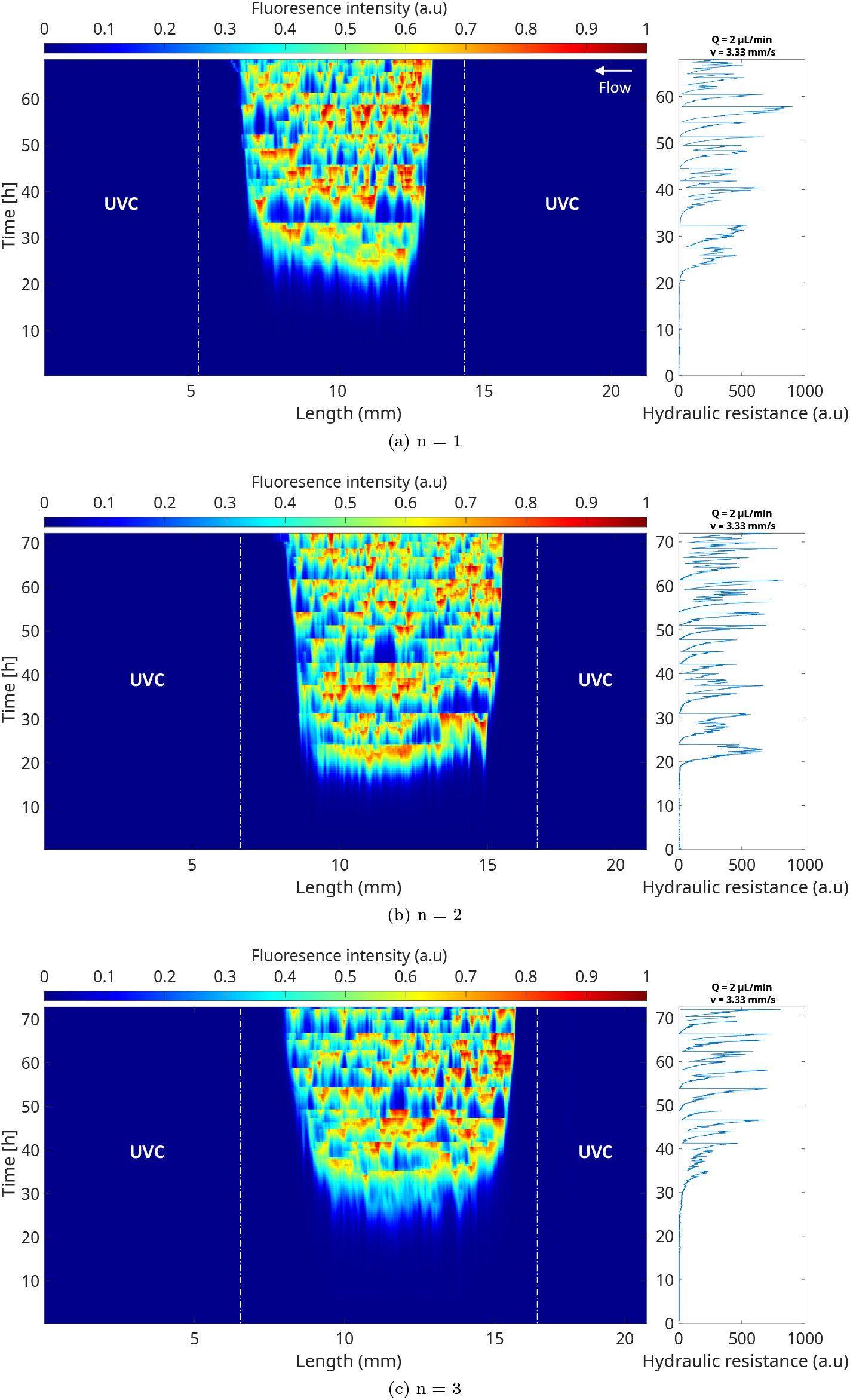
Kymograph representation for *Q* = 2 *µL/min* with [1X] BHI. Fluorescence intensity signal was normalized to the maximum signal intensity value. Plots on the right show the corresponding evolution of the hydraulic resistance.

**Figure S11.**
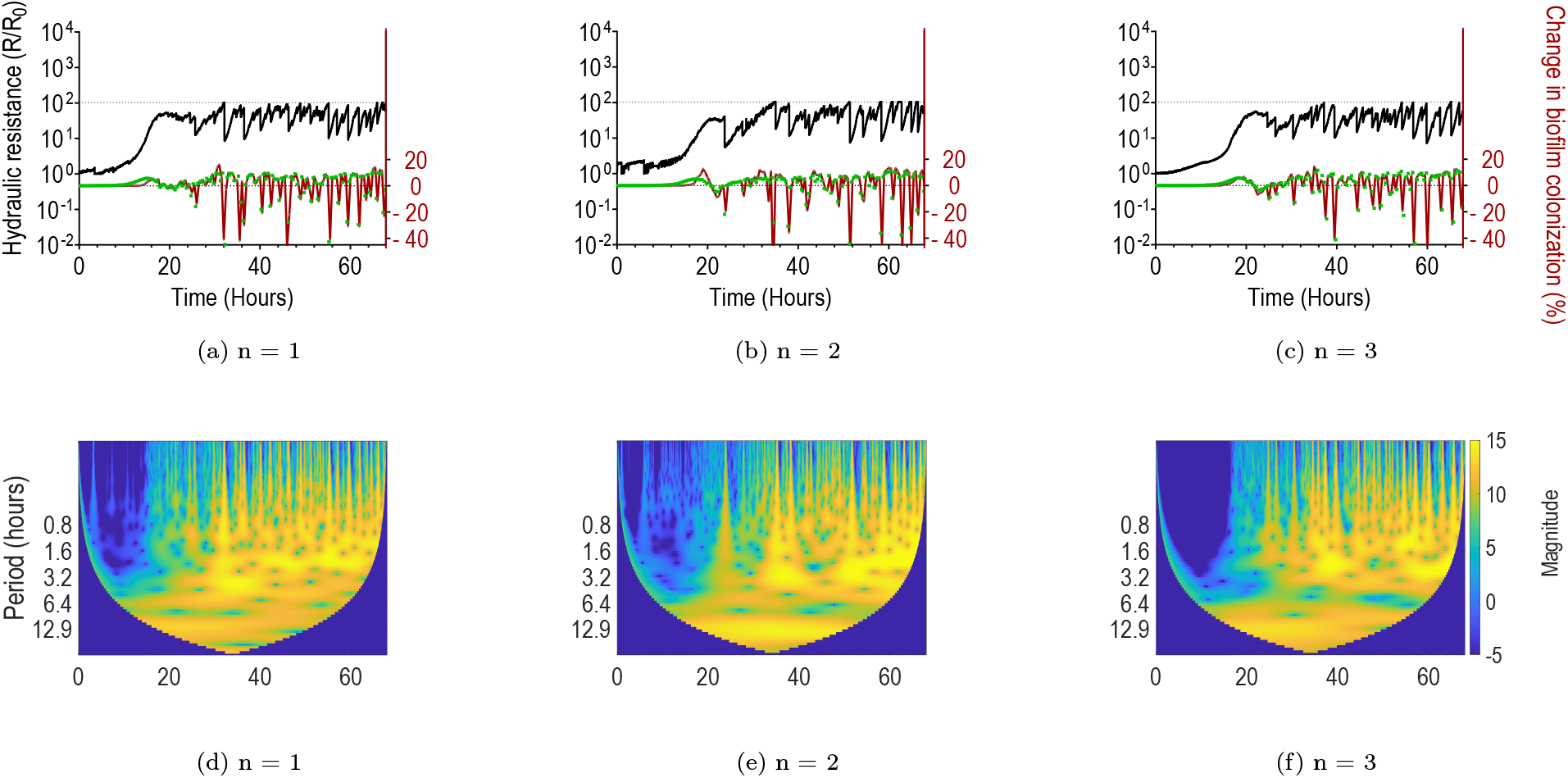
Temporal dynamics of growth and detachment for *Q* = 20 *µL/min*. (a), (b) and (c) show the variation of hydraulic resistance (black line) for three replicates, along with the percentage of colonization extracted from image segmentation (red line) and the integrated fluorescence intensity signal (green). Thin dotted lines on the top of y axis indicate the maximum value reached by hydraulic resistance. (d), (e) and (f): corresponding wavelet scalograms for the pressure signal.

**Figure S12.**
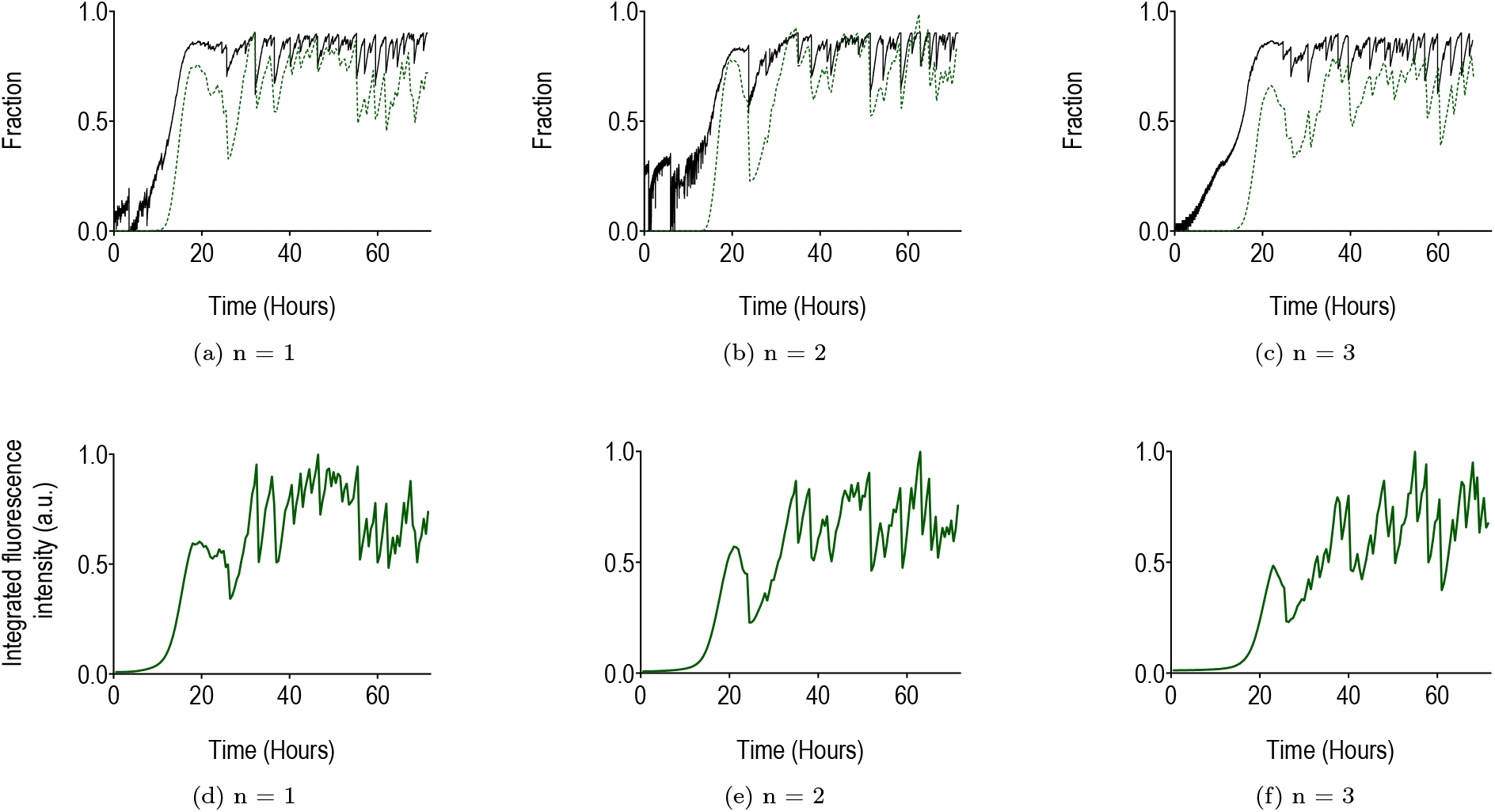
Temporal dynamics of growth and detachment for *Q* = 20 *µL/min*. (a), (b) and (c) show the evolution of the volume fraction as a function of time, with that calculated from hydraulic resistance (black line) and from segmentation (green dotted line). (d), (e) and (f): normalized integrated fluorescence intensity signal as a function of time.

**Figure S13.**
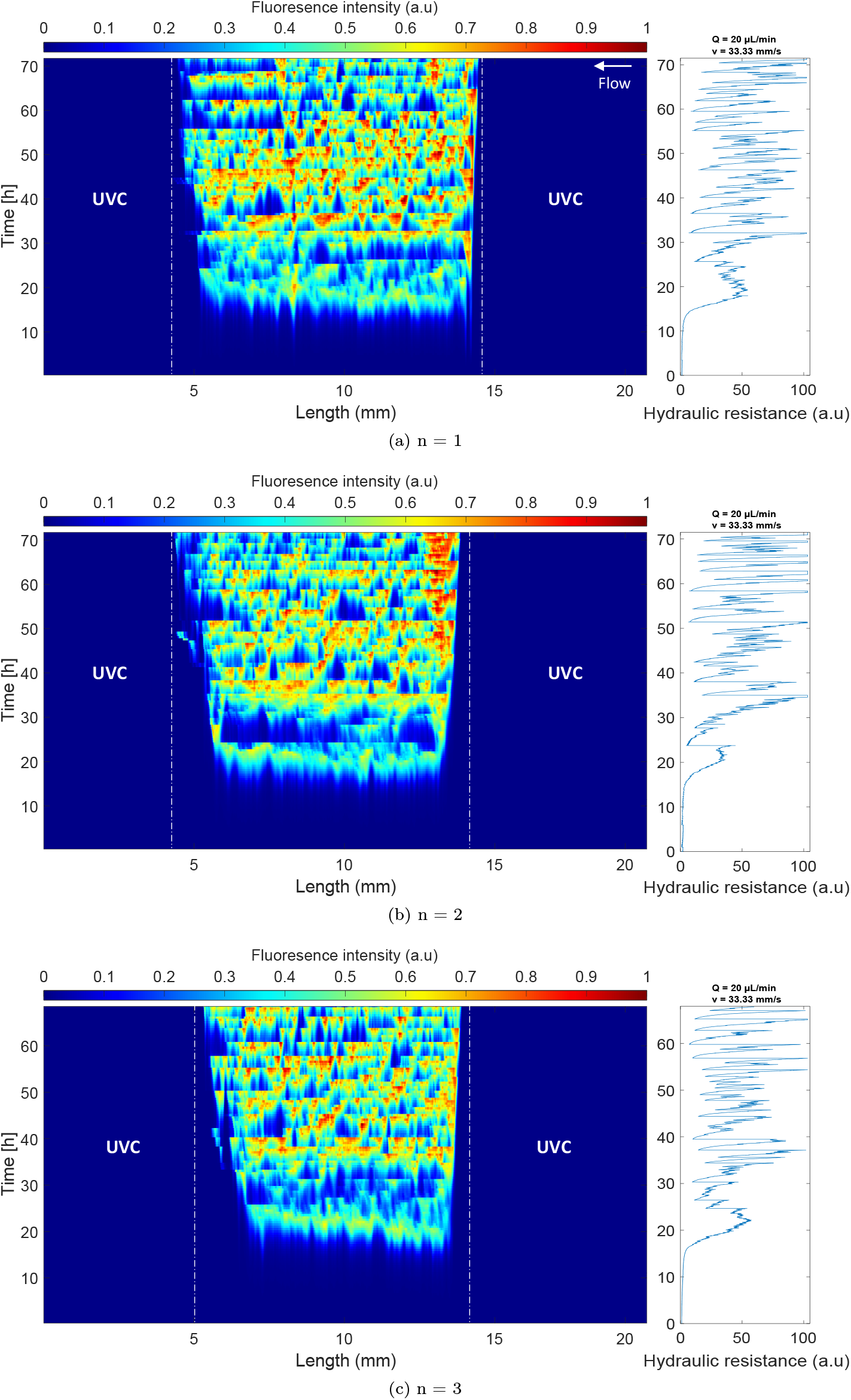
Kymograph representation for *Q* = 20 *µL/min* with [1X] BHI. Fluorescence intensity signal was normalized to the maximum signal intensity value. Plots on the right show the corresponding evolution of the hydraulic resistance.

**Figure S14.**
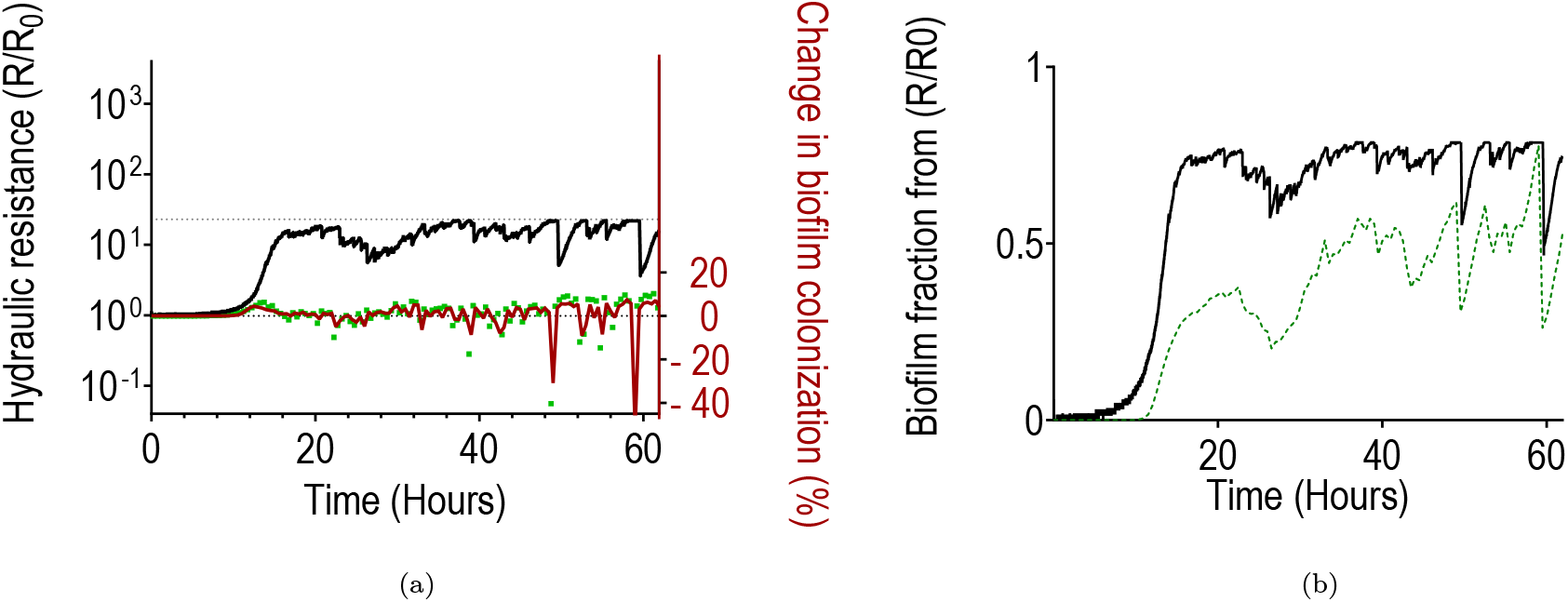
Temporal dynamics for *Q* = 200 *µL/min*. (a) evolution of the hydraulic resistance (black solid line) in time, calculated from pressure fluctuations. It also shows changes in biofilm colonization extracted from either integrated fluorescence intensity (green squares) or image segmentation (red solid line). (b): Fraction of biofilm in the microchannel, either calculated from hydraulic resistance (black solid line) or estimated from integrated GFP intensity (green dotted line), for the different flow rates.

**Figure S15.**
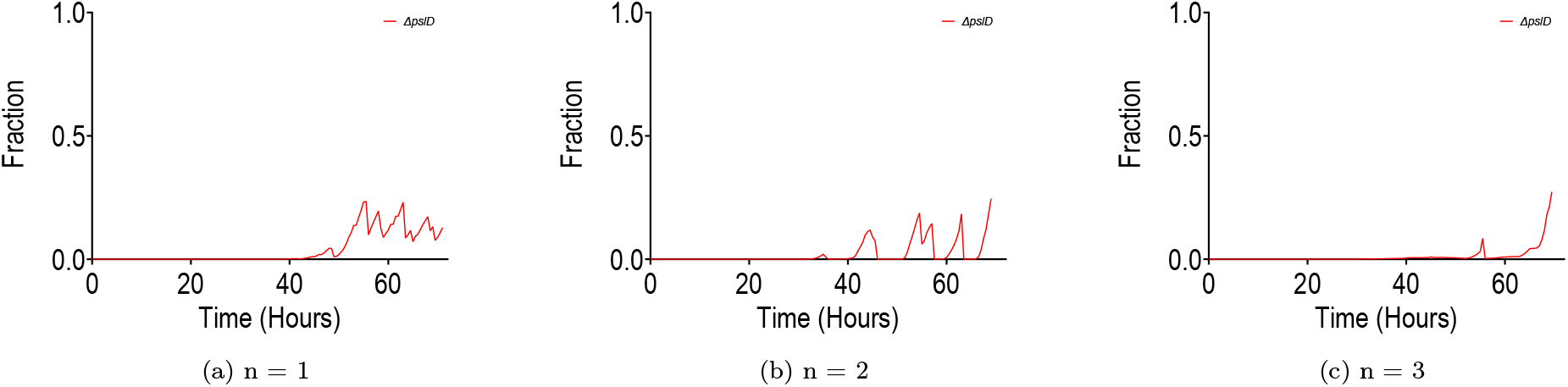
Temporal dynamics of growth and detachment for Δ*pslD* strain with *Q* = 0.2 *µL/min*. (a), (b) and (c): volume fraction calculated from image segmentation for three replicates.

**Figure S16.**
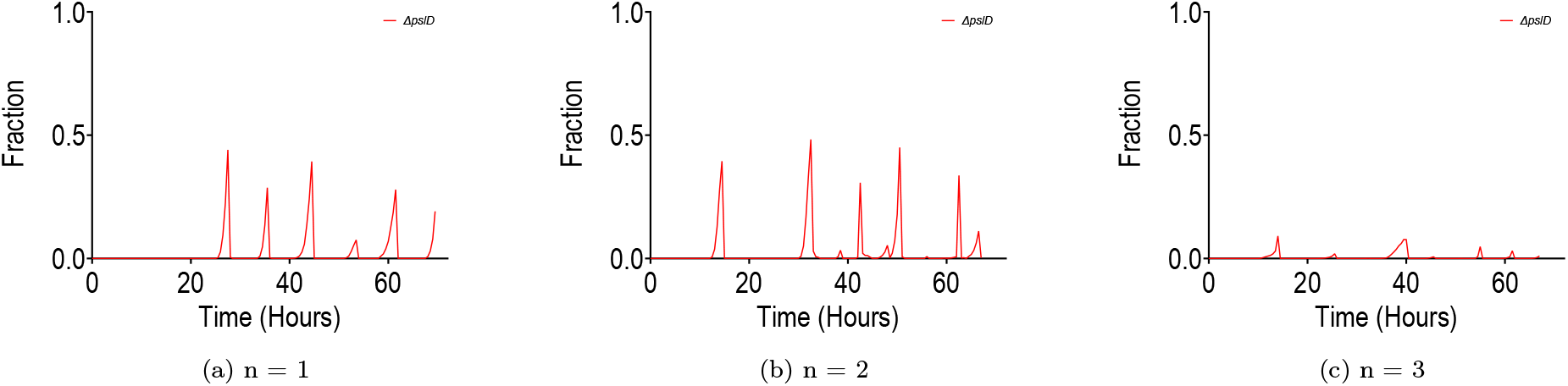
Temporal dynamics of growth and detachment for Δ*pslD* strain with *Q* = 2 *µL/min*. (a), (b) and (c): volume fraction calculated from image segmentation for three replicates.

**Figure S17.**
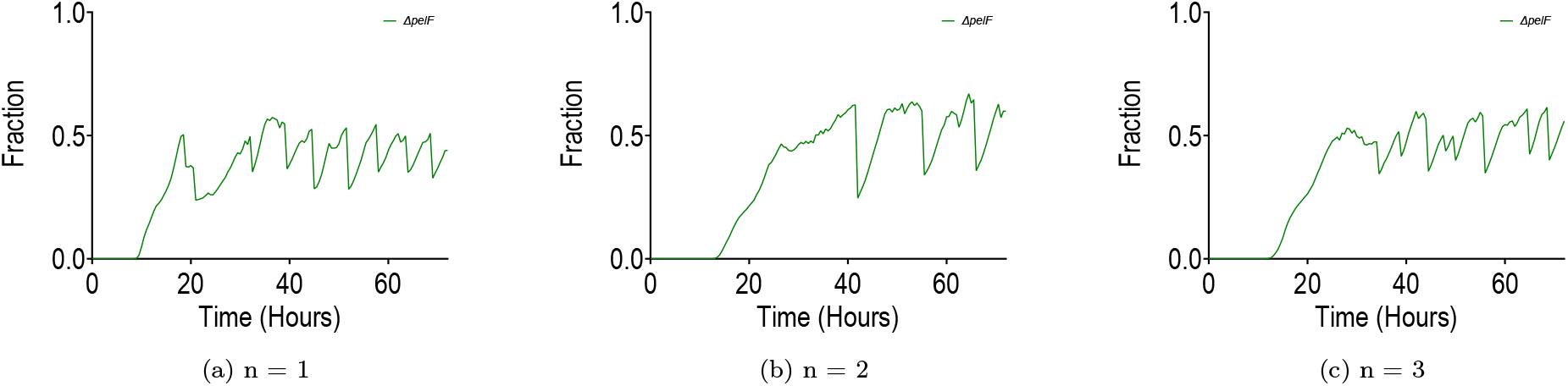
Temporal dynamics of growth and detachment for Δ*pelF* strain with *Q* = 0.2 *µL/min*. (a), (b) and (c): volume fraction calculated from image segmentation for three replicates.

**Figure S18.**
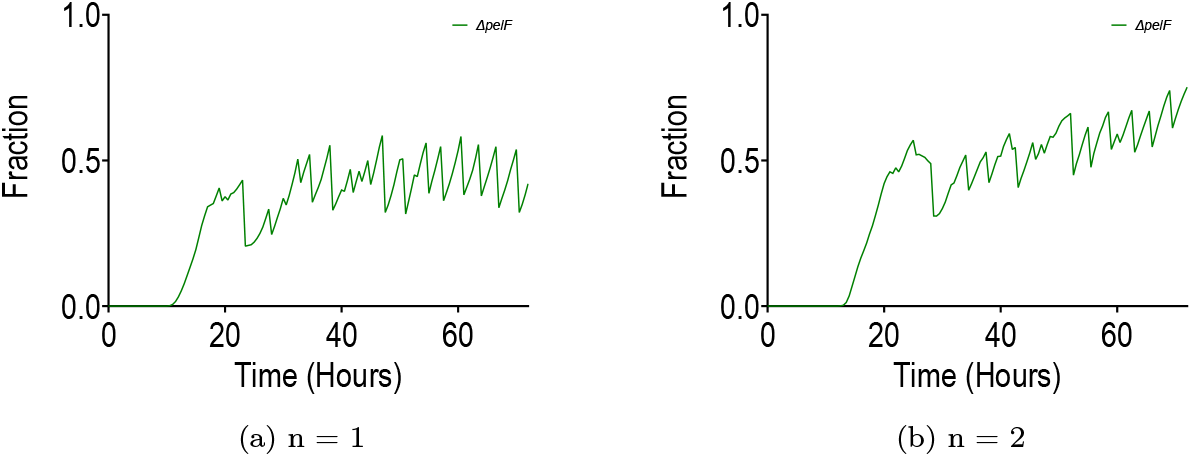
Temporal dynamics of growth and detachment for Δ*pelF* strain with *Q* = 2 *µL/min*. (a), (b) and (c): volume fraction calculated from image segmentation for two replicates.

**Figure S19.**
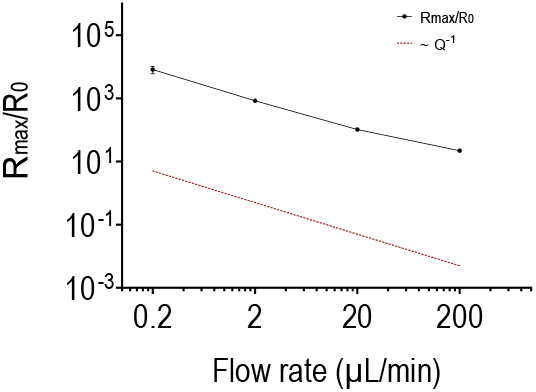
Log-log plot of 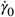 as a function of the flow rate (*Q* = 0.2 *µL/min*, 2 *µL/min*, 20 *µL/min* and 200 *µL/min*). The red dotted line simply shows the slope for an evolution with the inverse of the flow rate *Q*^−1^. Error bars represent standard deviation for n = 3 replicates, except for *Q* = 200 *µL/min*, for which n = 1.

**Figure S20.**
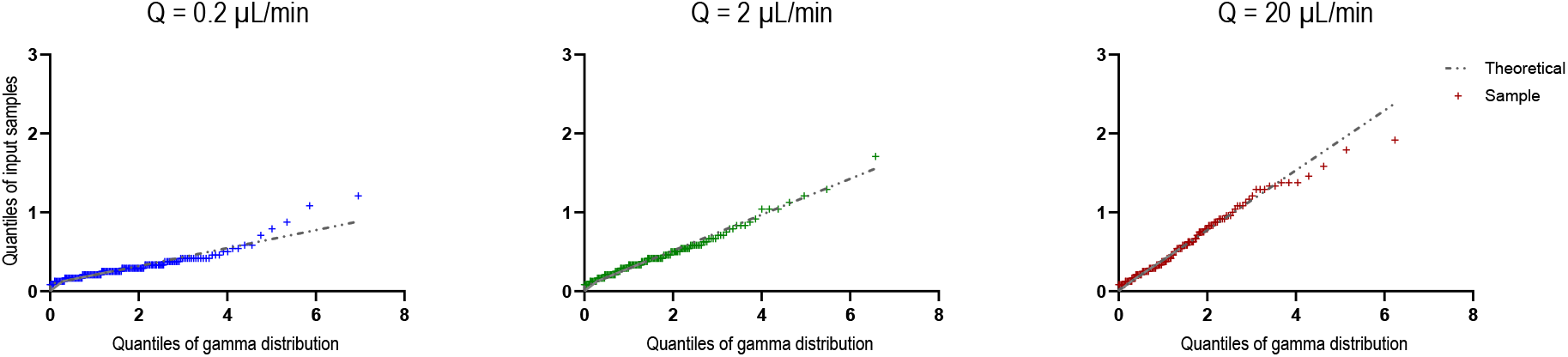
Quantile-quantile (Q-Q) plots of inter-event times fitted with Gamma distributions at three flow rates: *Q* = 0.2 *µL/min*, 2 *µL/min*, 20 *µL/min*. Sample quantiles (y-axis) are plotted against theoretical Gamma quantiles (x-axis). Close alignment with the diagonal reference line (red) indicates that the Gamma distribution provides a reasonable approximation of the overall distributional behavior.

**Figure S21.**
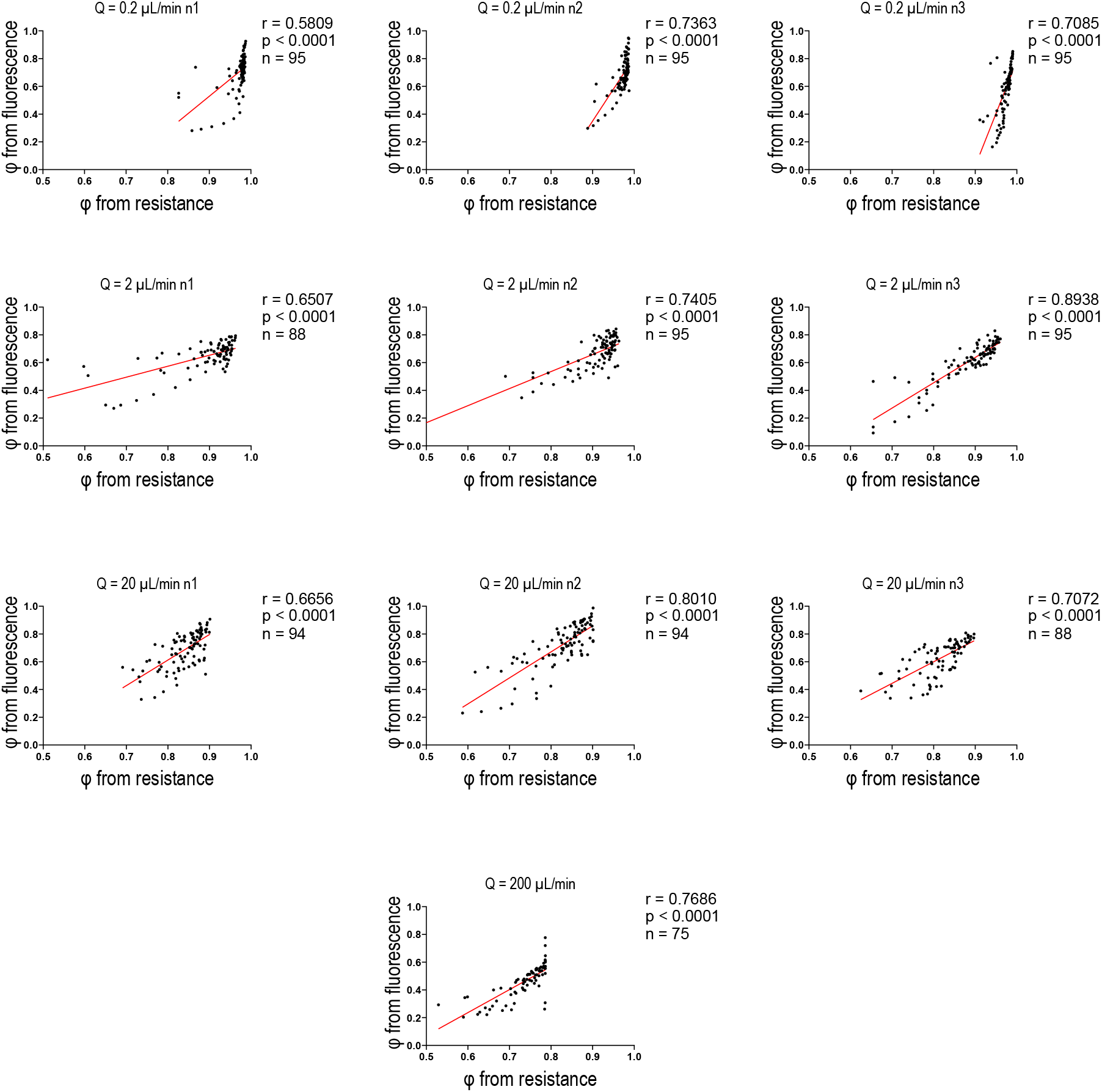
Pearson’s correlation between volume fractions obtained from image segmentation and calculated from hydraulic resistance for flow rates *Q* = 0.2 *µL/min*, 2 *µL/min*, 20 *µL/min* and 200 *µL/min*. To account for acquisition frequency mismatch, hydraulic resistance data were downsampled from 1 Hz to 1 point per 30 minutes. Each point represents one paired measurement. Data from all replicates were pooled for each condition (sample size n indicated on each plot). Red lines show linear regression fits: r = 0.68, 0.76, 0.72, 0.77 (mean across the 3 replicates) for *Q* = 0.2 *µL/min*, 2 *µL/min*, 20 *µL/min*, 200 *µL/min* respectively; p < 0.0001 for all conditions.

**Table S1.**
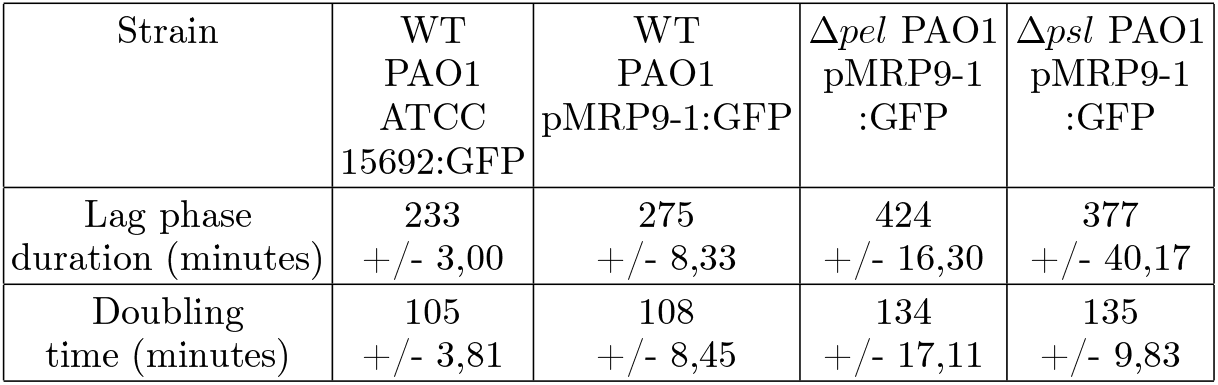
Lag phase and doubling time in minutes for *P. aeruginosa* WT PAO1 and EPS defective strains. Mean values were obtained from 3 replicates for each strain and +/− indicates the standard deviation.

## III. SUPPLEMENTARY MOVIES

- Movie S1: DIC timelapse of the initial development at 0.2 *µL/min*.
- Movie S2: DIC timelapse of the initial development at 2 *µL/min*.
- Movie S3: DIC timelapse of the initial development at 20 *µL/min*.
- Movie S4: composite (GFP and brightfield) timelapse of biofilm development at 20 *µL/min*.

