## Supplementary material for "Should I stay or should I go? Spatiotemporal dynamics of bacterial biofilms in confined flows": Main supplementary file

### Supplementary information

#### I. MODELS

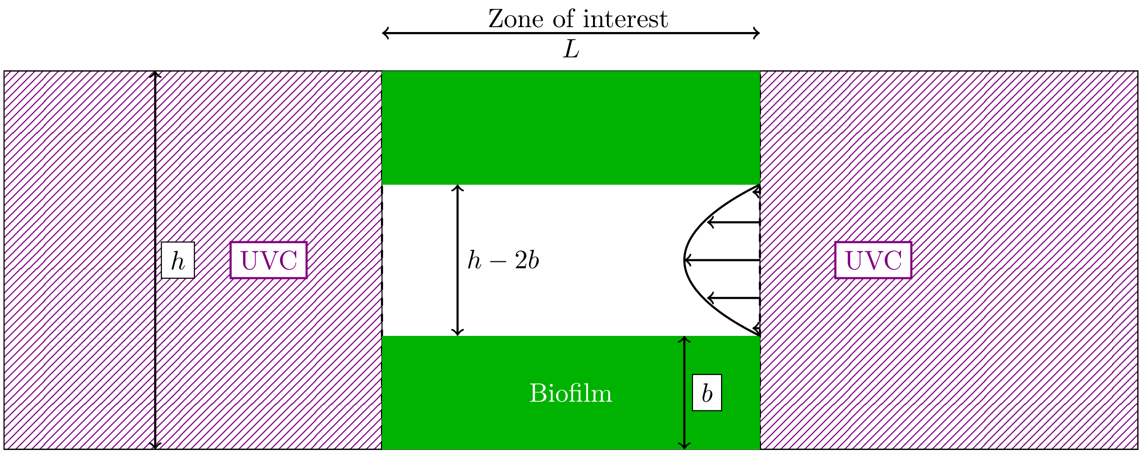


##### 1. Nutrient transport

The limiting nutrient is treated as a solute being transported by advection/diffusion and consumed by bacteria. We consider a cross-section average concentration of the form $C = \int_{A(x)} c(x, y, z)\, dydz$ with $A$ the section at any given $x$ – $x$ being the longitudinal direction. The transport reads

$$\partial_t C + \underbrace{V\partial_x C}_{\text{Advection}} = \underbrace{D\partial_{xx} C}_{\text{Diffusion}} - \underbrace{\phi\alpha\frac{C}{C+K}}_{\text{Nutrient uptake}}, \quad (1)$$

where $C$ $\left[kg \times m^{-3}\right]$ is the concentration, $x$ is the longitudinal coordinate system, $\phi$ is the cross-section volume fraction of biofilm, $V$ $\left[m \times s^{-1}\right]$ the average velocity in the empty channel, $D = 10^{-9} m^2 \times s^{-1}$ an estimate diffusion coefficient of the solute, $\alpha$ $\left[kg \times m^{-3} \times s^{-1}\right]$ the uptake rate and $K$ $\left[kg \times m^{-3}\right]$ the half-saturation constant. As we impose a constant flow rate for each experiment, the velocity averaged over the entire cross-section remains constant, even for different values of $\phi$. The term $\phi$ in the nutrient uptake indicates that consumption is proportional to the volume fraction of biofilm.

We considered that nutrient transport is quasi-steady, since solute transport is much faster than growth. We further nondimensionalized this equation as

$$\boxed{Pe\partial_x C^\star = \partial_{xx} C^\star - Da\phi\frac{C^\star}{C^\star + K^\star}} \quad (2)$$

with $C^\star = \frac{C}{C_0}$ with the inlet concentration $C_0$ $\left[kg \times m^{-3}\right]$, the length of the channel $L = 10\ mm$, the Péclet number $Pe = \frac{VL}{D}$ and $Da$ the Damköhler number defined as the ratio of diffusive to reactive times, $Da = \frac{\alpha L^2}{C_0 D}$.

With these hypotheses, we write the pressure drop across the channel as

$$\Delta P = R\left(\phi\right)Q, \tag{3}$$

with $\Delta P$ the pressure difference, $Q$ the flowrate and $R$ the hydraulic resistance (which is a function of the volume fraction of biofilm). The hydraulic resistance in the empty channel is

$$R_0 = A\frac{\mu L}{h^4} \tag{4}$$

with $A = \frac{12}{1-\sum_{n,\mathrm{odd}}^{\infty}\frac{1}{n^5}\frac{192}{\pi^5}\tanh\left(n\frac{\pi}{2}\right)}$ – which can be approximated as $R_0 = \frac{12\mu L}{h^4[1-0.917\times0.63]}$ with $A \simeq \frac{12}{1-0.917\times0.63}$. When the biofilm grows as a uniform layer, we also have

$$R = A\frac{\mu L}{\left(h-2b\right)^4} \tag{5}$$

To obtain an expression in terms of the volume fraction of biofilm, we use $\phi = 1 - \left(1 - 2\frac{b}{h}\right)^2$ and $b = h\frac{1-\sqrt{1-\phi}}{2}$ – and also $h - 2b = h\left(1-\phi\right)^{1/2}$. We thus have

$$\boxed{\frac{R}{R_0} = \frac{1}{\left(1-\phi\right)^2}}. \tag{6}$$

**Reference**: Kurz, D. L.; Secchi, E.; Stocker, R.; Jimenez-Martinez, J. Morphogenesis of biofilms in porous media and control on hydrodynamics. Environ. Sci. Technol. 2023, 57 (14), 5666—5677.

*3. Hydrodynamic stresses*

We first write $\sigma_0$ the shear stress on the liquid/solid surface of the channel when $\phi = 0$. To obtain an explicit expression for this stress, the easiest way to proceed is to notice that, in the $x$ direction, tangential to the surface of the channel, the force exerted by the pressure difference between the inlet and outlet is equal to the viscous force exerted by the fluid on the solid. This can be easily shown by integrating $\nabla \cdot \boldsymbol{\sigma} = 0$, with $\boldsymbol{\sigma} = -p\mathbf{I} + \mu\left(\nabla\mathbf{v} +^{\mathsf{T}} \nabla\mathbf{v}\right)$ and $p$ the pressure and $\mu$ the viscosity, over a control volume that is the the zone of interest between UVCs in the channel. With $4hL$ the solid surface in the zone of interest, we have $\sigma_0 4hL = h^2\Delta P$ where $L$ is the length of the control volume, $h$ is the width of the square channel and $\Delta P$ is the pressure difference between inlet and outlet. We therefore have

$$\sigma_0 = \frac{h\Delta P}{4L}. \tag{7}$$

We can also write $\Delta P = R_0 Q$ with the hydraulic resistance in the square cross-section channel, $R_0 = A\frac{\mu L}{h^4}$. Therefore, we have

$$\sigma_0 = \frac{A}{4}\frac{\mu Q}{h^3}. \tag{8}$$

With the same type of reasoning, the total force along $x$ applied by the fluid on the biofilm, including both shear and pressure is $h^2\Delta P = A\frac{\mu L Q}{h^2(1-\phi)^2}$. Therefore, the total stress on the biofilm solid surface is

$$\frac{\sigma}{\sigma_0} = \frac{1}{\left(1-\phi\right)^2}, \tag{9}$$

with a contribution from pressure

$$\frac{\sigma_{\text{pressure}}}{\sigma_0} = \frac{\phi}{\left(1-\phi\right)^2}, \tag{10}$$

and from shear

$$\frac{\sigma_{\text{shear}}}{\sigma_0} = \frac{1-\phi}{\left(1-\phi\right)^2}. \tag{11}$$

Also note that the shear stress at the biofilm fluid interface is

$$\frac{\tau_{\text{shear}}}{\sigma_0} = \frac{1}{\left(1-\phi\right)^{3/2}}, \tag{12}$$

with the difference being simply the ratio of biofilm/solid and biofilm/fluid surfaces.

#### *4. Biofilm growth*

We model biofilm development as

$$\partial_t \phi = \phi \left( \frac{Y\alpha}{X} \frac{C}{C+K} - \chi\sqrt{\sigma} \right) \tag{13}$$

with $Y$ a yield coefficient (mass of biofilm produced by mass of limiting nutrient consumed), $X$ the density of the biofilm $X$ $\left[kg \times m^{-3}\right]$ and $\chi$ the sensitivity to biofilm detachment $\chi$ $\left[\sqrt{kg^{-1} \times m}\right]$. We non-dimensionalize with time $\tau = \frac{X}{Y\alpha}\left(1+K^\star\right)$ in order to obtain

$$\partial_t \phi = \phi \left[ \frac{C^\star\left(1+K^\star\right)}{C^\star+K^\star} - \frac{X}{Y\alpha}\left(1+K^\star\right)\chi\sqrt{\sigma_0}\sqrt{\frac{\sigma}{\sigma_0}} \right]. \tag{14}$$

Recalling that $\sqrt{\frac{\sigma}{\sigma_0}} = (1-\phi)^{-1}$ and writing the dimensionless number $\mathcal{M} = \frac{X}{Y\alpha}\left(1+K^\star\right)\chi\sqrt{\sigma_0}$, we obtain

$$\partial_t \phi = \phi \left[ \frac{C^\star\left(1+K^\star\right)}{C^\star+K^\star} - \frac{\mathcal{M}}{1-\phi} \right]. \tag{15}$$

Assuming that $\mathcal{M} \leq 1$, which is consistent with the observation that biofilm is never completely removed from the channels, we can also write this as

$$\partial_t \phi = \phi \left[ \frac{C^\star\left(1+K^\star\right)}{C^\star+K^\star} - \frac{1-\phi_{\max}}{1-\phi} \right] \tag{16}$$

with $\phi_{\max}$ the maximum value of $\phi$.
In the case with no nutrient limitation, we assume $C^\star = 1$ everywhere in the channel, so that we have

$$\dot{\phi} = \phi \left( 1 - \frac{1-\phi_{\max}}{1-\phi} \right), \tag{17}$$

with $\phi$ describing an average volume fraction in the entire channel, and thus being only a function of time.

We can also describe sloughing by modifying this equation as

$$d\phi_t = \phi_t \left( 1 - \frac{1-\phi_{\max}}{1-\phi_t} \right) dt - \phi_{t^-} dN_t, \tag{18}$$

with $\phi_t$ describing the average fraction of biofilm in the microchannel, $N$ the jumps and $\phi_{t^-}$ the value of $\phi$ at time $t^-$ just before the jump.

#### II. SUPPLEMENTARY FIGURES

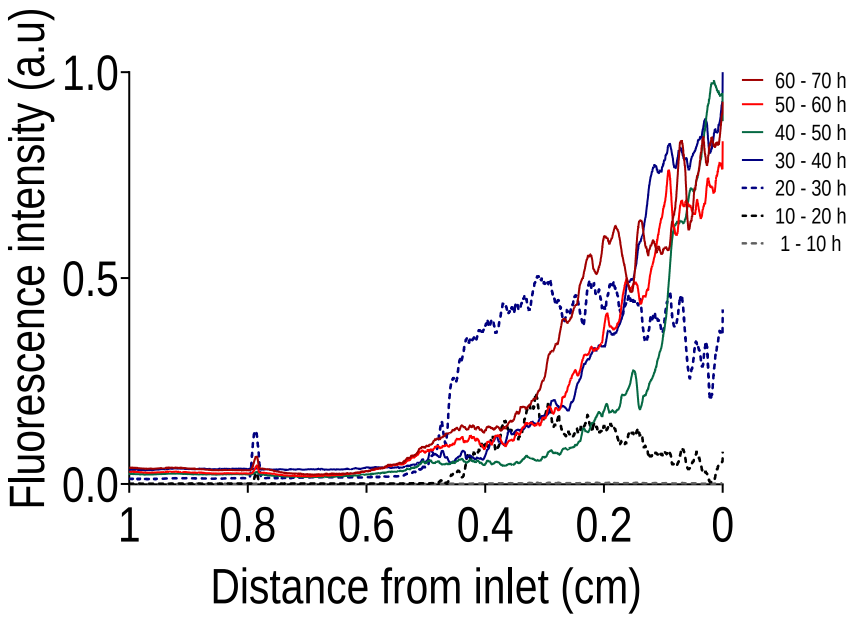


Figure 1: Time evolution (10 hours by curve) for $Q = 0.02\ \mu L/min$ of the averaged (n = 3) longitudinal distribution of the biofilm and the impact of nutrient limitation.

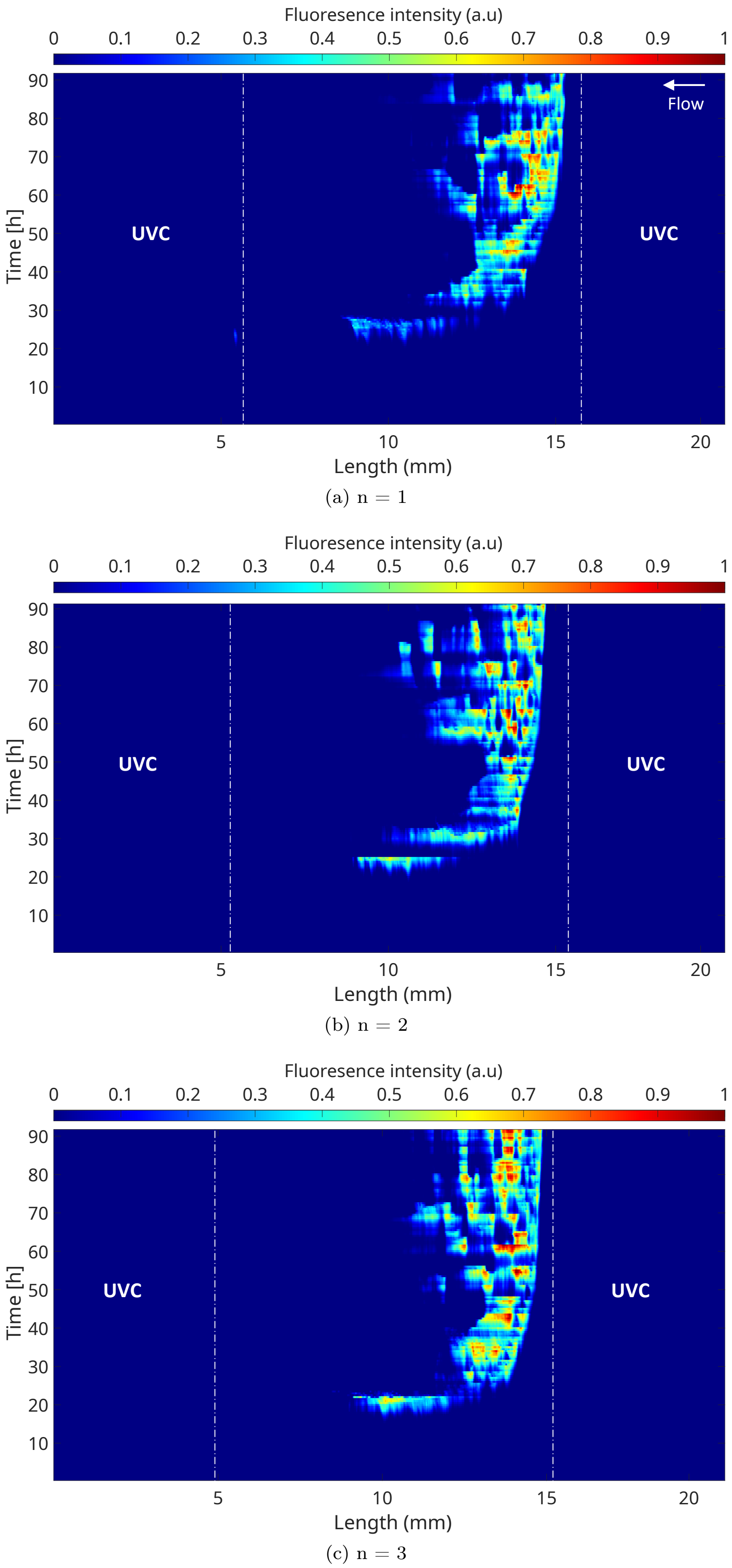


Figure 2: Kymograph representations based on the GFP signal for $Q = 0.02\ \mu L/min$ with [1X] BHI showing the spatio-temporal dynamics of the biofilm longitudinal distribution. Fluorescence intensity signal was normalized to the maximum signal intensity value.

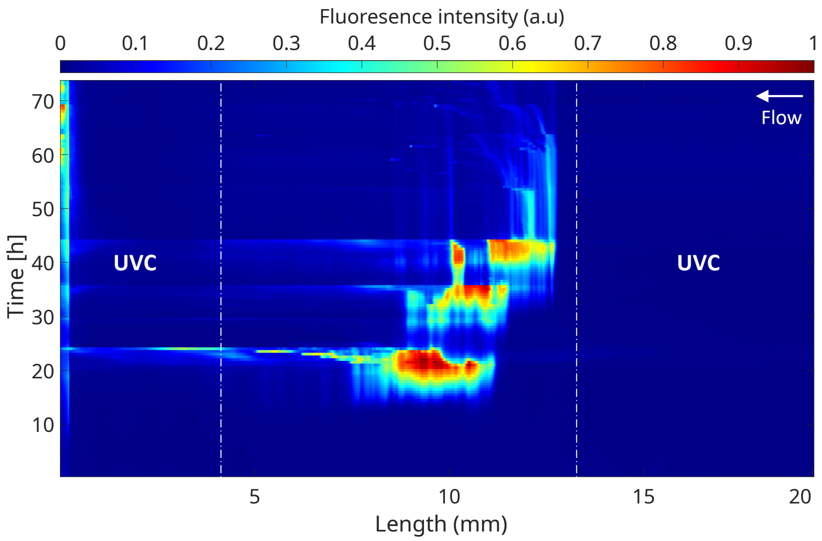


(a) n = 1

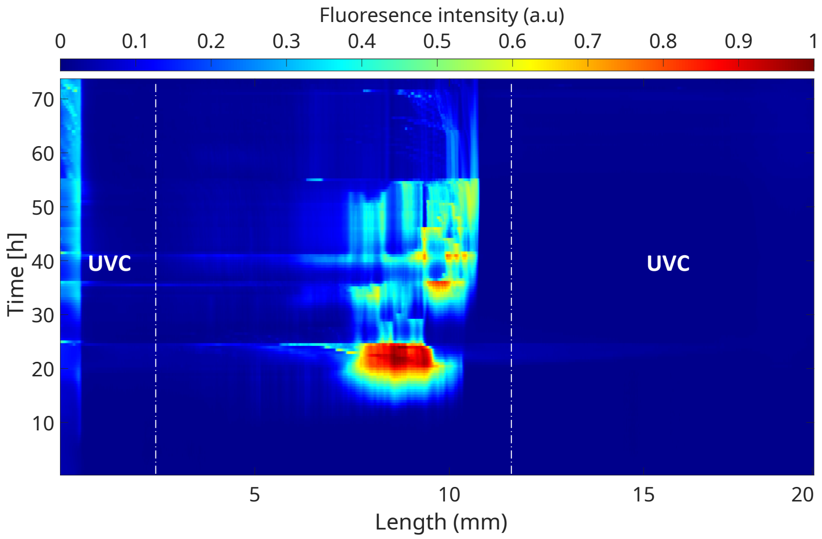


(b) n = 2

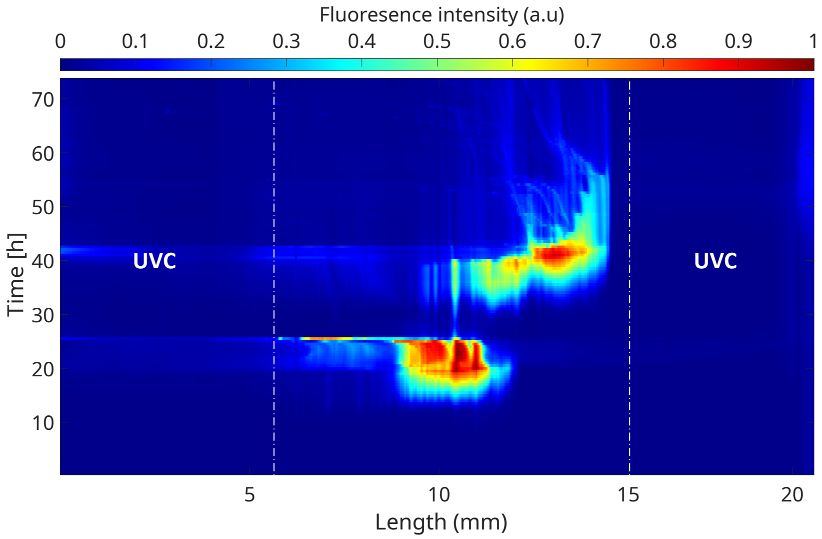


(c) n = 3

Figure 3: Kymograph representation for $Q = 0.02\ \mu L/min$ with [1X] BHI supplemented by 8 $g/L$ glucose. Fluorescence intensity signal was normalized to the maximum signal intensity value.

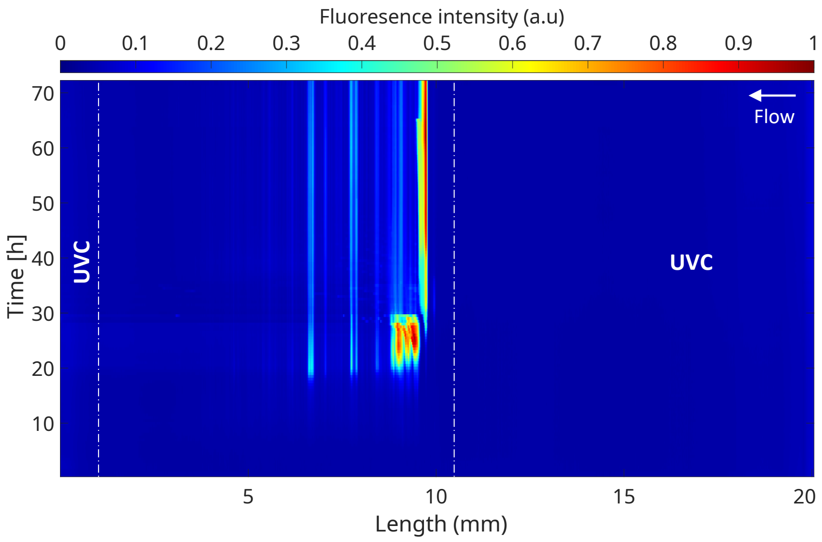


(a) n = 1

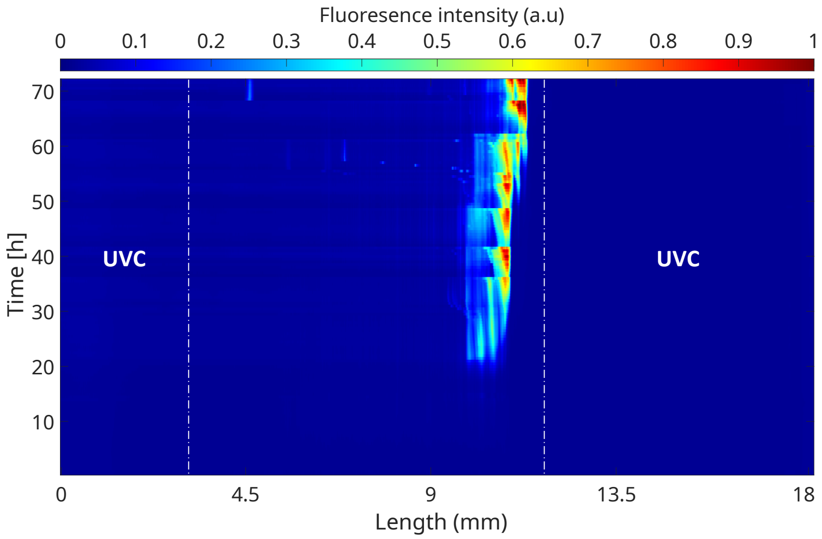


(b) n = 2

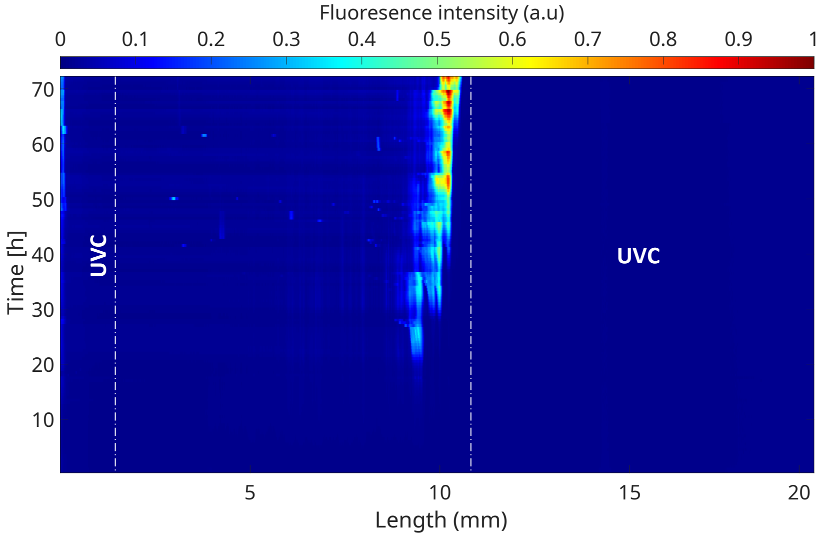


(c) n = 3

Figure 4: Kymograph representation for $Q = 0.02\ \mu L/min$ with [0.2X] BHI. Fluorescence intensity signal was normalized to the maximum signal intensity value.

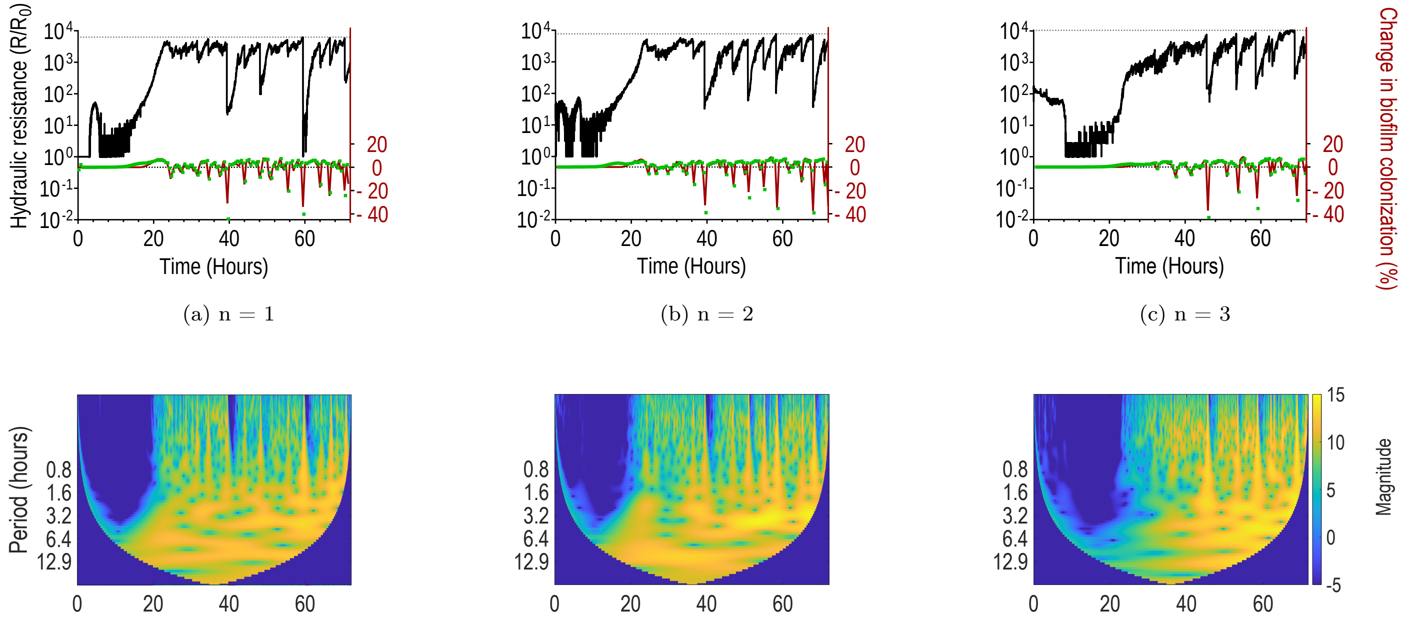


(a) n = 1 (b) n = 2 (c) n = 3

(d) n = 1 (e) n = 2 (f) n = 3

Figure 5: **Temporal dynamics of growth and detachment for** $Q = 0.2\ \mu L/min$**.** (a), (b) and (c) show the variation of hydraulic resistance (black line) for three replicates, along with the percentage of colonization extracted from image segmentation (red line) and the integrated fluorescence intensity signal (green). Thin dotted lines on the top of y axis indicate the maximum value reached by hydraulic resistance. (d), (e) and (f): corresponding wavelet scalograms for the pressure signal.

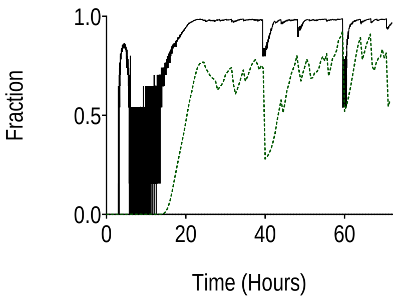


(a) n = 1

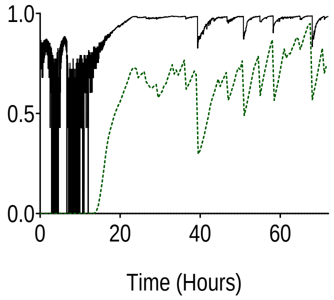


(b) n = 2

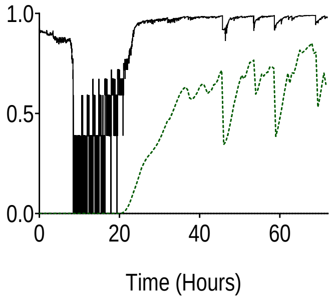


(c) n = 3

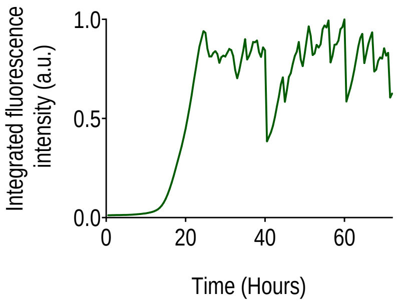


(d) n = 1

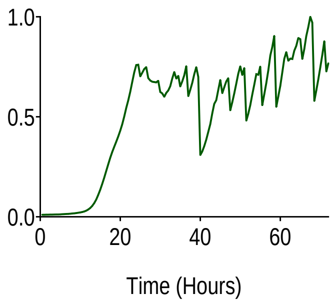


(e) n = 2

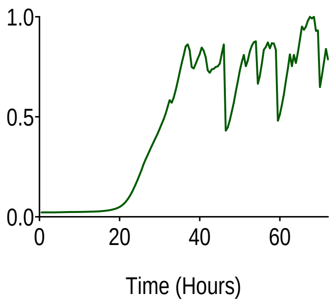


(f) n = 3

Figure 6: **Temporal dynamics of growth and detachment for $Q = 0.2\ \mu L/min$.** (a), (b) and (c) show the evolution of the volume fraction as a function of time, with that calculated from hydraulic resistance (black line) and from segmentation (green dotted line). (d), (e) and (f): normalized integrated fluorescence intensity signal as a function of time.

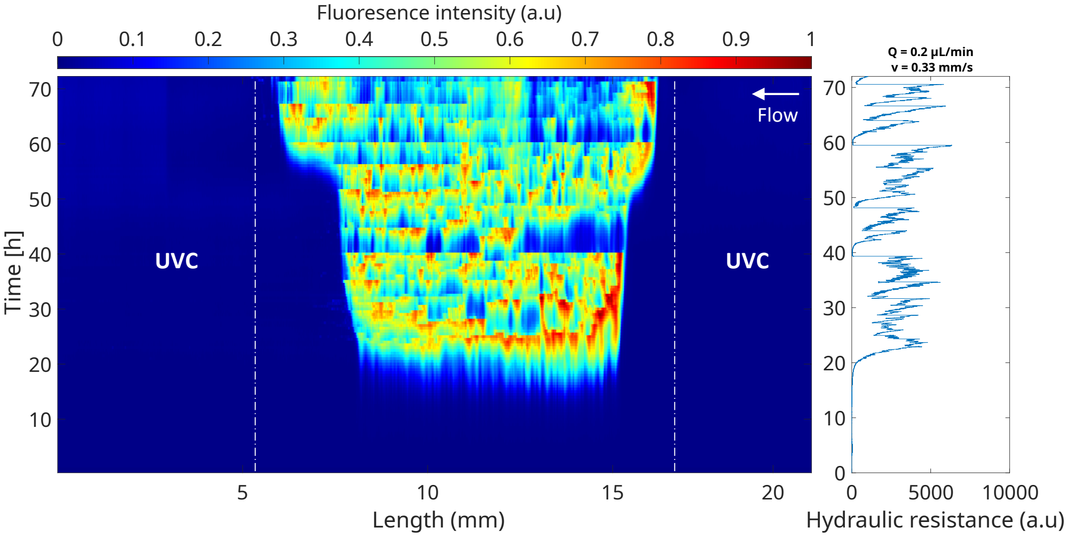


(a) n = 1

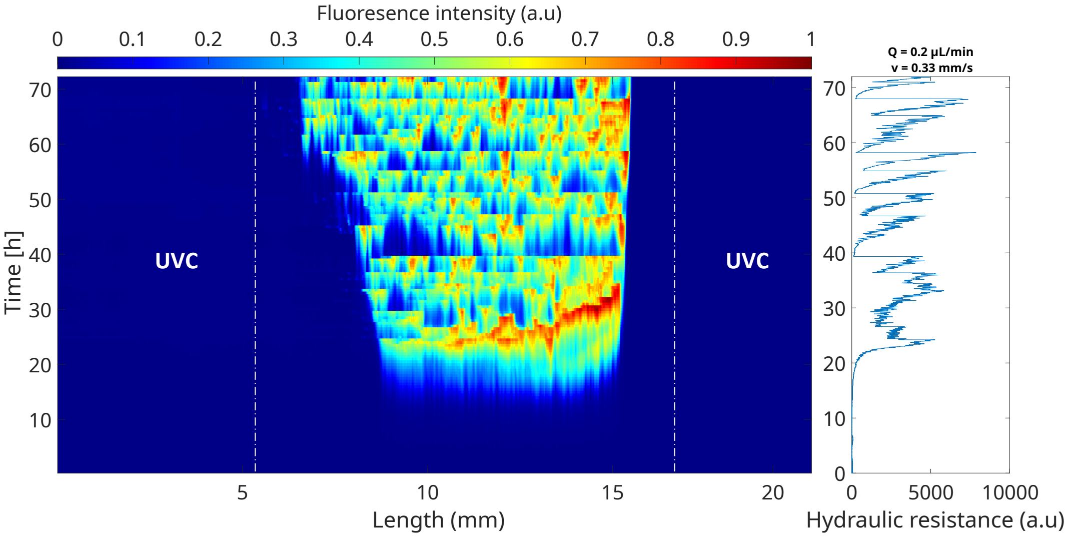


(b) n = 2

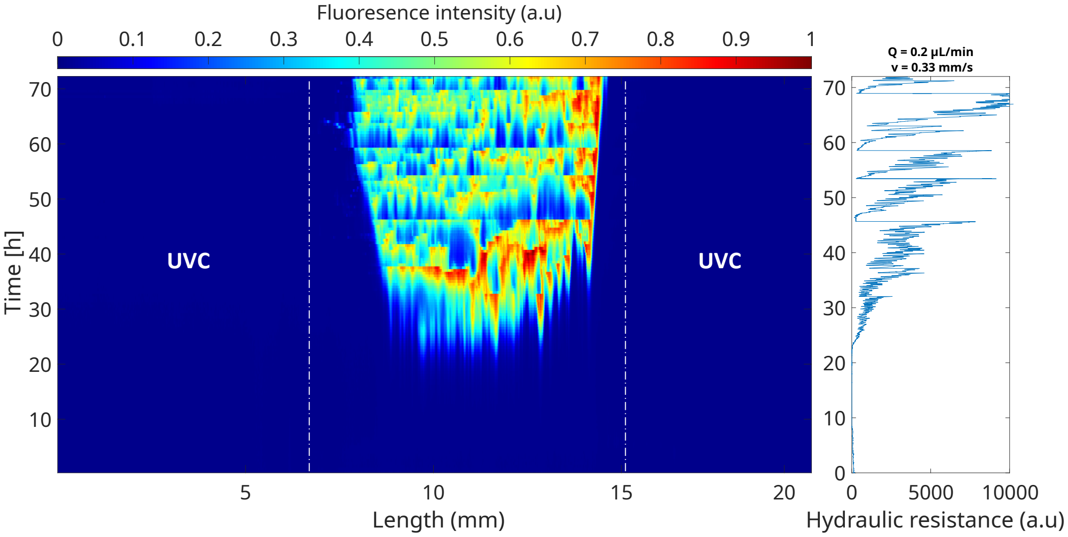


(c) n = 3

Figure 7: Kymograph representation for $Q = 0.2\ \mu L/min$ with [1X] BHI. Fluorescence intensity signal was normalized to the maximum signal intensity value. Plots on the right show the corresponding evolution of the hydraulic resistance.

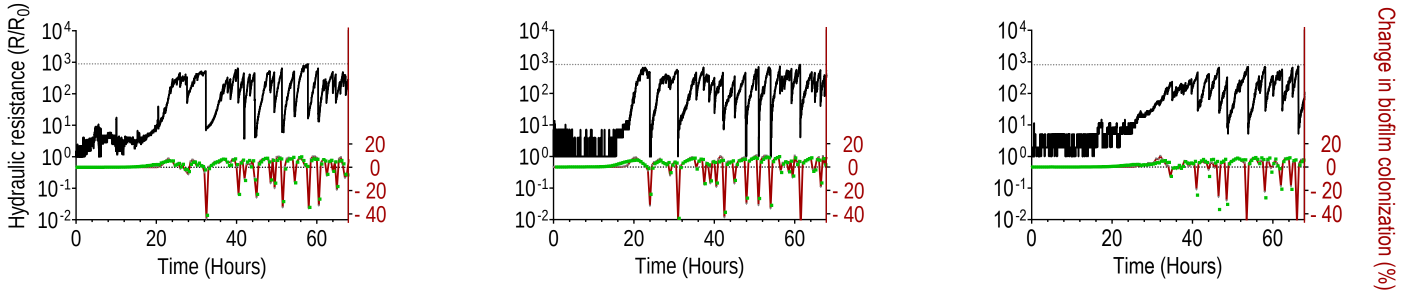


(a) n = 1 (b) n = 2 (c) n = 3

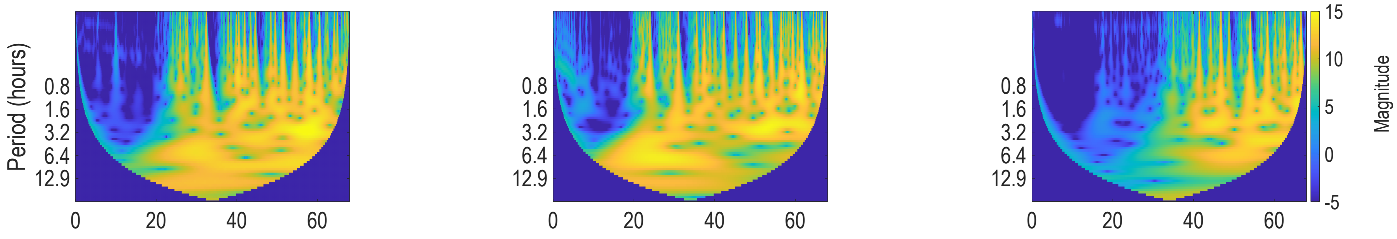


(d) n = 1 (e) n = 2 (f) n = 3

Figure 8: **Temporal dynamics of growth and detachment for $Q = 2\ \mu L/min$.** (a), (b) and (c) show the variation of hydraulic resistance (black line) for three replicates, along with the percentage of colonization extracted from image segmentation (red line) and the integrated fluorescence intensity signal (green). Thin dotted lines on the top of y axis indicate the maximum value reached by hydraulic resistance. (d), (e) and (f): corresponding wavelet scalograms for the pressure signal.

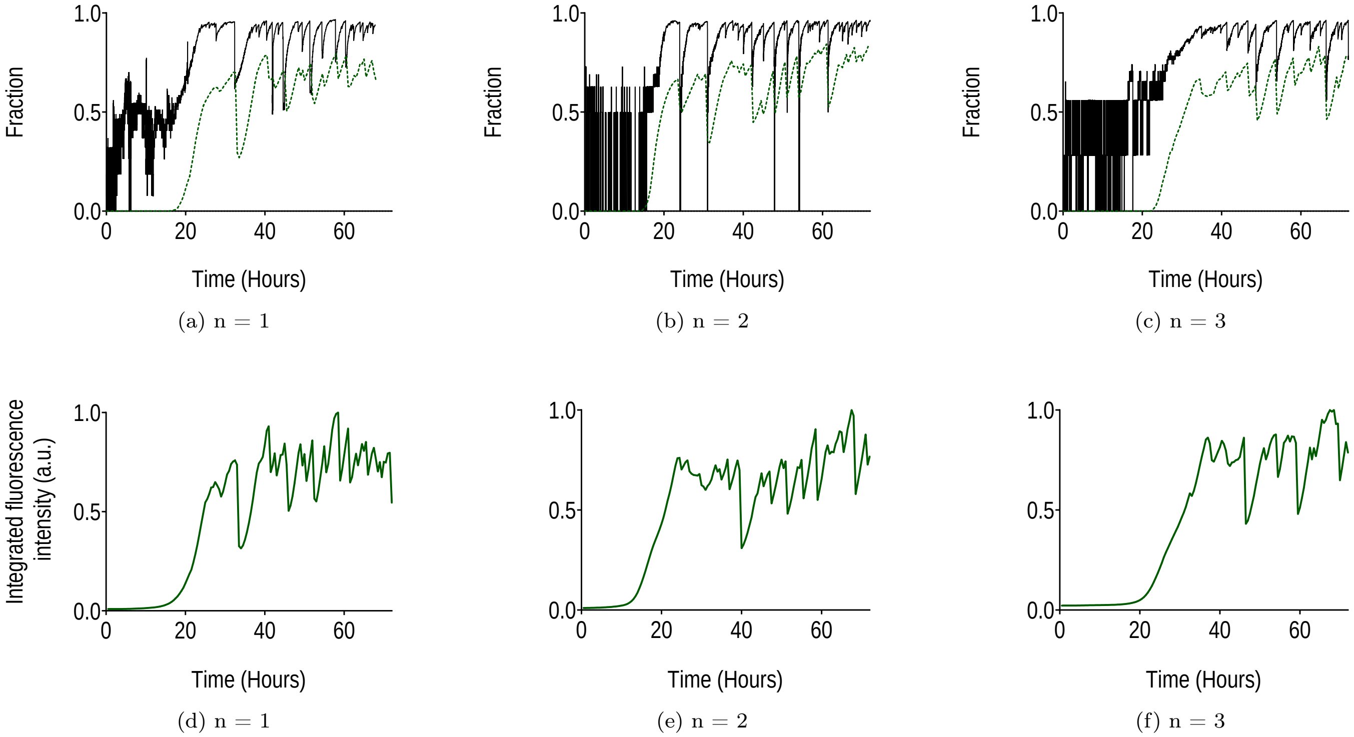


(a) n = 1 (b) n = 2 (c) n = 3

(d) n = 1 (e) n = 2 (f) n = 3

Figure 9: **Temporal dynamics of growth and detachment for $Q = 2\ \mu L/min$.** (a), (b) and (c) show the evolution of the volume fraction as a function of time, with that calculated from hydraulic resistance (black line) and from segmentation (green dotted line). (d), (e) and (f): normalized integrated fluorescence intensity signal as a function of time.

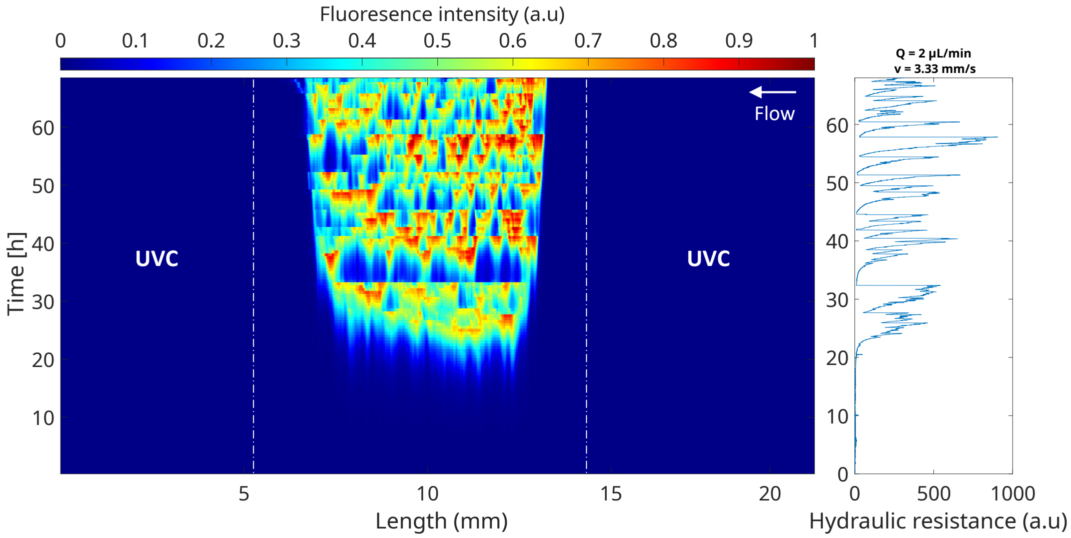


(a) n = 1

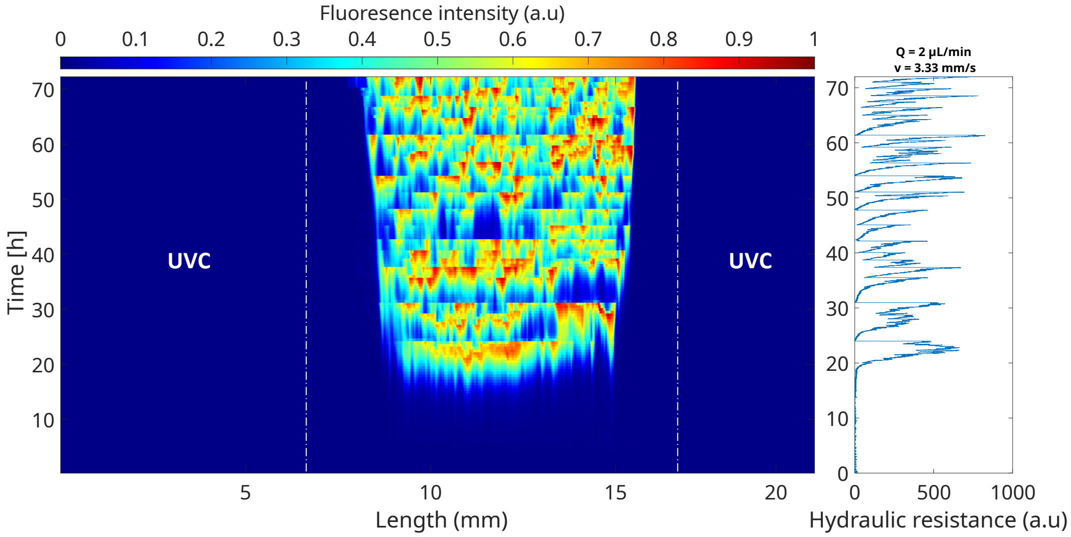


(b) n = 2

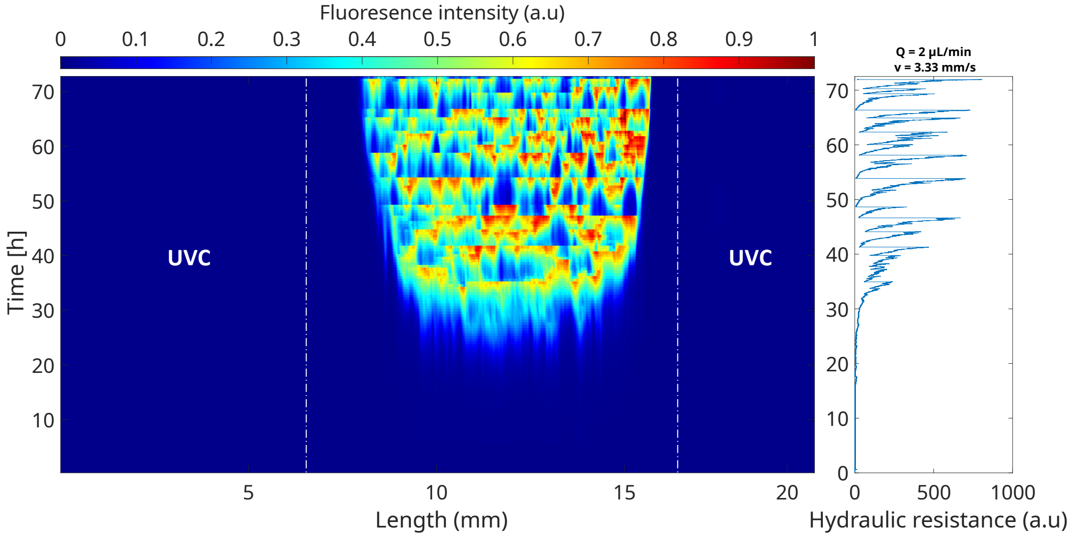


(c) n = 3

Figure 10: Kymograph representation for $Q = 2\ \mu L/min$ with [1X] BHI. Fluorescence intensity signal was normalized to the maximum signal intensity value. Plots on the right show the corresponding evolution of the hydraulic resistance.

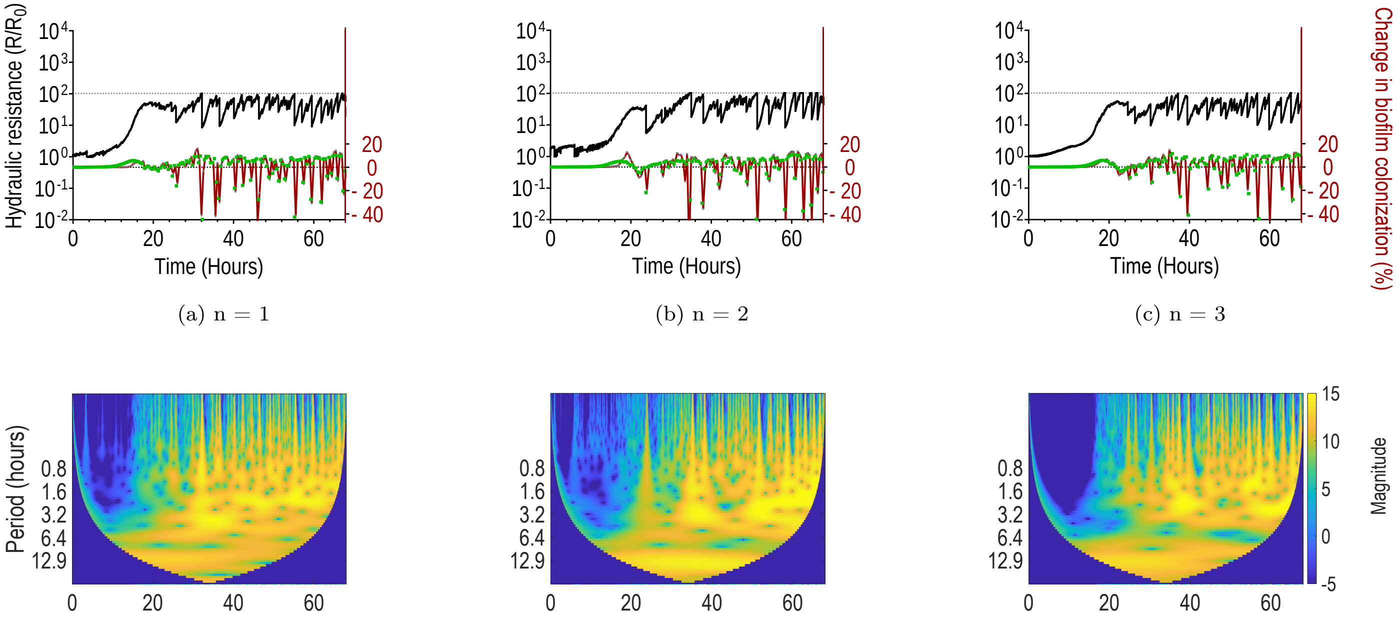


(a) n = 1 (b) n = 2 (c) n = 3

(d) n = 1 (e) n = 2 (f) n = 3

Figure 11: **Temporal dynamics of growth and detachment for** $Q = 20\ \mu L/min$**.** (a), (b) and (c) show the variation of hydraulic resistance (black line) for three replicates, along with the percentage of colonization extracted from image segmentation (red line) and the integrated fluorescence intensity signal (green). Thin dotted lines on the top of y axis indicate the maximum value reached by hydraulic resistance. (d), (e) and (f): corresponding wavelet scalograms for the pressure signal.

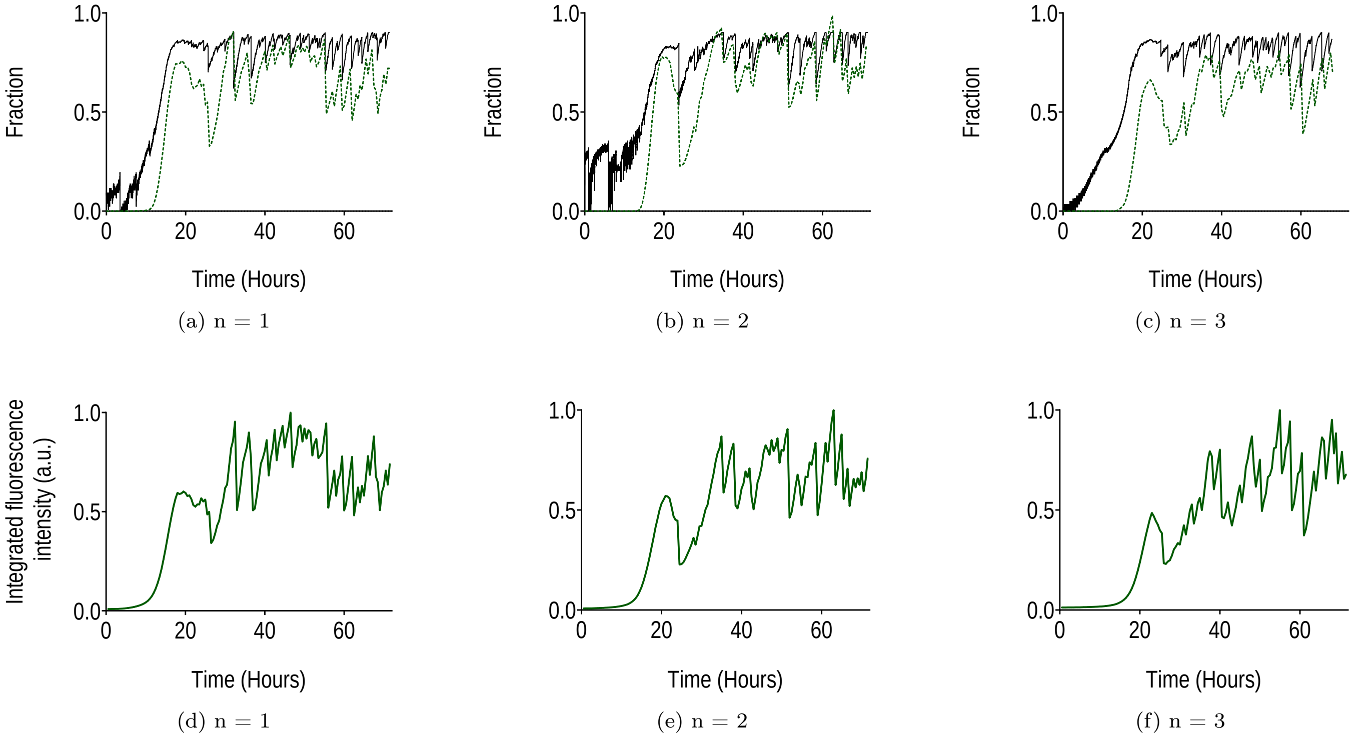


Figure 12: **Temporal dynamics of growth and detachment for** $Q = 20\ \mu L/min$**.** (a), (b) and (c) show the evolution of the volume fraction as a function of time, with that calculated from hydraulic resistance (black line) and from segmentation (green dotted line). (d), (e) and (f): normalized integrated fluorescence intensity signal as a function of time.

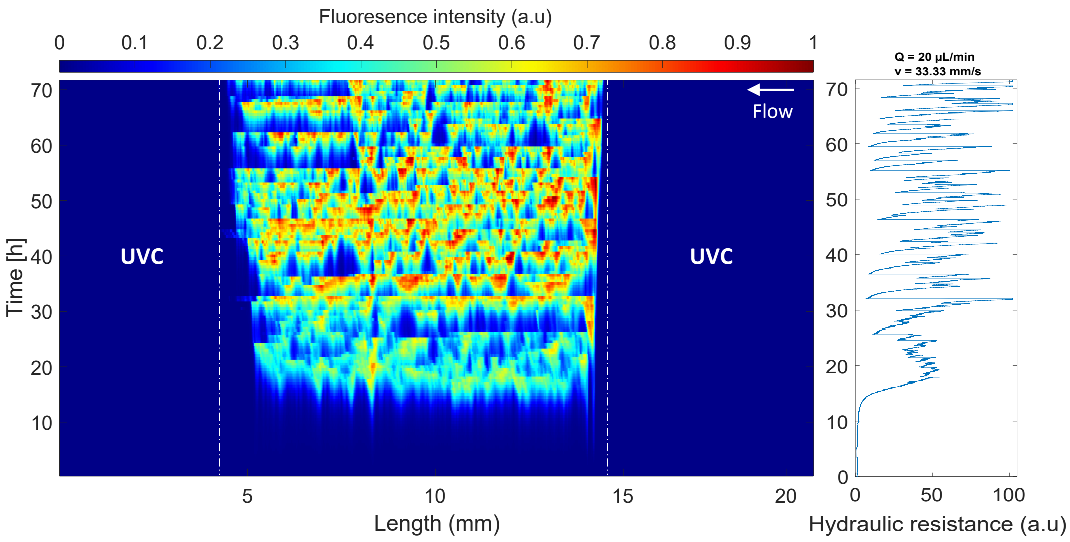


(a) n = 1

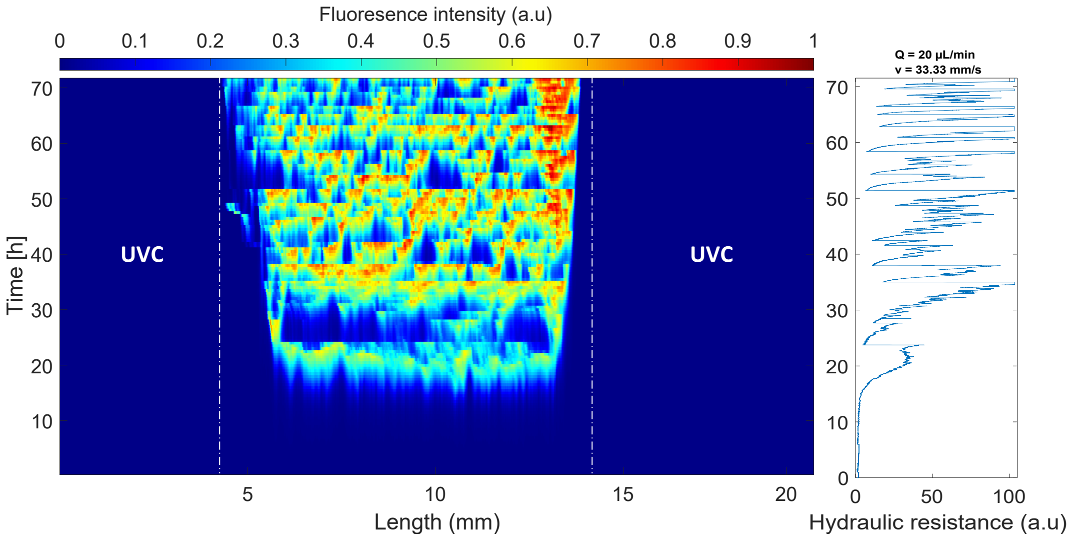


(b) n = 2

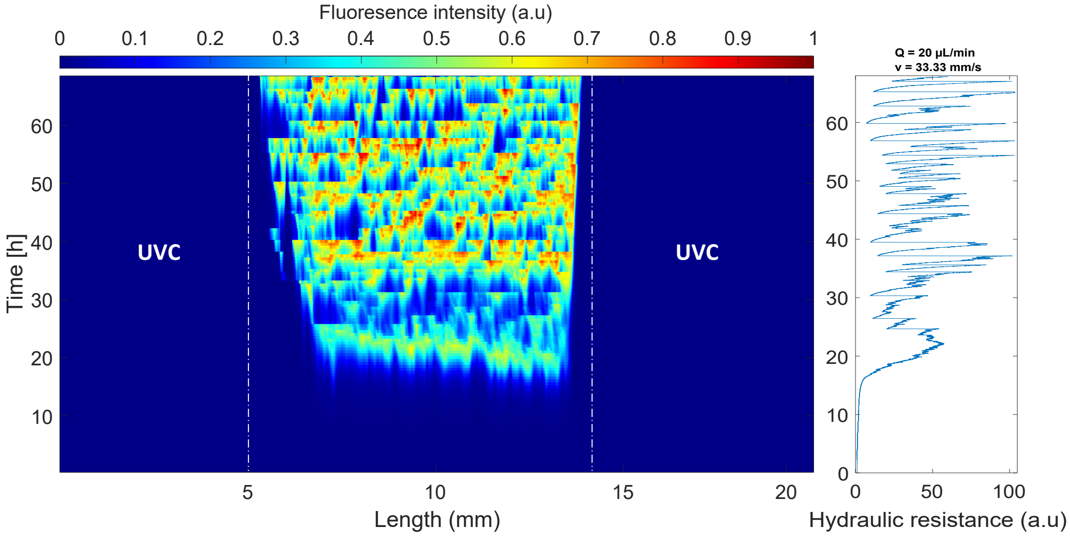


(c) n = 3

Figure 13: Kymograph representation for $Q = 20\ \mu L/min$ with [1X] BHI. Fluorescence intensity signal was normalized to the maximum signal intensity value. Plots on the right show the corresponding evolution of the hydraulic resistance.

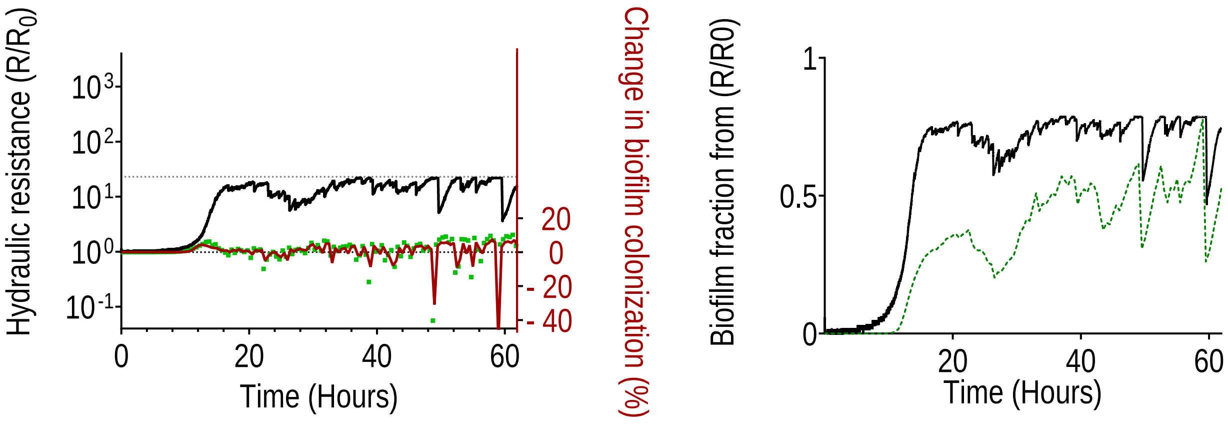


(a) (b)

Figure 14: Temporal dynamics for $Q = 200\ \mu L/min$. (a) evolution of the hydraulic resistance (black solid line) in time, calculated from pressure fluctuations. It also shows changes in biofilm colonization extracted from either integrated fluorescence intensity (green squares) or image segmentation (red solid line). (b): Fraction of biofilm in the microchannel, either calculated from hydraulic resistance (black solid line) or estimated from integrated GFP intensity (green dotted line), for the different flow rates.

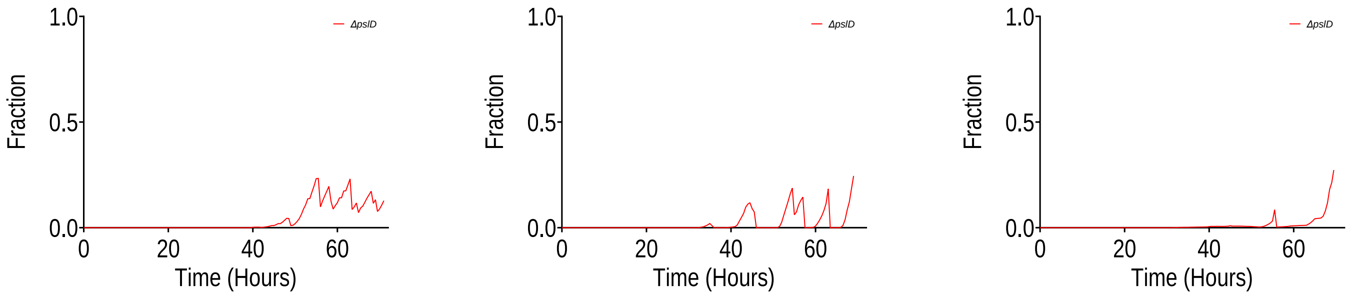


(a) n = 1 (b) n = 2 (c) n = 3

Figure 15: **Temporal dynamics of growth and detachment for** ***ΔpslD*** **strain with $Q = 0.2\ \mu L/min$.** (a), (b) and (c): volume fraction calculated from image segmentation for three replicates.

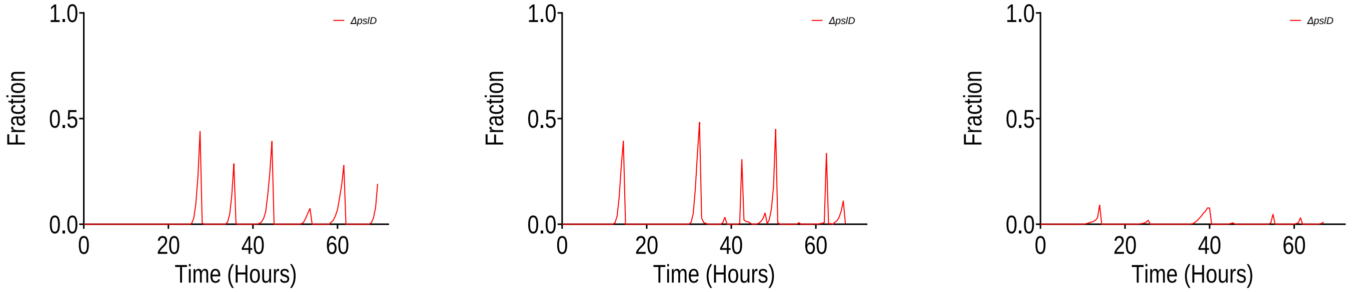


(a) n = 1 (b) n = 2 (c) n = 3

Figure 16: **Temporal dynamics of growth and detachment for** $\Delta pslD$ **strain with** $Q = 2\ \mu L/min$**.** (a), (b) and (c): volume fraction calculated from image segmentation for three replicates.

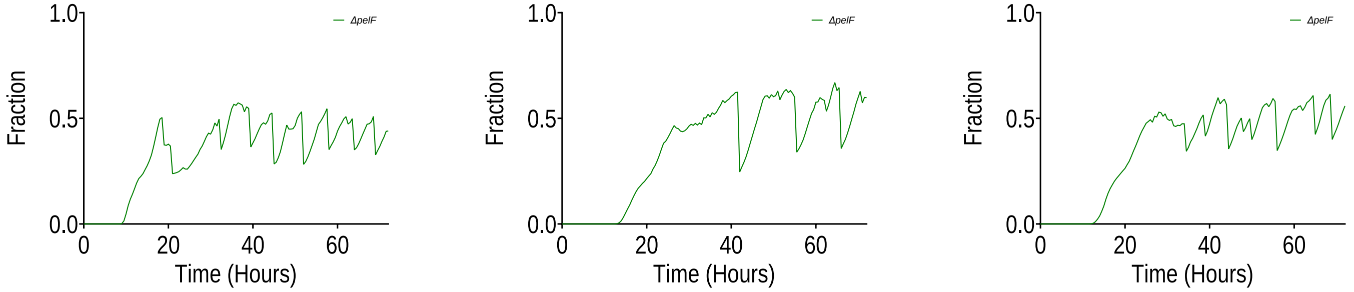


(a) n = 1 (b) n = 2 (c) n = 3

Figure 17: **Temporal dynamics of growth and detachment for** $\Delta pelF$ **strain with** $Q = 0.2\ \mu L/min$**.** (a), (b) and (c): volume fraction calculated from image segmentation for three replicates.

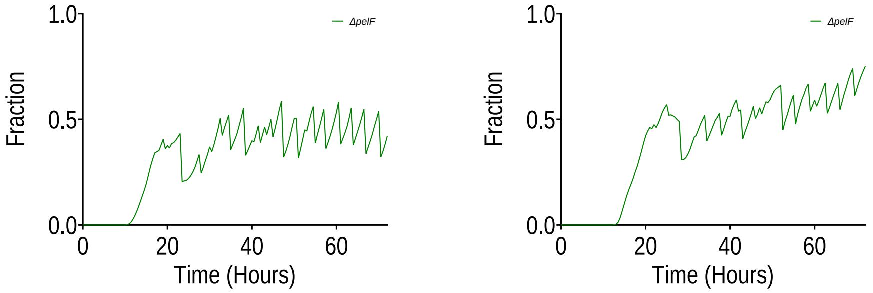


(a) n = 1 (b) n = 2

Figure 18: **Temporal dynamics of growth and detachment for** $\Delta pelF$ **strain with** $Q = 2\ \mu L/min$**.** (a), (b) and (c): volume fraction calculated from image segmentation for two replicates.

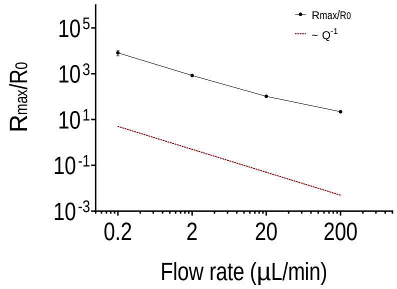


Figure 19: Log-log plot of $\frac{R_{\max}}{R_0}$ as a function of the flow rate ($Q = 0.2\ \mu L/min$, $2\ \mu L/min$, $20\ \mu L/min$ and $200\ \mu L/min$). The red dotted line simply shows the slope for an evolution with the inverse of the flow rate $Q^{-1}$. Error bars represent standard deviation for n = 3 replicates, except for $Q = 200\ \mu L/min$, for which n = 1.

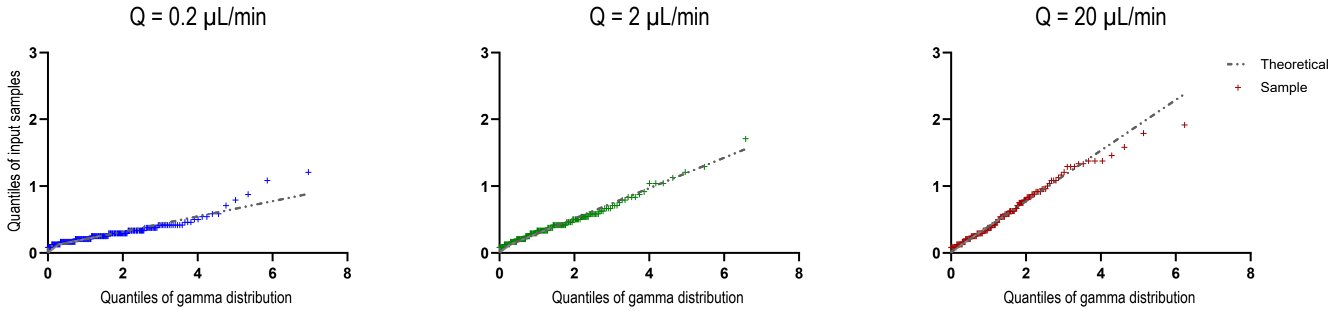


Figure 20: Quantile-quantile (Q-Q) plots of inter-event times fitted with Gamma distributions at three flow rates: $Q = 0.2\ \mu L/min$, $2\ \mu L/min$, $20\ \mu L/min$. Sample quantiles (y-axis) are plotted against theoretical Gamma quantiles (x-axis). Close alignment with the diagonal reference line (red) indicates that the Gamma distribution provides a reasonable approximation of the overall distributional behavior.

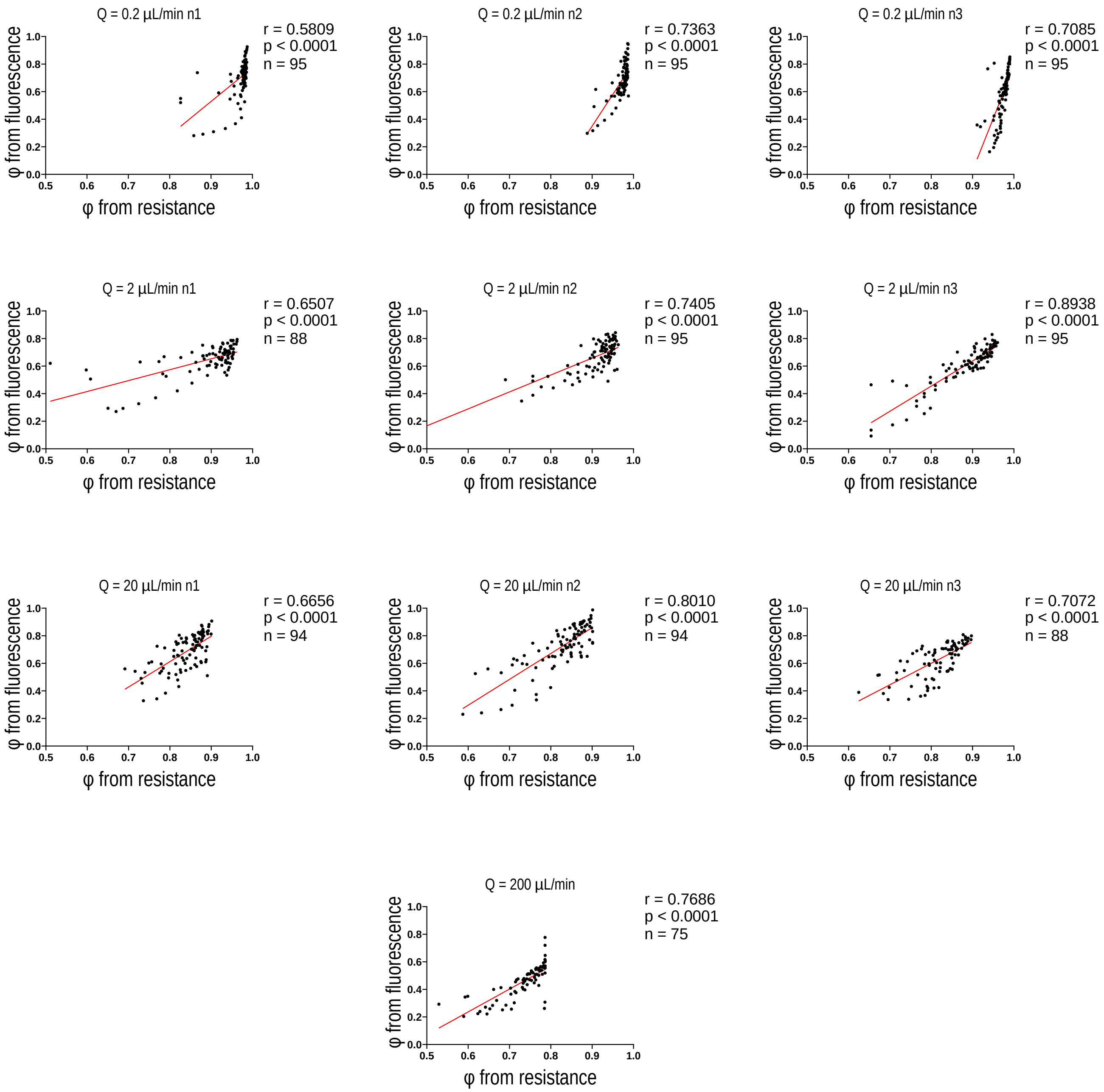


Figure 21: Pearson's correlation between volume fractions obtained from image segmentation and calculated from hydraulic resistance for flow rates $Q = 0.2\ \mu L/min$, $2\ \mu L/min$, $20\ \mu L/min$ and $200\ \mu L/min$. To account for acquisition frequency mismatch, hydraulic resistance data were downsampled from 1 Hz to 1 point per 30 minutes. Each point represents one paired measurement. Data from all replicates were pooled for each condition (sample size n indicated on each plot). Red lines show linear regression fits: r = 0.68, 0.76, 0.72, 0.77 (mean across the 3 replicates) for $Q = 0.2\ \mu L/min$, $2\ \mu L/min$, $20\ \mu L/min$, $200\ \mu L/min$ respectively; p $<$ 0.0001 for all conditions.

| Strain | WT PAO1 ATCC 15692:GFP | WT PAO1 pMRP9-1:GFP | $\Delta$*pel* PAO1 pMRP9-1 :GFP | $\Delta$*psl* PAO1 pMRP9-1 :GFP |
|---|---|---|---|---|
| Lag phase duration (minutes) | 233 +/- 3,00 | 275 +/- 8,33 | 424 +/- 16,30 | 377 +/- 40,17 |
| Doubling time (minutes) | 105 +/- 3,81 | 108 +/- 8,45 | 134 +/- 17,11 | 135 +/- 9,83 |

Table I: Lag phase and doubling time in minutes for *P. aeruginosa* WT PAO1 and EPS defective strains. Mean values were obtained from 3 replicates for each strain and +/- indicates the standard deviation.

#### III. SUPPLEMENTARY MOVIES

- Movie S1: DIC timelapse of the initial development at 0.2 $\mu L/min$.
- Movie S2: DIC timelapse of the initial development at 2 $\mu L/min$.
- Movie S3: DIC timelapse of the initial development at 20 $\mu L/min$.
- Movie S4: composite (GFP and brightfield) timelapse of biofilm development at 20 $\mu L/min$.
